# A nematode-specific gene underlies bleomycin-response variation in *Caenorhabditis elegans*

**DOI:** 10.1101/565218

**Authors:** Shannon C. Brady, Stefan Zdraljevic, Karol W. Bisaga, Robyn E. Tanny, Daniel E. Cook, Daehan Lee, Ye Wang, Erik C. Andersen

**Author notes:** **Corresponding author:** Erik C. Andersen, Department of Molecular Biosciences, Northwestern University, 4115 Pancoe NSUHS Life Sciences Pavilion, 2205 Tech Drive, Evanston, IL 60208.

## Abstract

Bleomycin is a powerful chemotherapeutic drug used to treat a variety of cancers. However, individual patients vary in their responses to bleomycin. The identification of genetic differences that underlie this response variation could improve treatment outcomes by tailoring bleomycin dosages to each patient. We used the model organism *Caenorhabditis elegans* to identify genetic determinants of bleomycin-response differences by performing linkage mapping on recombinants derived from a cross between the laboratory strain (N2) and a wild strain (CB4856). This approach identified a small genomic region on chromosome V that underlies bleomycin-response variation. Using near-isogenic lines and strains with CRISPR-Cas9 mediated deletions and allele replacements, we discovered that a novel nematode-specific gene (*scb-1*) is required for bleomycin resistance. Although the mechanism by which this gene causes variation in bleomycin responses is unknown, we suggest that a rare variant present in the CB4856 strain might cause differences in the potential stress-response function of *scb-1* between the N2 and CB4856 strains, thereby leading to differences in bleomycin resistance.

**Article summary:** We performed linkage mapping on a panel of recombinant lines generated between two genetically divergent strains of *Caenorhabditis elegans* and identified a bleomycin-response QTL. We generated CRISPR-Cas9 deletions and reciprocal allele-replacement strains for all six candidate genes across the QTL confidence interval. Deletions of one gene, *H19N07.3*, caused increased bleomycin sensitivity in both divergent genetic backgrounds. This gene might act in stress responses and detoxification in nematodes. We further compared our linkage mapping to a genome-wide association mapping and showed that a rare expression variant in the CB4856 strain likely underlies bleomycin-response differences.

## INTRODUCTION

Cancer is the second leading cause of death worldwide (World Health Organization, 2018), which has led to extensive research for treatments, including the identification of over 100 effective chemotherapeutic drugs (Santos *et al*. 2017). One of these drugs is bleomycin, an anti-tumor antibiotic that interacts with oxygen and transition metals to cause double-stranded DNA breaks (Chen and Stubbe 2005). Although the cytotoxicity of bleomycin can reliably induce cell death in tumor cells, off-target effects can lead to a range of harmful consequences from mild gastrointestinal irritation to severe bleomycin-induced pulmonary fibrosis (Blum *et al*. 1973). The tradeoff between efficacy and toxicity varies across individuals, and understanding the genetic variants that affect bleomycin response might yield opportunities to broaden the therapeutic range (Relling and Dervieux 2001; de Haas *et al*. 2008).

Bleomycin sensitivity has been shown to be heritable, suggesting that genetic markers can be used to predict bleomycin responses (Cloos *et al*. 1999). Many studies have attempted to identify the genetic variant(s) that underlie bleomycin-response differences across cancer patients, and some have identified potential connections between the metabolic enzyme bleomycin hydrolase (BLMH) and patient outcomes. However, none of these studies established a causal connection between genetic differences in BLMH and variation in bleomycin responses (Lazo and Humphreys 1983; Nuver *et al*. 2005; de Haas *et al*. 2008; Gu *et al*. 2011; Altés *et al*. 2013). The inability to identify a human genetic variant that causes differences in bleomycin responses might be attributed to limited sample sizes (Sham and Purcell 2014), confounding environmental factors (Hunter 2005; Liu *et al*. 2013), variation in drug regimens across patients (Low *et al*. 2013), or tumor complexity and progression (McClellan and King 2010; Dagogo-Jack and Shaw 2018). However, the DNA-damage pathways that might be implicated in bleomycin responses are evolutionarily conserved across eukaryotes (Taylor and Lehmann 1998). Therefore, studying bleomycin responses in a model organism with natural genetic variation can offer insights into how bleomycin response differs across individuals and can potentially be applied to the clinic (Zdraljevic and Andersen 2017).

*Caenorhabditis elegans* is a soil-associated microscopic roundworm that is an excellent model for basic cellular and organismal processes (A. K. Corsi, B. Wightman, M. A. Chalfie 2015). Not only does *C. elegans* have a well annotated reference genome (C. elegans Sequencing Consortium 1998; Stein *et al*. 2001; Hillier *et al*. 2005); www.wormbase.org WS268), but this species also has broad genomic diversity across global populations (Cook *et al*. 2017). Notably, the N2 strain and the CB4856 strain are well characterized and genetically divergent with approximately one single nucleotide variant per 850 bp (Wicks *et al*. 2001; Swan *et al*. 2002; Thompson *et al*. 2015). These two strains were used to generate a panel of recombinant inbred advanced intercross lines (RIAILs) (Rockman and Kruglyak 2009; Andersen *et al*. 2015), which has been used to correlate genetic variants with differences in quantitative traits (Kammenga *et al*. 2007; Seidel *et al*. 2008, 2011; Palopoli *et al*. 2008; Reddy *et al*. 2009; McGrath *et al*. 2009; Bendesky *et al*. 2011, 2012; Andersen *et al*. 2014; Schmid *et al*. 2015; Zdraljevic *et al*. 2017, 2019; Lee *et al*. 2017).

Here, we used a high-throughput fitness assay to measure bleomycin responses across a panel of 249 RIAILs (Andersen *et al*. 2015) and then performed linkage mapping to identify quantitative trait loci (QTL) that underlie bleomycin-response variation. We used near-isogenic lines (NILs) to validate the largest effect QTL on chromosome V. Our results from the NIL assays suggested that epistatic loci underlie bleomycin responses, but a two-factor genome scan was unable to detect significant genetic interactions. Next, we created and tested CRISPR-Cas9 mediated deletion alleles to investigate all six candidate genes in the QTL region. We identified a nematode-specific gene, *H19N07.3*, that underlies this QTL. Although this gene does not contain a protein-coding variant between the N2 and CB4856 strains, its gene expression varies across the RIAIL panel. Interestingly, a genome-wide association (GWA) approach identifies different QTL than the linkage mapping approach, suggesting that both common and rare variants underlie bleomycin response variation. Given the genetic complexity underlying the bleomycin response phenotype, this study highlights the power of the *C. elegans* model system to identify elusive causal genes.

## MATERIALS AND METHODS

### Strains

Animals were grown at 20° on 6 cm plates of modified nematode growth medium (NGMA), which contained 1% agar and 0.7% agarose, spotted with OP50 bacteria (Ghosh *et al*. 2012). The two parental strains used in this study were N2 and CB4856. N2 is the canonical laboratory strain of *C. elegans* that has been extensively studied (Brenner 1974). CB4856 is a well studied Hawaiian wild isolate that is genetically divergent from N2 and has a characterized genome (Wicks *et al*. 2001; Swan *et al*. 2002; Thompson *et al*. 2015). The N2 and CB4856 strains were crossed for several generations to create a panel of recombinant inbred advanced intercross lines (RIAILs) that contain regions of the genome derived from each parental strain. These RIAILs were constructed previously (Rockman and Kruglyak 2009; Andersen *et al*. 2015) and have well characterized genotypes and allele frequencies, and we used this panel of RIAILs in our study to identify regions of the genome correlated with drug response. The construction of near-isogenic lines (NILs), as well as CRISPR-Cas9 mediated deletion and allele-replacement strains, is detailed below. All strains and reagents used in strain constructions are listed in the **Supplementary Information**.

### High-throughput fitness assays

We used the high-throughput assay (HTA) described previously (Evans *et al*. 2018) and following is a summary of that assay (**Figure S1**). Populations of each strain were passaged on 6 cm plates for four generations to amplify animal numbers and reduce the effects of starvation (Andersen *et al*. 2014). Gravid adults were bleached for stage synchronization, and approximately 25 embryos from each strain were aliquoted into 96-well plates at a final volume of 50 µL of K medium (Boyd *et al*. 2012). The following day, arrested L1 larvae were fed 5 mg/mL HB101 bacterial lysate in K medium (Pennsylvania State University Shared Fermentation Facility, State College, PA; (García-González *et al*. 2017) and were grown for 48 hours at 20° with constant shaking. A large-particle flow cytometer (COPAS BIOSORT, Union Biometrica, Holliston, MA) was used to sort three L4 larvae into each well of a 96-well plate that contained 50 µL K medium plus HB101 lysate at 10 mg/mL, 50 µM kanamycin, and either 1% distilled water (control) or 1% distilled water and bleomycin (drug). The sorted L4 larvae were grown and propagated for 96 hours at 20° with constant shaking. The population of parents and progeny were treated with sodium azide (50 mM in M9) and quantified by the BIOSORT for several fitness parameters. Because bleomycin exposure can affect animal proliferation (brood size), animal growth (length), and animal development (optical density), the fitness parameters we measured with the BIOSORT included brood size (n), animal length (time of flight, TOF), and optical density (extinction time, EXT).

### Bleomycin-response trait measurements and processing

Phenotypic measurements collected by the BIOSORT were processed using the R package *easysorter* (Shimko and Andersen 2014). Using this package, *read_data* imported measurements from the BIOSORT and *remove_contamination* was used to remove contaminated wells from analysis. The *sumplate* function then calculated normalized measurements (norm.n -- brood size normalized to number of animals sorted, norm.EXT -- EXT normalized by TOF measurements) and summary statistics (mean, median, 10^th,^ 25^th^, 75^th^, 90^th^ percentile, interquartile range, covariance, and variance) of each trait for the population of animals. A total of 26 HTA traits were measured. When strains were phenotyped across multiple days, the *regress(assay=TRUE)* function was used to fit a linear model with the phenotype as the dependent variable and assay as the independent variable (*phenotype ∼ assay*) to account for variation among assay days. Next, the *prune_outliers()* function removed phenotypic values that were beyond two standard deviations of the mean (unless at least 5% of the strains were outside this range in the case of RIAIL assays). Finally, bleomycin-specific effects were calculated using the *regress(assay=FALSE)* function from *easysorter*, which fits a linear model with the phenotype as the dependent variable and control phenotype as the independent variable (*phenotype ∼ control phenotype*). The residual phenotypic values account for differences among strains that were present in control conditions.

### Bleomycin dose response

A dose-response high-throughput assay was performed using quadruplicates of four genetically divergent strains (N2, CB4856, JU258, and DL238) tested in various concentrations of bleomycin (**File S1**). The broad-sense heritability at each concentration was calculated using the *lmer* function within the *lme4* R package with the phenotype as the dependent variable and strain as a random effect *phenotype ∼ 1 + (1|strain)* (**File S2**). The concentration of bleomycin that provided the highest mean broad-sense heritability across the 26 HTA traits was selected for linkage mapping experiments (50 µM, mean *H*^2^ = 0.58). Bleomycin sulfate was purchased from Biotang Inc via Fisher Scientific (Catalog No. 50-148-546).

### Whole-genome sequence library prep and analysis

Whole-genome sequencing was performed on recombinant advanced intercross lines (RIAILs) and near-isogenic lines (NILs) using low-coverage sequencing. DNA was isolated from 100-300 µL of packed worms using Omega BioTek’s EZ 96 Tissue DNA Kit (catalog no. D1196-01). All samples were diluted to 0.2 ng/µL and incubated with diluted Illumina transposome (catalog no. FC-121-1031). Tagmented samples were amplified with barcoded primers. Unique libraries (192) were pooled by adding 8 µL of each library. The pooled material was size-selected by separating the material on a 2% agarose gel and excising the fragments ranging from 400-600 bp. The sample was purified using Qiagen’s Gel Extraction Kit (catalog no. 28706) and eluted in 30 µL of buffer EB. The concentration of the purified sample was determined using the Qubit dsDNA HS Assay Kit (catalog no. Q32851). RIAILs and NILs were sequenced at low coverage (mean = 2.13x) using the Illumina HiSeq 2500 platform with a paired-end 100 bp reaction lane. RIAIL and NIL genotypes were imputed using VCF-kit (Cook and Andersen 2017). To determine genotypes, a list of filtered, high-quality sites (n = 196,565) where parental strains possess different genotypes was extracted from a previously established variant dataset (Cook *et al*. 2016). All RIAIL genotypes can be accessed by downloading the *linkagemapping* R package at github.com/AndersenLab/linkagemapping and by running *load_cross_obj(“N2xCB4856cross_full”)*. NIL genotypes are described below and are available in **File S5** and **File S6**.

### Linkage mapping

Linkage mapping was performed on each of the 26 HTA traits described above using the R package *linkagemapping* (www.github.com/AndersenLab/linkagemapping, **Figure S4, Figure S5, File S4**). The function *load_cross_obj*(“N2xCB4856cross_full”) was executed to load a cross object containing 13,003 SNPs that describe locations of genetic recombination in the RIAIL panel (out of the 195,565 high-quality SNPs at which genotypes were called). The genotypic data and residual phenotypic data (after control-condition regression) were merged into a cross object using the *merge_pheno* function with *set* = 2 to include strains with a reduced allele-frequency skew on chromosome I. The *fsearch* function was used to calculate logarithm of odds (LOD) scores for each marker and each trait as -n(ln(1 - R^2^)/2ln(10)), where *R* is the Pearson correlation coefficient between RIAIL genotypes at the marker and trait values (Bloom *et al*. 2013; Zdraljevic *et al*. 2017; Lee *et al*. 2017; Evans *et al*. 2018). A 5% genome-wide error rate was calculated by permuting phenotype and genotype data 1,000 times. Mappings were repeated iteratively, each time using the marker with the highest LOD score as a cofactor until no significant QTL were detected. The *annotate_lods* function was used to calculate the percent of variance in RIAIL phenotypes explained by each QTL and a 95% confidence interval around each peak marker, defined by any marker on the same chromosome within a 1.5-LOD drop from the peak marker.

### Generation of near-isogenic lines (NILs)

A near-isogenic line (NIL) is genetically identical to another strain aside from a small genomic region that is derived from an alternate strain. NILs are used to test the effect of modifications to particular genomic regions in a consistent genetic background. To make each NIL, males and hermaphrodites of the desired RIAIL (QX131 for ECA230 and QX450 for ECA232) and parental background (CB4856 for ECA230 and N2 for ECA232) were crossed in bulk, then male progeny were crossed to the parental strain in bulk for another generation. For each NIL, eight single-parent crosses were performed followed by six generations of propagating isogenic lines to ensure homozygosity of the genome. For each cross, PCR was used to select non-recombinant progeny genotypes within the introgressed region by amplifying insertion-deletion (indel) variants between the N2 and CB4856 genotypes on the left and right side of the introgressed region. NIL strains ECA411 and ECA528 were generated by crossing ECA230 and CB4856 in bulk. Heterozygous F1 hermaphrodites were crossed to CB4856 males and the F2 L4 hermaphrodites were placed into individual wells of a 96-well plate with K medium and 5 mg/mL bacterial lysate and grown to starvation. After starvation, each well of the 96-well plate was genotyped to identify recombinants in the desired region. Recombinant strains were plated onto 6 cm plates and individual hermaphrodites were propagated for several generations to ensure homozygosity across the genome. NIL strains were whole-genome sequenced as described above to confirm their genotypes (**File S5, File S6**). Reagents used to generate all NIL strains and a summary of each introgressed region are detailed in the **Supplementary Information**.

### Two-factor genome scan

We performed a two-factor genome scan to identify potentially epistatic loci that might contribute to traits of interest (either bleomycin responses or gene-expression levels). We used the *scantwo* function in the R *qtl* package to perform this analysis. Each pairwise combination of loci was tested for correlations with trait variation in the RIAILs. The summary of each scantwo output includes the maximum interactive LOD score for each pair of chromosomes. To determine a threshold for significant interactions, we performed 1000 permutations of the scantwo analysis and selected the 95^th^ percentile from these permutations. For the bleomycin-response variation scantwo, the significant interaction threshold was a LOD score of 4.08. The significant interaction threshold for the *scb-1* expression variation scantwo was 4.05. The bleomycin-response variation scantwo summary is available as **File S8**, and the scantwo summary for *scb-1* expression variation is available as **File S17**.

### Identification of genes with non-synonymous variants

The *get_db* function within the *cegwas* R package was used to query WormBase build WS245 for genes in the QTL confidence interval (V:11,042,745-11,189,364). Our initial linkage mappings used the 1,454 marker set (Rockman and Kruglyak 2009) and had a QTL confidence interval larger than the interval found using the whole-genome marker set (described above). Because this confidence interval was larger and more conservative, we kept it for subsequent testing of candidate genes. This expanded interval contained an additional 20 kb on either side of the whole-genome marker set confidence interval (**File S9**). The *snpeff* function within the *cegwas* R package was used to identify variants within the region of interest (**File S11**). We identified variants predicted to have MODERATE (coding variant, inframe insertion/deletion, missense variant, regulatory region ablation, or splice region variant) or HIGH (chromosome number variant, exon loss, frameshift variant, rare amino acid variant, splice donor/acceptor variant, start loss, stop loss/gain, or transcript ablation) phenotypic effects according to Sequence Ontology (Eilbeck *et al*. 2005) and selected variants at which the CB4856 strain contains the alternate allele. This search found five candidate genes in the interval: *C45B11.8*, *C45B11.6*, *jmjd-5*, *srg-42*, and *cnc-10*, which each contain one non-synonymous variant between the N2 and CB4856 strains.

### Generation of deletion strains

Deletion alleles for genes of interest were generated to test the effects of loss-of-function variants on bleomycin responses. For each desired deletion, we designed a 5’ and a 3’ guide RNA with the highest possible on-target and off-target scores, calculated using the Doench algorithm (Doench *et al*. 2016). The N2 and CB4856 L4 hermaphrodite larvae were picked to 6 cm agar plates seeded with OP50 *E. coli* 24 hours before injection. The CRISPR injection mix was assembled by first incubating 0.88 µL of 200 µM AltR® CRISPR-Cas9 tracrRNA (IDT, catalog no. 1072532), 0.82 µL of each of the 5’ and 3’ AltR® CRISPR-Cas9 crRNAs at a stock concentration of 100 µM in Duplex Buffer (IDT), and 0.12 µL of 100 µM *dpy-10* co-injection crRNA at 95° for five minutes. 2.52 µL of 69 µM AltR® *S. pyogenes* Cas9 Nuclease (IDT, catalog no. 1081058) was added to the tracrRNA/crRNA complex mixture and incubated at room temperature for five minutes. Finally, 0.5 µL of 10 µM *dpy-10* repair construct and distilled water were added to a final volume of 10 µL. The injection mixture was loaded into the injection capillary using a mouth pipet to avoid bubbles in the injection solution. Young adult animals were mounted onto agar injection pads, injected in either the anterior or posterior arm of the gonad, and allowed to recover on 6 cm plates in bulk. Twelve hours after injection, survivors were transferred to individual 6 cm plates. Broods with successful edits were easily observed because of the *dpy-10* co-injection marker, which created animals with an obvious locomotive defect, roller (Rol), or morphological phenotype, dumpy (Dpy). Three to four days after injections, plates were checked for the presence of Rol or Dpy F1 progeny. These Rol F1 progeny were transferred to individual 6 cm plates, allowed to lay offspring, and genotyped for the desired edit 24 hours later. Genotyping was performed with PCR amplicons designed around the desired site of the deletion. Plates with heterozygous or homozygous deletions were propagated and genotyped for at least two more generations to ensure homozygosity and to cross out the Rol mutation. Deletion amplicons were Sanger sequenced to identify the exact location of the deletion. Reagents used to generate deletion alleles are detailed in **Supplemental Information.**

### Generation of CRISPR-mediated *jmjd-5* allele replacements

Allele replacement strains were created to test the effect of a particular amino acid substitution on bleomycin responses. A guide RNA was designed to cut two bp downstream of the natural variant, with an on-target score of 31 and off-target score of 47 (Doench *et al*. 2016). Two repair constructs were designed, one for the N2 to CB4856 replacement and *vice versa*. Repair oligonucleotides were homologous to the background strain except for the nucleotide variant, a silent mutation in the PAM site (A339A) to eliminate repair construct cleavage, and a silent mutation that introduces a *BsaA*I restriction enzyme cut site (T336T). Repair constructs contained a 35-bp homology arm on the PAM-proximal side of the edit and a 91-bp homology arm on the PAM-distal side of the edit. Injection mixes were made as above, with 0.6 µL of 100 µM *jmjd-5* repair construct added with the *dpy-10* repair construct in the last step of the protocol. Animals were injected as above and Rol F1 progeny were genotyped using PCR and restriction enzyme digestion. As with the deletion alleles, edited strains were homozygosed and their genotypes were confirmed with Sanger sequencing. Reagents used to generate these allele replacement strains are detailed in the **Supplemental Information**.

### Linkage mapping of expression QTL

Microarray data for 13,107 probes were collected for synchronized young adult populations of 209 recombinant lines previously (Rockman *et al*. 2010). We performed linkage mapping as described above on these expression data and identified significant peaks for 3,298 probes (**File S13**). For the *H19N07.3* probe (A_12_P104350), the full annotated-LOD data frame is provided (**File S14**).

### Hemizygosity high-throughput assay

Hemizygosity assays were used to test the effect of zero (two deletion alleles), one (one deletion allele and one wild-type allele), or two (two wild-type alleles) functional copies of a gene product on bleomycin responses. If the phenotype is affected by gene function, one would expect to see bleomycin responses scale with the number of functional alleles of the gene present in each strain. For each heterozygous/hemizygous genotype, 30 hermaphrodites and 60 males were placed onto each of four 6 cm plates and allowed to mate for 48 hours. The same process was completed for homozygous strains to remove biases introduced by the presence of male progeny in the assay. Mated hermaphrodites were transferred to a clean 6 cm plate and allowed to lay embryos for eight hours. After the egg lay period, adults were manually removed from egg lay plates, and embryos were washed into 15 mL conicals using M9 and a combination of pasteur pipette rinsing and scraping with plastic inoculation loops. Embryos were rinsed and centrifuged four times with M9 before being resuspended in K medium and 50 µM kanamycin. Embryos hatched and arrested in the L1 larval stage in conicals overnight at 20° with shaking. The next morning, 50 L1 larvae were sorted into each well of a 96-well plate containing K medium, 10 mg/mL bacterial lysate, 50 µM kanamycin, and either 1% distilled water or 1% distilled water plus bleomycin using the BIOSORT (dose-response assay, **Figure S10, File S1**) or by titering (hemizygosity assays, **Figure 6, Figure S11, Figure S12, File S1**). Animals were incubated in these plates for 48 hours at 20° with shaking and were paralyzed with 50 µM sodium azide in M9 before being scored for phenotypic parameters using the BIOSORT.

### Statistical analysis

All statistical tests of phenotypic differences in the NIL, deletion strain, and allele-replacement strain assays were performed in R (version 3.3.1) using the *TukeyHSD* function (R Core Team) and an ANOVA model with phenotype as the dependent variable and strain as the independent variable (*phenotype ∼ strain*). The p-values of individual pairwise strain comparisons were reported, and p-values less than 0.05 were deemed significant. For each figure (with the exception of hemizygosity tests), phenotypes of NILs, deletion strains, and allele-replacement strains were compared to phenotypes of their respective background strains (either N2 or CB4856), and statistical significance is denoted by an asterisk above the boxplot for each strain. P-values of all pairwise comparisons are reported in **File S7**. Correlation between RIAIL *H19N07.3* expression and median optical density in bleomycin was tested using a Spearman’s correlation test. Statistical difference between N2 and CB4856 expression of *H19N07.3* measured by RNA-seq was tested using a likelihood-ratio and a Wald test with a threshold of *p* < 0.05.

### RNA-seq

Three independent replicates of RNA were sampled as follows. Bleach-synchronized embryos (∼2,000) of the N2 and CB4856 strains were grown on 10 cm plates of NGMA for 66-69 hours. When F1 embryos appeared on the plate, synchronized young adults were collected by washing each plate twice with M9 buffer and incubating for 30 minutes in M9 to remove remaining bacteria. Then, samples were washed twice again with M9 buffer, and then washed with sterile water. Animals were pelleted and homogenized in Trizol (Ambion) by repeating freezing-thawing with liquid nitrogen five times. To extract RNA from each sample, chloroform, isopropanol, and ethanol were used for phase separation, precipitation and washing steps, respectively. RNA pellets were resuspended in TE buffer, and RNA quality was measured with a 2100 Bioanalyzer (Agilent). Library preparation and RNA sequencing (HiSeq4000, Illumina) were performed by the Genomics Facility at the University of Chicago. RNA-seq data were quantified with kallisto and then within-sample and between-sample normalization was performed using sleuth, which is based on DESeq (Anders and Huber 2010; Bray *et al*. 2016; Pimentel *et al*. 2017). Significant differences between samples were determined by likelihood-ratio and Wald tests. RNA-seq data are reported in **File S16**.

### Genome-wide association mapping

Bleomycin responses were measured for 83 *C. elegans* isotypes using the high-throughput fitness assay (**File S18**). Genome-wide association mapping was performed as described previously (Zdraljevic *et al*. 2019) using genotype data from CeNDR Release 20180527 (Cook *et al*. 2017). In short, BCFtools was used to remove variants with missing genotype calls and variants with a minor allele frequency below 5% (Li 2011), and PLINK v1.9 was used to prune the genotypes at a linkage disequilibrium threshold of r^2^ < 0.8 (Purcell *et al*. 2007; Chang *et al*. 2015), for a total of 59,241 pruned markers. A kinship matrix was generated using the *A.mat* function in the *rrBLUP* R package (Endelman 2011; Endelman and Jannink 2012). The *GWAS* function in the *rrBLUP* package was used to perform genome-wide association mapping with EMMA algorithm to correct for kinship (Kang *et al*. 2008; Covarrubias-Pazaran 2016). The relatedness among these wild isolates was described previously (Andersen *et al*. 2012; Hahnel *et al*. 2018; Zdraljevic *et al*. 2019).

### Identification of rare variants

VCF release 20180527 was downloaded from elegansvariation.org (Cook *et al*. 2017). The VCF was filtered to select all variants within the linkage mapping confidence interval (V:11042745-11189364) where CB4856 contains the alternate allele (**File S20**). Variants with a minor allele frequency less than 0.05 within the 83 wild isolates that have a bleomycin median optical density measurement were deemed “rare”.

### Creation of neighbor-joining tree

Protein sequences for homologs of the *C. elegans* H19N07.3 protein (**File S21**) were input to MUSCLE (Edgar 2004a; b) to generate a multiple-sequence alignment. CLUSTALW was then used to generate a neighbor-joining tree and output as a Newick formatted file (**File S22**).

### Data availability

All data are available on Figshare. **File S1** contains all pruned data from the high-throughput bleomycin assays. **File S2** contains the broad-sense heritability estimates calculated for each drug concentration for all 26 HTA traits for the HTA dose response as well as for the 24 HTA traits in the modified HTA dose response. **File S3** contains all control-regressed data for the 26 HTA traits for all assays. **File S4** contains the annotated linkage mapping data for the 26 control-regressed HTA traits. **File S5** is a VCF that reports the genotype from whole-genome sequence for all NILs in the manuscript. **File S6** is a simplified version of **File S5** that contains information on recombination locations for all NILs and can be used for more user-friendly visualization of NIL genotypes. **File S7** contains all statistical information for HTA phenotypic differences reported in the manuscript. **File S8** is a summary of the scantwo analysis for bleomycin responses in the RIAILs and reports the maximum interaction LOD score for each chromosome pair. **File S9** contains information on all genes in the QTL confidence interval plus 20 kb on either side. **File S10** contains locations of the exons, introns, and transcription start and stop sites for all genes in the region. **File S11** reports predicted non-synonymous variants between the N2 and CB4856 strains in the region. **File S12** is derived from the Rockman *et al*. 2010 RIAIL microarray expression data, and reports the expression measurements for each of the 13,107 microarray probes across 209 RIAILs. **File S13** contains all significant QTL identified by linkage mapping of **File S12** data. **File S14** contains the annotated linkage mapping of the *H19N07.3* expression data. **File S15** reports the *H19N07.3* expression and residual median optical density for strains of the RIAIL panel that were assayed for both of those traits. **File S16** contains *H19N07.3* RNA-seq expression data for populations of young adults of N2 and CB4856. **File S17** is a summary of the scantwo analysis for *H19N07.3* expression in the RIAILs and reports the maximum interaction LOD score for each chromosome pair. **File S18** contains control-regressed phenotypic data for all wild isolates assayed in response to bleomycin. **File S19** contains genome-wide association mapping for the phenotypes in **File S18**. **File S20** contains genotype information for each strain measured in **File S18** across all variants within the linkage mapping confidence interval around the QTL for which CB4856 contains the alternate allele. **File S21** is a FASTA file containing the protein sequences for all *H19N07.3* homologs. **File S22** is a neighbor-joining tree derived from a multiple sequence alignment of all sequences from **File S21** in Newick tree format.

## RESULTS

### Genetic differences underlie bleomycin-response variation

Bleomycin causes double-stranded DNA breaks, which ultimately lead to cytotoxicity of rapidly dividing cell populations. Therefore, exposure to bleomycin can affect the development of *C. elegans* larvae as well as germ-cell production of adult animals. We used a high-throughput assay (HTA) to measure the effects of bleomycin on development and brood size (**Figure S1,** Materials and Methods). To determine the concentration of bleomycin that would maximize among-strain while minimizing within-strain phenotypic variation, we used the HTA to perform a dose-response assay. We assessed bleomycin responses for four divergent strains (N2, CB4856, JU258, and DL238) across each of 26 HTA traits (**File S1, Figure S2**). For each concentration of bleomycin, we calculated the broad-sense heritability of the traits (Materials and Methods) and found that heritability was maximized at 50 µM bleomycin (mean *H*^2^ across all traits = 0.58, **File S2**). Given these results, we exposed animals to 50 µM bleomycin for all future HTA experiments.

Two of the strains used in the dose response assay, N2 and CB4856, have been extensively characterized at the genome level (Wicks *et al*. 2001; Swan *et al*. 2002; Thompson *et al*. 2015) and displayed divergent bleomycin responses (**Figure S2**). Recombinant inbred advanced intercross lines (RIAILs) were previously constructed between these two strains (Rockman and Kruglyak 2009; Andersen *et al*. 2015), and these RIAILs have been leveraged to identify genetic variants that cause phenotypic differences between the N2 and CB4856 strains (Kammenga *et al*. 2007; Seidel *et al*. 2008, 2011; Palopoli *et al*. 2008; Reddy *et al*. 2009; McGrath *et al*. 2009; Bendesky *et al*. 2011, 2012; Andersen *et al*. 2014; Schmid *et al*. 2015; Zdraljevic *et al*. 2017; Lee *et al*. 2017). We used these RIAILs to identify genetic variants that contribute to differential bleomycin responses between the N2 and CB4856 strains. Using our HTA, we measured each of the 26 fitness parameters for 249 RIAILs (Materials and Methods, **File S3**). Correlations between each pairwise combination of the 26 HTA measurements revealed several clusters of highly correlated traits (**Figure S3**). Therefore, the summary statistics measured by the BIOSORT should not be considered independent traits for linkage mapping. We selected median optical density (median.EXT) for future analyses, which is related to both animal length and optical extinction, because this trait was highly correlated with many of the 26 HTA traits and was highly heritable (*H^2^* = 0.73, **File S2**).

### The QTL on the center of chromosome V strongly impacts bleomycin response

We performed linkage mapping on the residual median optical density measurements in bleomycin (**Figure S4, Figure S5**, **File S4,** Materials and Methods) and identified four significant quantitative trait loci (QTL, **Figure 1A**). The QTL on the center of chromosome V was highly significant (explained 43.58% of the total variation and 55.60% of the genetic variation) with a LOD score of 32.57, and it was detected for 25 of the 26 HTA traits (**Figure S5, File S4**). The QTL 95% confidence interval was approximately 147 kb. Strains with the N2 allele at the peak marker had a lower median optical density in bleomycin and were interpreted to be more sensitive than those RIAILs with the CB4856 allele (**Figure 1B**).

**Figure 1:**
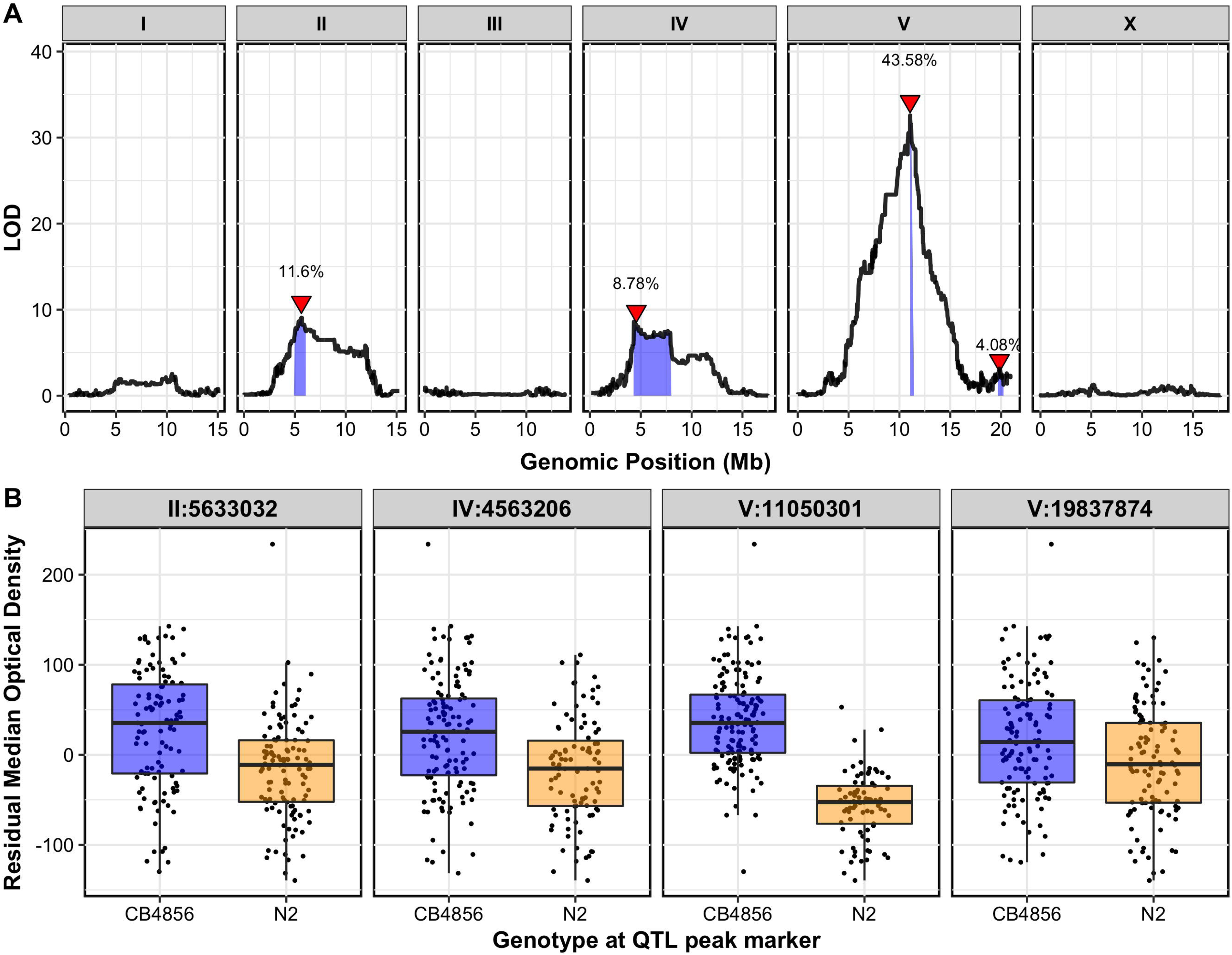
Linkage-mapping analysis of bleomycin-response variation is shown for residual median optical density in bleomycin. **A.** On the x-axis, each of 13,001 genomic markers, split by chromosome, were tested for correlation with phenotypic variation across the RIAIL panel. The log of the odds (LOD) score for each marker is reported on the y-axis. Each significant quantitative trait locus (QTL) is indicated by a red triangle at the peak marker, and a blue ribbon shows the 95% confidence interval around the peak marker. The total amount of phenotypic variance across the RIAIL panel explained by the genotype at each peak marker is shown as a percentage. **B.** Residual median optical density phenotypes (y-axis), split by allele at each QTL peak marker (x-axis), are shown. For each significant QTL, phenotypes of RIAILs that contain the N2 allele (orange) are compared to RIAILs that contain the CB4856 allele (blue). Phenotypes are shown as Tukey box plots with the phenotypes of each individual strain shown as points behind the box plots.

We isolated this QTL in a controlled genetic background by generating near-isogenic lines (NILs, **File S5**, **File S6**) that each contain a genetic background derived from either the N2 or CB4856 strain and a region of chromosome V from the opposite parental genotype. We used the HTA to test these strains in response to bleomycin (**File S3**). The NIL with the N2 genotype across this QTL introgressed into the CB4856 background (ECA230) was statistically more sensitive to bleomycin than CB4856 (**Figure S6, File S7**, *p* = 1.3e-14, Tukey HSD). This phenotype indicated that the N2 genotype within the introgressed region (which includes the QTL confidence interval) confers sensitivity to bleomycin. However, the reciprocal NIL with the CB4856 locus introgressed into the N2 background (ECA232) had a bleomycin-response phenotype that was not significantly different from the N2 strain (**Figure S6, File S7,** *p* = 0.053, Tukey HSD), suggesting that interacting loci could underlie bleomycin responses in a background-dependent manner. We performed a two-factor genome scan to map potential epistatic loci but did not identify a significant interaction between the QTL on chromosome V and other loci (**Figure S7, File S8**). However, the failure to detect significant interacting QTL could be because we have too few recombinant strains or because too few replicates of each RIAIL were phenotyped. Alternatively, more than two loci might underlie the transgressive phenotype of ECA230 and a two-factor genome scan might not be able to capture this complexity.

Nonetheless, because ECA230 recapitulated the expected QTL phenotype, we generated two NILs (ECA411 and ECA528) that narrowed this introgressed region to more precisely locate the causal variant (**File S5, File S6**). In addition, the N2 region on the left of chromosome V was removed from both NIL strains to ensure that this region of introgression did not underlie the phenotypic difference between ECA230 and CB4856. The genotypes of ECA411 and ECA528 differ in a small region of chromosome V that includes the QTL confidence interval (**Figure 2, File S5, File S6**). Both of these strains were more sensitive to bleomycin than the background parental strain, CB4856. This result could suggest that the introgressed region shared between these strains, which does not include the QTL, conferred some bleomycin-response variation between the N2 and CB4856 strains (**Figure 2**). Alternatively, the hypersensitivity of these NILs could suggest the presence of Dobzhansky-Muller incompatibilities between the N2 and CB4856 genotypes (Snoek *et al*. 2014) that might affect stress responses of the NILs. However, ECA528 was much more sensitive to bleomycin than ECA411 (**Figure 2, File S7**). Because ECA528 has the N2 genotype across the QTL region and ECA411 has the CB4856 genotype, these results suggest that the QTL genotype strongly affects bleomycin sensitivity (**Figure 2**, ECA528 vs. each other strain *p* < 1e-14, Tukey HSD). The empirically defined region lies between 10,339,727 and 11,345,443 bp on chromosome V and fully encompasses the linkage mapping confidence interval (from 11,042,745 to 11,189,364 bp on chromosome V).

**Figure 2:**
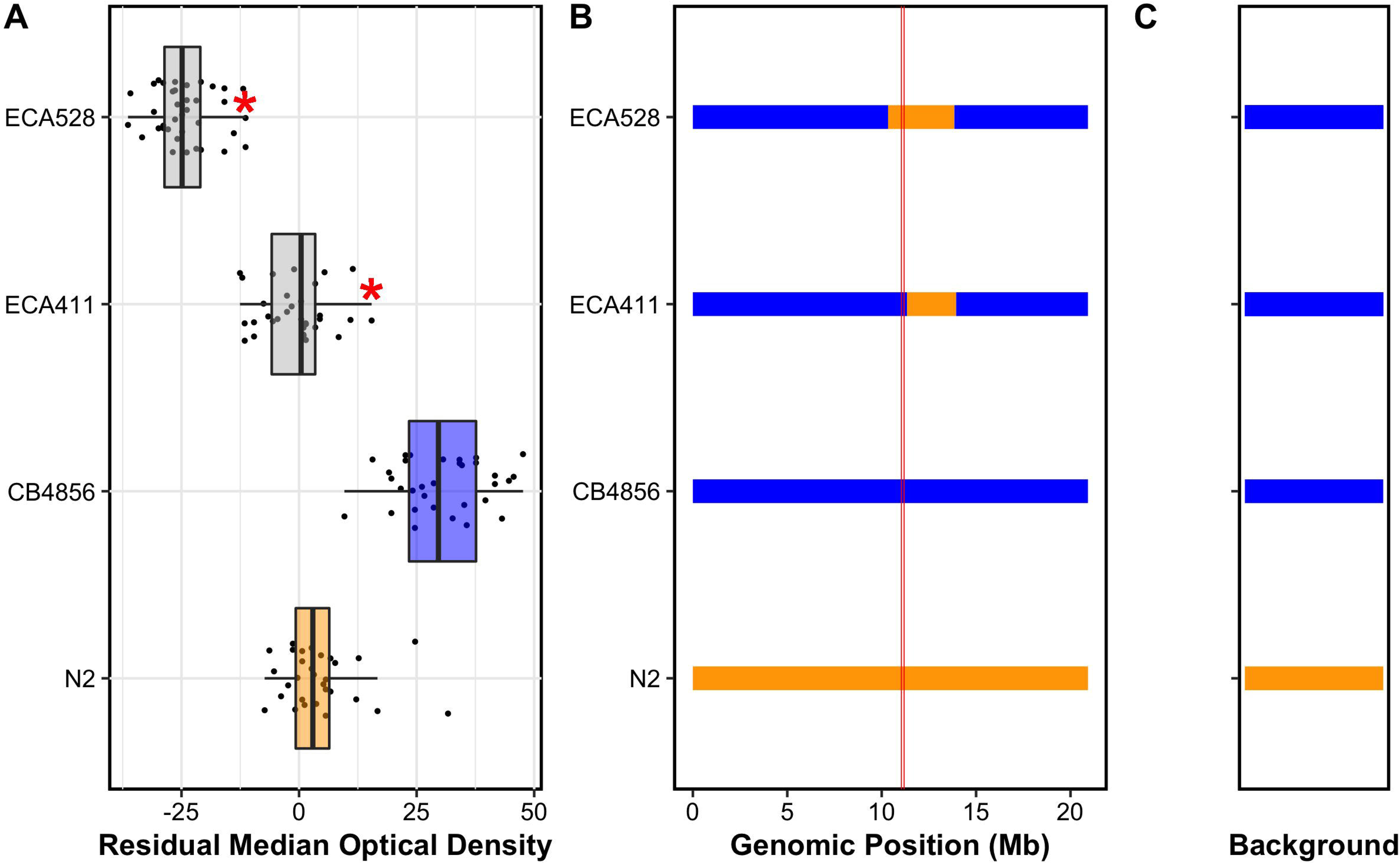
Phenotypes and genotypes of near-isogenic lines (NILs) are shown. (**A**) Phenotypes for each strain are shown as Tukey box plots, with strain name on the y-axis and residual bleomycin median optical density on the x-axis. Each point is a biological replicate. Parental strain box plots are colored by their genetic background, with orange indicating an N2 background and blue indicating a CB4856 genetic background. NILs are shown as grey box plots. A red asterisk indicates a significant difference between the phenotype of a given strain and the phenotype of the corresponding parental strain (*p* < 0.05, Tukey HSD). (**B**) Chromosomal representations of chromosome V are shown for each of the strains in **A**. Strain names are reported on the y-axis, and genomic position (Mb) is shown on the x-axis. Blocks of color indicate genotypes of genomic regions with orange indicating the N2 genotype and blue indicating the CB4856 genotype. Vertical red lines mark the confidence interval of the QTL from linkage mapping. (**C**) Background genotypes are represented as rectangles with colors indicating N2 (orange) or CB4856 (blue) genetic backgrounds.

### Genes with non-synonymous variants in the QTL region do not impact bleomycin responses

Because the recombination rate in the centers of *C. elegans* chromosomes is lower than chromosome arms (Rockman and Kruglyak 2009), it was difficult to generate additional NILs to narrow the QTL region further. Therefore, we took a targeted approach and created CRISPR-Cas9 directed modifications of candidate genes in the region. The 147 kb confidence interval on chromosome V contains 93 genes, including pseudogenes, piRNA, miRNA, ncRNA, and protein-coding genes (**File S9**). Given the narrow confidence interval, we expanded our search to include an additional 20 kb on each side of the 147 kb interval (Materials and Methods). Of the 118 genes included in the wider region, five genes, *C45B11.8*, *C45B11.6*, *jmjd-5*, *srg-42*, and *cnc-10*, contain predicted non-synonymous variants between the N2 and CB4856 strains (**Figure 3A**, **File S10**, **File S11**). These variants could cause differential bleomycin sensitivity between the N2 and CB4856 strains.

**Figure 3:**
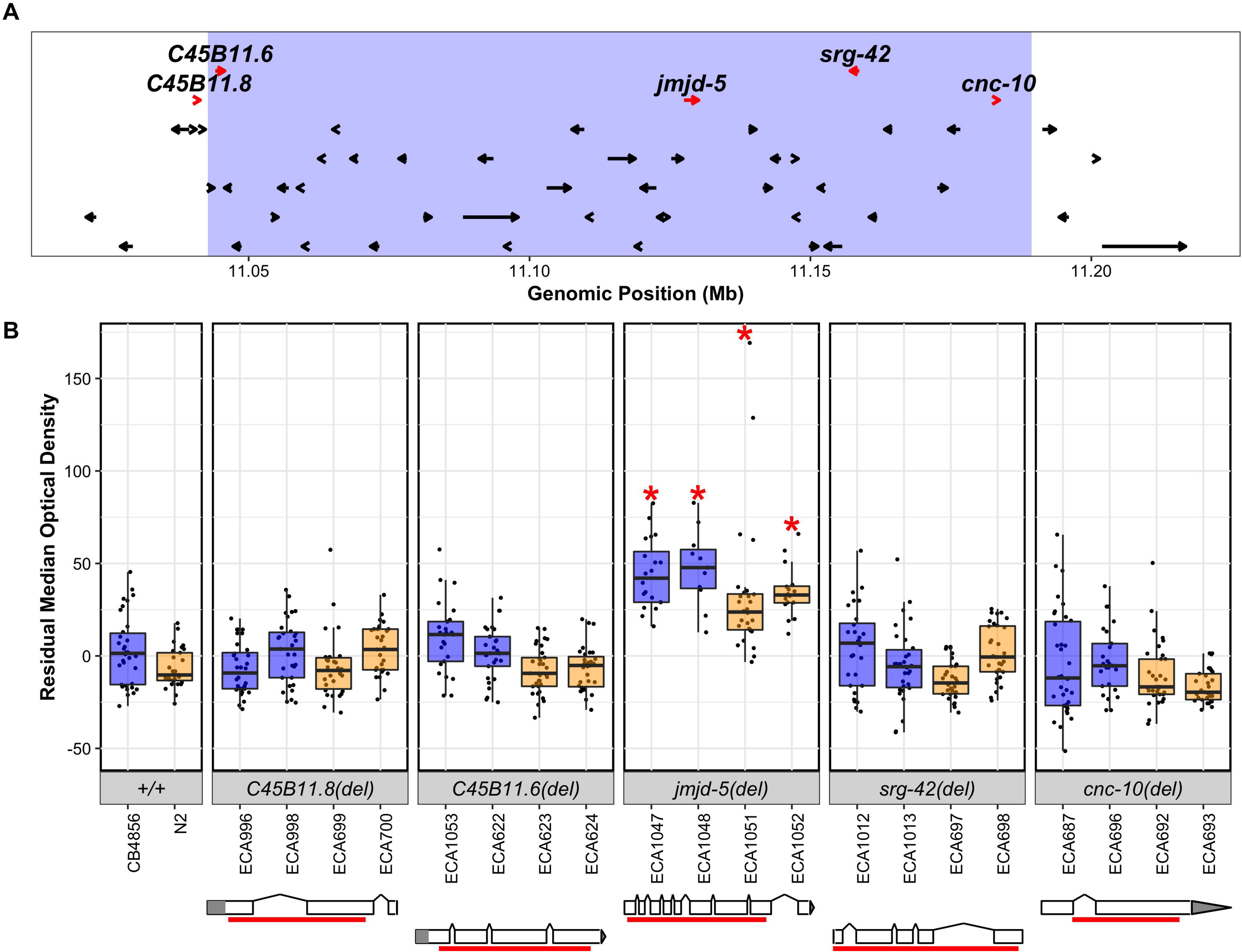
Bleomycin responses of the deletion alleles for each candidate gene are shown. **A.** The linkage mapping QTL confidence interval (light blue) with 20 kb on the left and the right is displayed. Each protein-coding gene in the region is indicated by an arrow that points in the direction of transcription. Genes with non-synonymous variants between the N2 and CB4856 strains are shown as red arrows and are labeled with their gene name. **B**. Deletion alleles for each of these genes were tested in response to bleomycin. Bleomycin responses are shown as Tukey box plots, with the strain name on the x-axis, split by gene, and residual median optical density on the y-axis. Each point is a biological replicate. Strains are colored by their background genotype (orange indicates an N2 genetic background, and blue indicates a CB4856 genetic background). For each gene, two independent deletion alleles in each background were created and tested. Red asterisks indicate a significant difference (*p* < 0.05, Tukey HSD) between a strain with a deletion and the parental strain that has the same genetic background. Depictions of each deletion allele are shown below the phenotype for each candidate gene. White rectangles indicate exons and diagonal lines indicate introns. The 5’ and 3’ UTRs are shown by grey rectangles and triangles, respectively. The region of the gene that was deleted by CRISPR-Cas9 directed genome editing is shown as a red bar beneath each gene model.

To test these genes in bleomycin-response variation, we systematically deleted each of the candidate genes in both the N2 and CB4856 backgrounds. We used CRISPR-Cas9 mediated genome editing to generate two independent deletion alleles of each gene in each genetic background to reduce the possibility that off-target mutations could cause phenotypic differences (Materials and Methods, Supplemental Information). We tested the bleomycin response of each deletion allele in comparison to the N2 and CB4856 parental strains (**Figure 3B, File S3**). The deletion alleles of *C45B11.8*, *C45B11.6*, *srg-42*, and *cnc-10* each had a bleomycin response similar to the respective parent genetic background, which suggested that the functions of each of these genes did not affect bleomycin responses (**Figure 3B**, **File S7**, *p* > 0.05, Tukey HSD). By contrast, the *jmjd-5* deletion alleles in the N2 and the CB4856 backgrounds were each more resistant to bleomycin than their respective parental strains (**Figure 3B, File S7**, ECA1047 vs. CB4856 *p* = 3.8e-10, ECA1048 vs. CB4856 *p* = 0.026, ECA1051 vs. N2 *p* = 7.4e-4, ECA1052 vs. N2 *p* = 2.9e-6, Tukey HSD). However, we also noted that these strains were more sensitive in the control condition than other deletion strains (**Figure S8, File S1, File S7**). Therefore, the relative increased bleomycin resistance observed in the *jmjd-5* deletion strains could be caused by their increased sensitivity in the control condition.

We tested if the non-synonymous variant in *jmjd-5* between the N2 and CB4856 strains caused bleomycin-response differences. At residue 338 of JMJD-5, the N2 strain has a proline, whereas the CB4856 strain has a serine (S338P, **Figure S9A**). We used CRISPR-Cas9 to generate reciprocal allele replacements of the *jmjd-5* single-nucleotide polymorphism that encodes the putative amino-acid change in the N2 background *jmjd-5(N2 to CB4856)* and in the CB4856 background *jmjd-5(CB4856 to N2)* (Materials and Methods, Supplemental Information). We created two independent allele replacements in each genetic background and measured each strain for bleomycin-response differences as compared to the parental strains (**Figure S9B, File S3**). Although the allele-replacement strains with the CB4856 allele in the N2 genetic background *jmjd-5(N2 to CB4856)* were significantly different from the N2 parental strain, these strains were more sensitive to bleomycin than N2 (**Figure S9B, File S7**, ECA576 vs. N2 *p* = 0.006, ECA577 vs. N2 *p* = 1.6e-6, Tukey HSD). This increased sensitivity was unexpected, because the CB4856 allele at the *jmjd-5* locus should confer resistance. However, the NIL with the CB4856 genotype across the QTL was not different from the N2 parental strain (ECA232 in **Figure S6, File S7**), suggesting that the QTL might only confer increased sensitivity when the N2 allele is in the CB4856 background. Therefore, it remained unclear whether an allele replacement of *jmjd-5* in the N2 parental background could confer resistance. Neither of the two strains with the N2 allele in the CB4856 background, *jmjd-5(CB4856 to N2)*, conferred a significantly more sensitive phenotype than the CB4856 parental strain (**Figure S9B, File S7**). Given that the QTL explained 43.58% of phenotypic variation among the RIAILs, the causal variant should have a clear impact on bleomycin response. Additionally, the NILs with the N2 allele at the QTL introgressed into the CB4856 background displayed a significant increase in bleomycin sensitivity compared to the parental CB4856 strain (**Figure S6, File S7**). Taken together, the phenotypes of the reciprocal allele-replacement strains showed that the amino-acid change in JMJD-5 likely does not underlie bleomycin-response variation between the N2 and CB4856 strains, although deletion of this gene did cause resistance to bleomycin regardless of the genetic background. We performed a reciprocal hemizygosity assay to test if natural variation in *jmjd-5* function affected bleomycin responses (**Figure S10, Figure S11, Figure S12**). The results of this assay supported the previously identified increase in bleomycin resistance of homozygous *jmjd-5* deletions in both parental backgrounds (**Figure S11, File S7,** *p* < 0.05, Tukey HSD), which again might be caused by an increased sensitivity in the control condition (**Figure S12**). However, the increases in bleomycin resistance between each *jmjd-5* deletion strain and the strain with the same genetic background were similar, and the reciprocal hemizygous strains show equivalent bleomycin responses (**Figure S11**). Taken together, these results suggest that natural variation in *jmjd-5* function does not underlie this QTL.

### The nematode-specific gene *H19N07.3* impacts bleomycin variation

Because none of the genes with a non-synonymous variant between the N2 and CB4856 strains explained the QTL, we explored other ways in which natural variation could impact bleomycin responses. We used the 13,001 SNPs to perform linkage mapping on the gene expression data of the RIAILs and identified 4,326 expression QTL (eQTL) across the genome (Rockman *et al*. 2010, **File S12, File S13**, Materials and Methods). Of the 118 genes in the 187 kb surrounding the bleomycin-response QTL, expression for 50 genes were measured in the previous microarray study. We identified a significant eQTL for eight of these 50 genes, four of which mapped to chromosome V (**File S13**). eQTL for two of those four genes, *H19N07.3* and *cnc-10,* mapped to the center of chromosome V and overlapped with the bleomycin-response QTL. Because *cnc-10* did not underlie bleomycin response variation (**Figure 3B**), we hypothesized that *H19N07.3* might underlie the bleomycin-response QTL. The *H19N07.3* eQTL explains 45.70% of the variation in *H19N07.3* expression among the RIAILs (**Figure 4A and B, File S14**). The length of animals and expression of *scb-1* was correlated in the RIAIL strains (**Figure 4C, File S15**, *r*^2^= 0.61, *p* < 9.5e-13 Spearman’s correlation). Although this gene does not have a non-synonymous variant between the N2 and CB4856 strains, natural variation in gene expression could impact bleomycin responses.

**Figure 4:**
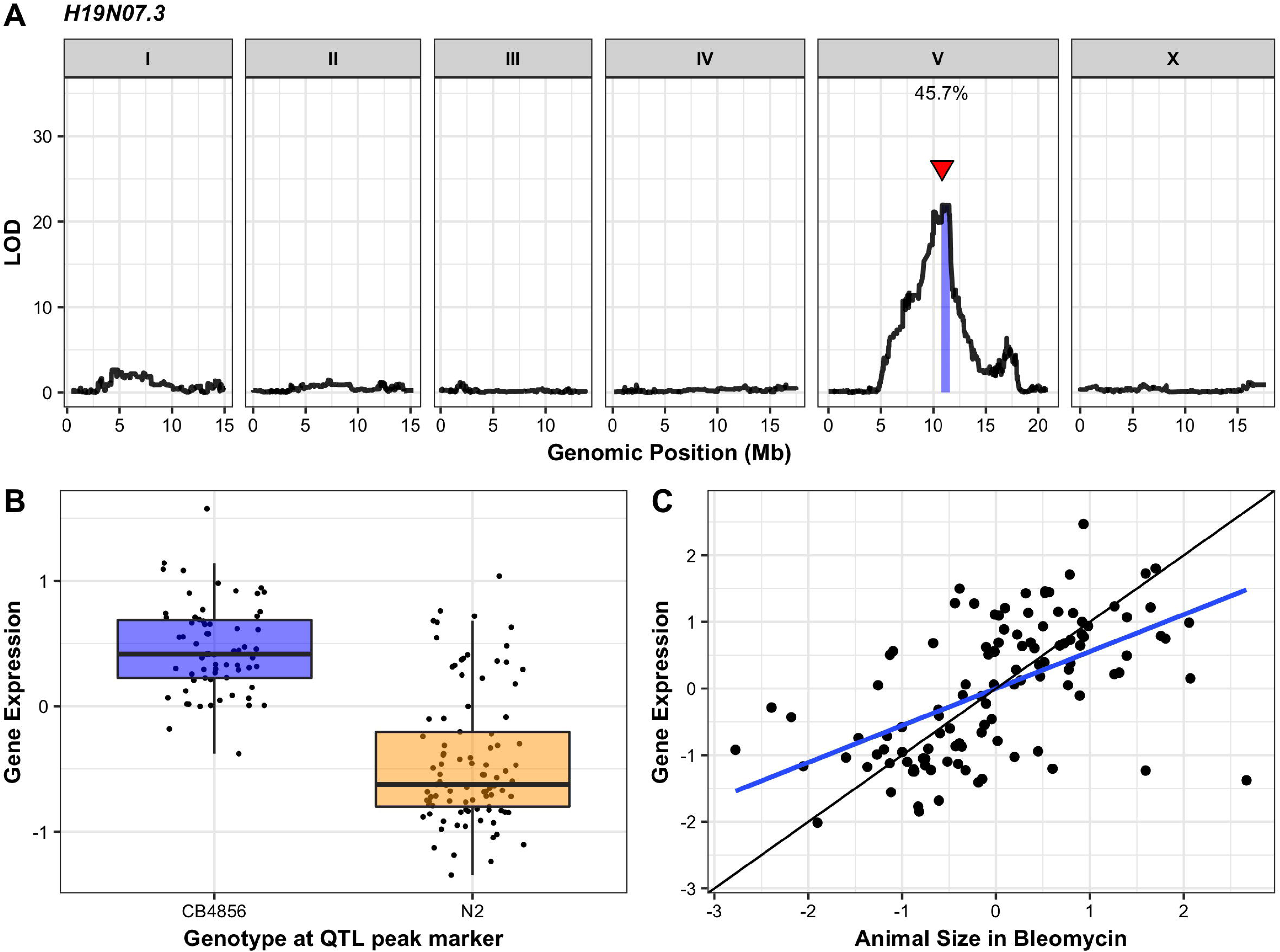
Linkage mapping of the *H19N07.3* expression difference among RIAILs is shown. **A.** On the x-axis, each of 13,001 genomic markers, split by chromosome, were tested for correlation with *H19N07.3* expression variation across the RIAIL panel. The log of the odds (LOD) score for each marker is reported on the y-axis. The significant quantitative trait locus (QTL) is indicated by a red triangle at the peak marker, and a blue ribbon shows the 95% confidence interval around the peak marker. The total amount of expression variance across the RIAIL panel explained by the genotype at the peak marker is printed as a percentage. **B.** RIAIL gene expression (y-axis), split by allele at the QTL peak marker (x-axis) is shown. Phenotypes of RIAILs containing the N2 allele (orange) are compared to RIAILs containing the CB4856 allele (blue). Phenotypes are shown as Tukey box plots, and each point is the *H19N07.3* expression of an individual strain. **C.** The correlation between animal size in bleomycin and *H19N07.3* expression is shown as a scatterplot, with each RIAIL shown as a point. Each axis was scaled to have a mean of zero and a standard deviation of one. The line of best fit is shown in blue. The identity line is shown in black for reference.

We created two independent CRISPR-Cas9 mediated deletion alleles of *H19N07.3* in the N2 and the CB4856 backgrounds and measured the bleomycin responses of these strains compared to the parental strains (**Figure 5, File S3, File S7,** Supplemental Information, Materials and Methods). Each *H19N07.3* deletion strain was more sensitive to bleomycin than the respective parental strain (**Figure 5, File S3, File S7,** ECA1133 vs. CB4856 *p* < 1.4e-14, ECA1134 vs. CB4856, *p* < 1.4e-14, ECA1132 vs. N2, *p* = 6.9e-5, ECA1135 vs. N2, *p* = 0.006, Tukey HSD). These results suggest that *H19N07.3* function is required for resistance to bleomycin. Therefore, we renamed this gene *scb-1* for sensitivity to the chemotherapeutic bleomycin. Unlike with the *jmjd-5* deletion strains, the *scb-1* deletion strains had no significant differences in the control condition (**File S7**). Therefore, the bleomycin sensitivity of the *scb-1* deletion strains were not caused by control-condition phenotypes.

**Figure 5:**
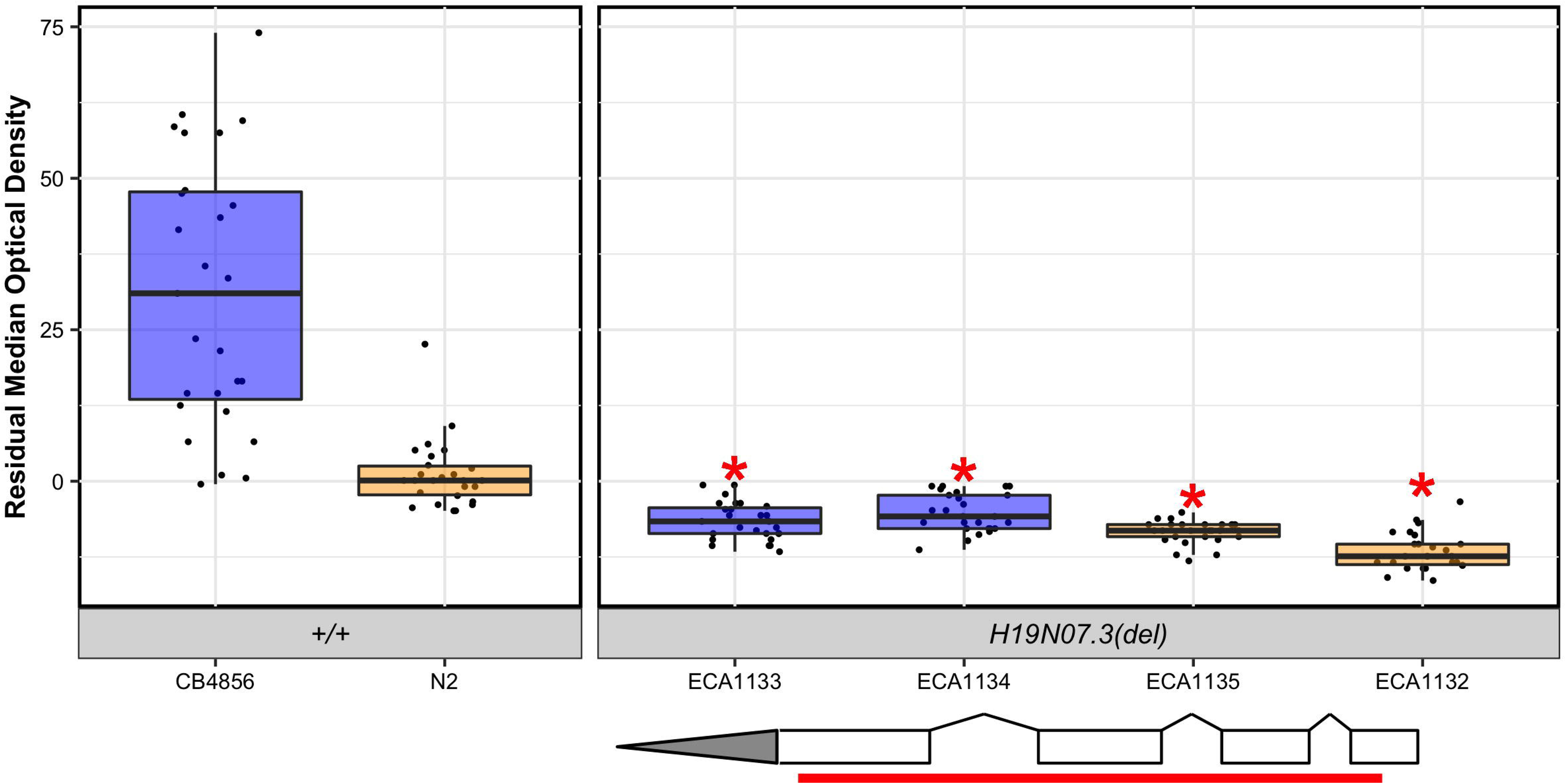
Bleomycin responses of *H19N07.3* deletion alleles are shown as Tukey box plots, with the strain name on the x-axis, split by genotype, and residual median optical density on the y-axis. Each point is a biological replicate. Strains are colored by their background genotype (orange indicates an N2 genetic background, and blue indicates a CB4856 genetic background). Two independent deletion alleles in each genetic background were created and tested. Red asterisks indicate a significant difference (*p* < 0.05, Tukey HSD) between a strain with a deletion and the parental strain that has the same genetic background. A depiction of the deletion allele is shown below the box plots. White rectangles indicate exons, and diagonal lines indicate introns. The 5’ and 3’ UTRs are shown by grey rectangles and triangles, respectively. The region of the gene that was deleted by CRISPR-Cas9 directed genome editing is shown as a red bar beneath the gene model.

Because an *scb-1* non-synonymous variant has not been identified between the N2 and CB4856 strains, changes to protein function likely do not cause bleomycin response differences. RIAILs with the CB4856 allele at the QTL peak marker have increased expression of *scb-1* and increased bleomycin resistance compared to RIAILs with the N2 allele (**Figure 1**, **Figure 4**). Therefore, *scb-1* expression differences might cause the bleomycin-response variation between the parental strains. We performed RNA-seq of the N2 and CB4856 strains to assess *scb-1* expression differences between the parental strains and did not identify a significant increase in expression in the CB4856 strain (**Figure S13, File S16,** *p* = 0.20, Wald test; *p* = 0.17, likelihood ratio test). This result could be caused by the low sample size (n = 3) in the RNA-seq experiment, or the RIAIL strains could have a novel variant that arose during strain construction that causes *scb-1* expression variation. Alternatively, the expression difference observed in the RIAIL strains could be attributed to epistatic loci. We performed a two-factor genome scan to identify epistatic loci that underlie *scb-1* expression variation in the RIAILs and identified two significant interactions: one between loci on chromosomes IV and X and another between loci on chromosomes II and V (**Figure S14, File S17**). This result might suggest that epistatic loci underlie *scb-1* expression variation in the RIAILs and could explain why *scb-1* expression is not variable in the parental strains.

To test the role of natural variation in *scb-1* function, we performed a reciprocal hemizygosity test in control and bleomycin conditions (**Figure 6, File S3**). These results matched the increase in sensitivity of homozygous deletions in both parental backgrounds observed previously. The hemizygous strain with the CB4856 allele of *scb-1* had a bleomycin phenotype similar to the heterozygous strain, whereas the hemizygous strain with the N2 allele of *scb-1* was more sensitive to bleomycin than the heterozygous strain. Taken together, these results suggest that natural variation in *scb-1* function underlies the bleomycin-response difference between the N2 and CB4856 strains.

**Figure 6:**
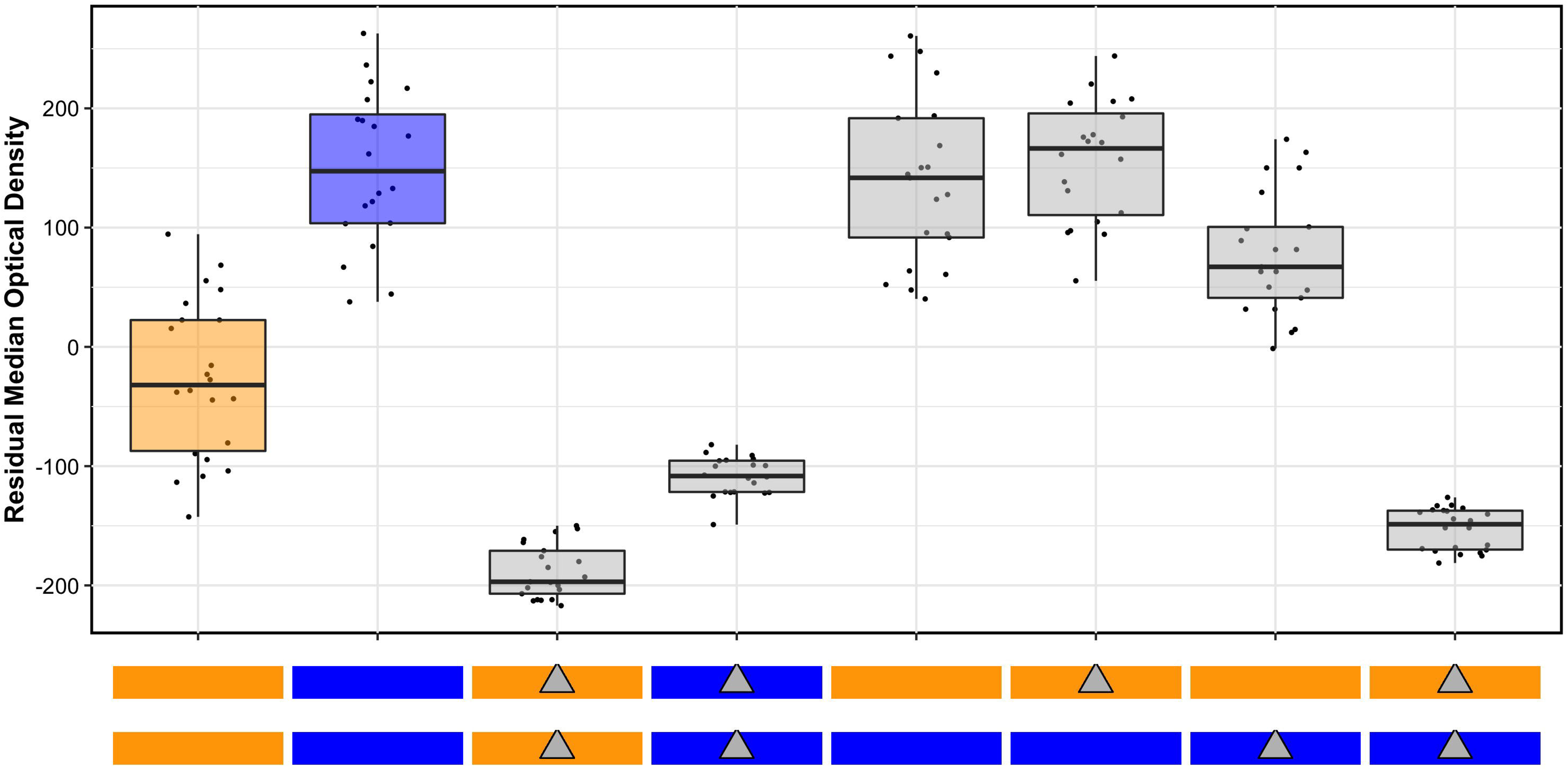
Results of the *scb-1* reciprocal hemizygosity assay are shown. The y-axis shows the residual median optical density for each strain reported along the x-axis. Bleomycin responses are reported as Tukey box plots where each point is a biological replicate. The genotypes of each strain are shown as colored rectangles beneath each box plot, where each rectangle represents a homolog (orange rectangles are an N2 genotype, and blue rectangles are a CB4856 genotype). Maternal homologs are shown on top and paternal homologs are shown on bottom. Grey triangles indicate a deletion of *scb-1,* placed on the rectangle showing the background into which the deletion was introduced. The box plots for the parental strains (N2 and CB4856, on the left) are colored according to genotype.

### Differences in *scb-1* function might be regulated by a rare variant

The *scb-1* natural variant that underlies the bleomycin-response differences remains unknown. Because this gene does not have a predicted non-synonymous variant between the N2 and CB4856 strains, *scb-1* gene expression might underlie bleomycin response differences. Potential candidate variants that could cause this expression difference include one variant two kilobases upstream of the gene and one variant in the third intron of *scb-1*. However, gene expression can be regulated by distant loci, so the identification of the specific variant is difficult. To understand whether natural variation of *scb-1* underlies bleomycin-response differences in other strains, we compared the bleomycin-response linkage mapping to a genome-wide association mapping (GWA). We used the HTA to measure median optical density in bleomycin for 83 divergent wild isolates and performed GWA mapping (**Figure 7, File S18, File S19**). Six QTL were identified from the GWA, but none of these QTL regions overlapped the QTL from linkage mapping (**Figure 7, File S19**). Therefore, the CB4856 strain might have a rare variant that underlies its increase in bleomycin resistance compared to the N2 strain.

**Figure 7:**
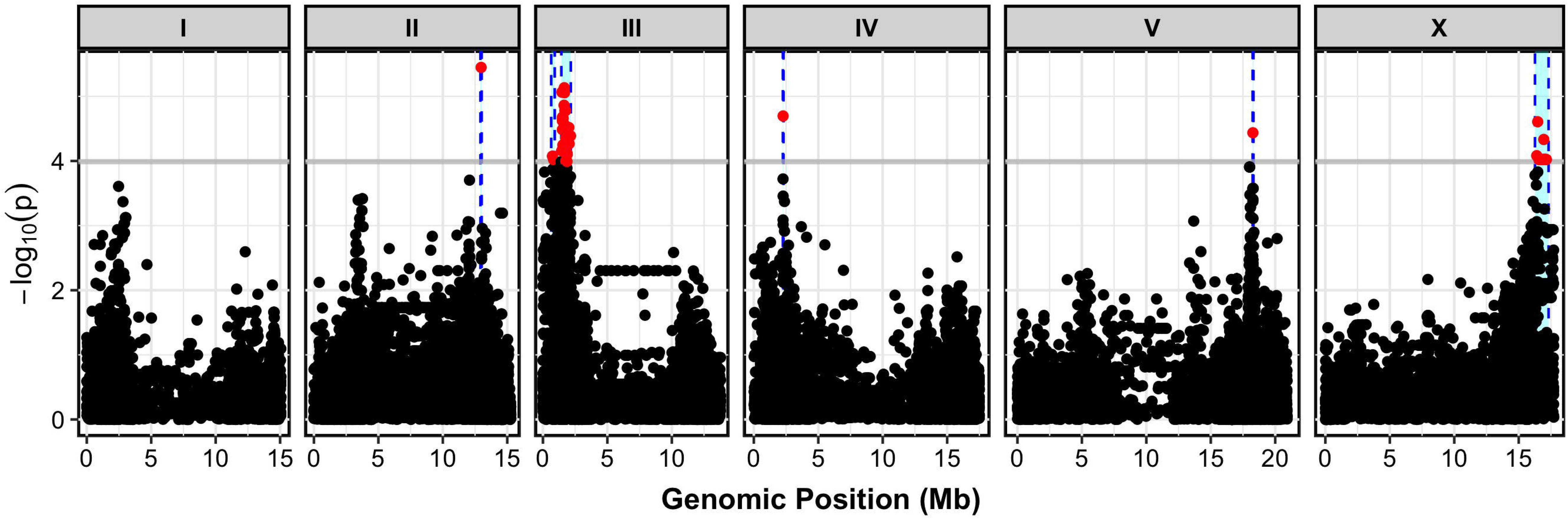
A genome-wide association study for residual median optical density in bleomycin for 83 wild isolates is shown. On the x-axis, each genomic marker, split by chromosome, was tested for correlation with phenotypic variation across the wild isolates. The log_10_(*p*) value of these correlations are reported on the y-axis. Each marker that reached a significance threshold determined by eigenvalue decomposition of the SNP correlation matrix is colored in red. QTL regions of interest are indicated by blue regions surrounding the significant markers.

We identified all single nucleotide variants (SNVs), small insertion/deletions (indels), and structural variants (SVs) present in these 83 strains for which the CB4856 strain contains the alternate allele compared to the N2 reference strain. We found 105 variants within the QTL confidence interval (79 SNVs, 26 indels, 0 SVs) for which the CB4856 strain contains the alternate allele (**Figure S15, File S20)**. We then identified SNVs and indels with a minor-allele frequency less than 5% within the 83 strains, because these low-frequency variants are likely to have insufficient power to map by GWA. Seventy-two of the 105 variants in the region were identified as rare variants that might underlie the bleomycin-response difference between the N2 and CB4856 strains (**Figure S15, File S20**). Twenty-eight of these rare variants were not unique to CB4856, and other strains in the wild isolate panel shared these variants. However, none of these variants showed phenotypic trends consistent with an alternate allele conferring resistance to bleomycin (**Figure S16**). Forty-four of the 72 rare variants were unique to CB4856 within this set of 83 strains. One or more of these 44 variants could underlie the bleomycin-response QTL, but further work must be performed to identify which, if any, of these variants underlies the *scb-1* bleomycin-response difference between N2 and CB4856.

We searched for homologs of *scb-1* in other species using a BLASTp search (www.wormbase.org, Release WS268) and identified homologs in nine other *Caenorhabditis* species but none outside of the Nematoda phylum (**Figure S17**, **File S21, File S22**) (Edgar 2004a; b). None of the homologs of SCB-1 have previously identified functions. We used Phyre2 to predict protein domains within the SCB-1 protein and were unable to detect any functional domains by sequence homology. Twenty-three percent of the SCB-1 protein sequence matched a hydrolase of a Middle East respiratory syndrome-related coronavirus (Zhang *et al*. 2018). However, the low confidence of the model (21.5%) should be considered before making conclusions about the function of *scb-1* based on these results.

## DISCUSSION

Here, we performed linkage mapping of bleomycin-response variation and identified a highly significant QTL on chromosome V. We tested all six candidate genes in the QTL region to identify a causal gene that underlies bleomycin-response variation between two divergent strains. Deletions of four of these genes, *C45B11.8*, *C45B11.6*, *srg-42*, and *cnc-10* did not impact bleomycin responses. Deletions in one gene, *jmjd-5*, showed increased bleomycin resistance in both parental backgrounds. However, we concluded that the QTL cannot be explained by differences in *jmjd-5* after further analysis of allele-replacement strains and hemizygosity tests. Deletions in a gene with an expression difference, *scb-1* (previously named *H19N07.3*), caused an increase in bleomycin sensitivity in both the N2 and the CB4856 strains. Results from a reciprocal hemizygosity assay indicated that natural variation in *scb-1* function caused differences in bleomycin responses between the N2 and CB4856 strains. Because loss-of-function alleles in *scb-1* caused increased bleomycin sensitivity (**Figure 5**) and the RIAILs with lower *scb-1* expression levels show increased bleomycin sensitivity (**Figure 4**), natural differences in *scb-1* expression might cause the bleomycin-response variation between the N2 and CB4856 strains.

The function of *scb-1*, and particularly how it regulates bleomycin response, remains unknown. A previous study found that RNAi of *scb-1* impaired the DAF-16/FOXO-induced lifespan extension of *daf-2(e1370ts)* mutants, which suggests that *scb-1* might play a role in stress response (Riedel *et al*. 2013). Because bleomycin causes double-stranded DNA breaks and introduces oxidative stress to cells (Stubbe and Kozarich 1987), reduction of *scb-1* function might inhibit the ability of an animal to respond to bleomycin. This model is in agreement with our observation that *scb-1* deletions and RIAILs with lower *scb-1* expression are sensitive to bleomycin (**Figure 4**, **Figure 5**). We used the amino acid sequence of SCB-1 to query the Phyre2 database and found weak homology to a viral hydrolase (Kelley *et al*. 2015; Zhang *et al*. 2018). This result could suggest that SCB-1 might function as a hydrolase, which could be the mechanism by which *scb-1* regulates cellular stress. This finding would be similar to clinical studies that have suggested a role of bleomycin hydrolase (BLMH) in bleomycin-response variation (Lazo and Humphreys 1983; Nuver *et al*. 2005; de Haas *et al*. 2008; Gu *et al*. 2011; Altés *et al*. 2013). Because *scb-1* is expressed in the nucleus of all somatic cells, this gene might impact the ability of bleomycin to cause DNA damage within the cell nucleus (Turek *et al*. 2016). Alternatively, *scb-1* could impact bleomycin import, export, or another mechanism. If the mechanism of *scb-1* is conserved in humans, this discovery could offer insights into the clinical applications of bleomycin. Our results also suggested the presence of genes that interact with *scb-1* to cause bleomycin-response differences. These interacting genes could be conserved in humans and therefore inform the use of bleomycin in the clinic.

Despite its lack of conservation in humans, the SCB-1 protein is homologous to other proteins in other nematode species. Bleomycin is produced by the soil bacterium, *Streptomyces verticillus (Calcutt and Schmidt 1994; Du et al. 2000; Shen et al. 2002)*, which might be found in association with nematodes such as *C. elegans* in the wild (Samuel *et al*. 2016). A shared niche between *C. elegans* and *S. verticillus* could cause bleomycin resistance to be selected. Additionally, the CB4856 wild isolate is more resistant to bleomycin than the laboratory-adapted strain, N2. In fact, the N2 strain is the most sensitive to bleomycin across all strains tested in our HTA (**Figure S16**), which could indicate that bleomycin resistance is beneficial for wild isolates. Given its potential role in the highly conserved insulin-like pathway, *scb-1* could be beneficial in responses to multiple toxins. Interestingly, the *scb-1* gene lies within a toxin-response QTL hotspot on chromosome V (Evans *et al*. 2018). Understanding the mechanism of the role of *scb-1* in toxin responses might offer insights into evolutionary processes that shaped the genomic diversity of *C. elegans* and other nematode species.

Previous studies have leveraged both linkage mapping and GWA in *C. elegans* to identify genetic variants that underlie drug-response differences (Zdraljevic *et al*. 2017, 2019). In each of these studies, drug-response QTL overlap between linkage mapping and GWA, and variants in common between both mapping strain sets have been shown to underlie drug-response QTL. In the case of the bleomycin response, the linkage-mapping QTL did not overlap with the QTL identified through GWA. Therefore, the variant that underlies the QTL likely is not present at an allele frequency above 5% in the panel of wild isolates used for the bleomycin GWA. The difference between linkage mapping and GWA results indicates that both rare and common natural variants underlie bleomycin-response variation.

This study emphasizes the power of the *C. elegans* model system to dissect complex traits. Although linkage mapping detected a highly significant QTL, the manner in which genetic components affect bleomycin responses is not simple. Certain near-isogenic lines showed transgressive phenotypes (**Figure 2, Figure S6**), which indicates that multiple loci interact to create extreme bleomycin sensitivity in particular strains with mixed genetic backgrounds. Our attempts to identify epistatic loci that underlie bleomycin responses in the RIAILs were unsuccessful, potentially because of the complexity of these epistatic interactions. Despite this complexity, *scb-1* deletions showed increased bleomycin sensitivity in both parental backgrounds, and expression variation among the RIAIL panel mapped to the same locus as the bleomycin response QTL. Interestingly, the parental strains do not seem to vary in *scb-1* expression, as measured by RNA-seq (**Figure S13**). We found evidence of epistatic loci that underlie *scb-1* expression variation in the RIAILs, which might explain why the parental strains do not differ in *scb-1* expression (**Figure S14**). Additional complexities of this trait include the lack of overlap between GWA and linkage mapping QTL and the potential effect of *jmjd-5* loss-of-function on bleomycin responses. Despite the complicated manner in which genetic variants seem to affect bleomycin responses, we leveraged the powerful model of *C. elegans* to identify a single gene that underlies this complex trait.

## ACKNOWLEDGEMENTS

This work was supported by an American Cancer Society Research Scholar Grant to ECA (127313-RSG-15-135-01-DD) to ECA. Additionally, SCB received support from the Biotechnology Training Program training grant (T32GM008449) and the Dr. John N. Nicholson Fellowship. SZ received support from the Cell and Molecular Basis of Disease training grant (T32GM008061) and The Bernard and Martha Rappaport Fellowship. KWB received support from a Northwestern Undergraduate Research Grant. DEC was supported by the National Science Foundation Graduate Research Fellowship (DGE-1324585). YW was supported as a joint PhD student by China Scholarship Council (No. 201706910052). We would like to thank members of the Andersen Lab for helpful comments on the manuscript.

