## Supplemental Figures for "A nematode-specific gene underlies bleomycin-response variation in *Caenorhabditis elegans*"

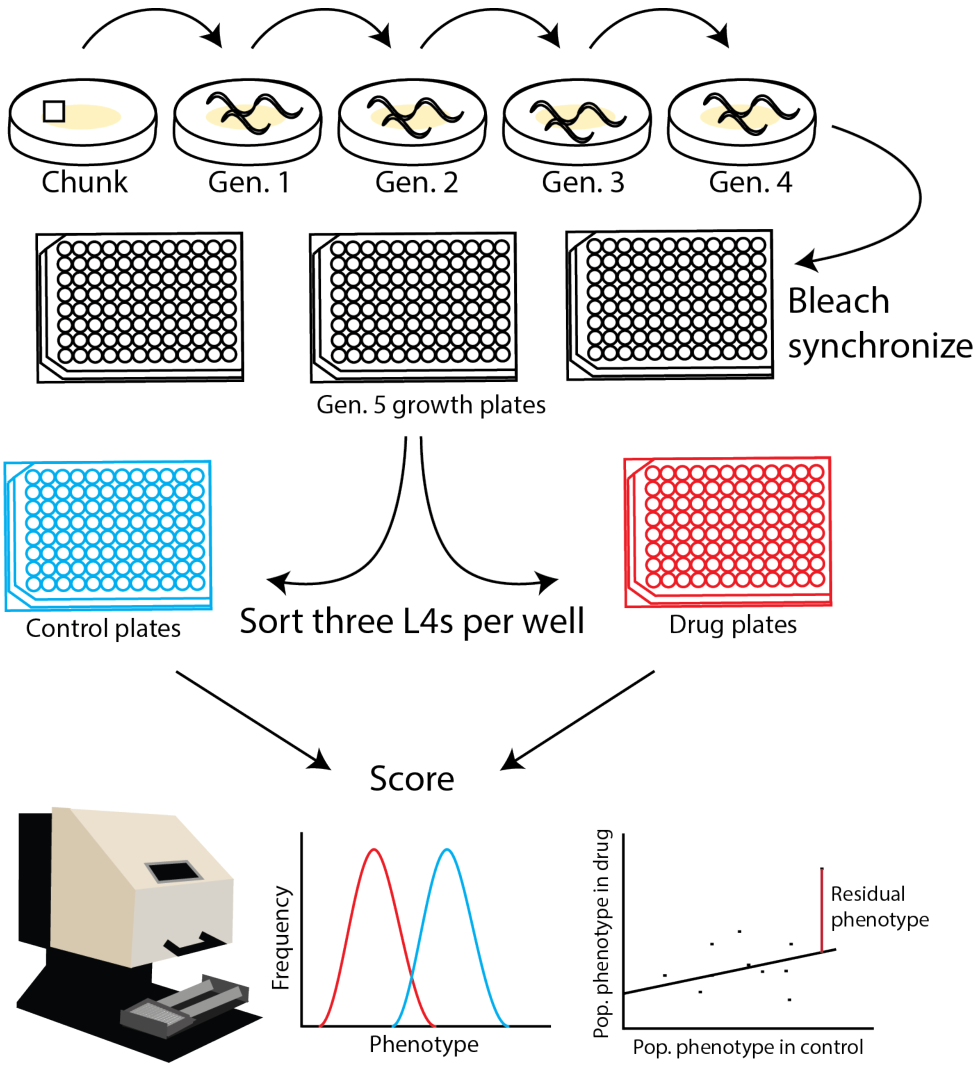


**Figure S1**

A diagram shows the high-throughput fitness assay protocol. From top to bottom, left to right: Each strain is chunked onto a fresh plate, and animals of the L4 stage are transferred for four generations. Gravid adults from the fourth generation are bleached to synchronize growth of all strains. Embryos are aliquoted to 96-well growth plates containing K medium and 5 mg/mL of bacterial lysate. After 48 hours, three L4 larvae are sorted from each well of the growth plate to the corresponding well of either a control plate (which contains K medium, 10 mg/mL bacterial lysate, and 1% distilled water) or a drug plate (which contains K medium, 10 mg/mL bacterial lysate, 1% distilled water, and bleomycin). Animals are grown in the control and drug plates for four days at 20$^{o}$ with shaking. Then, animals are scored using the COPAS BIOSORT. For each well, all animals are measured for length (TOF) and optical density (EXT). The normal distributions represent hypothetical distributions of all animals in a given well of a control (red) plate or a drug (blue) plate. Summary statistics of these distributions are measured to obtain a population growth estimate for all animals within a well. For each trait measured, the average control phenotype for a given strain is plotted against the drug phenotype for each replicate of that strain. A linear model is fit to those data, and the residual phenotype is calculated and used for further analysis.

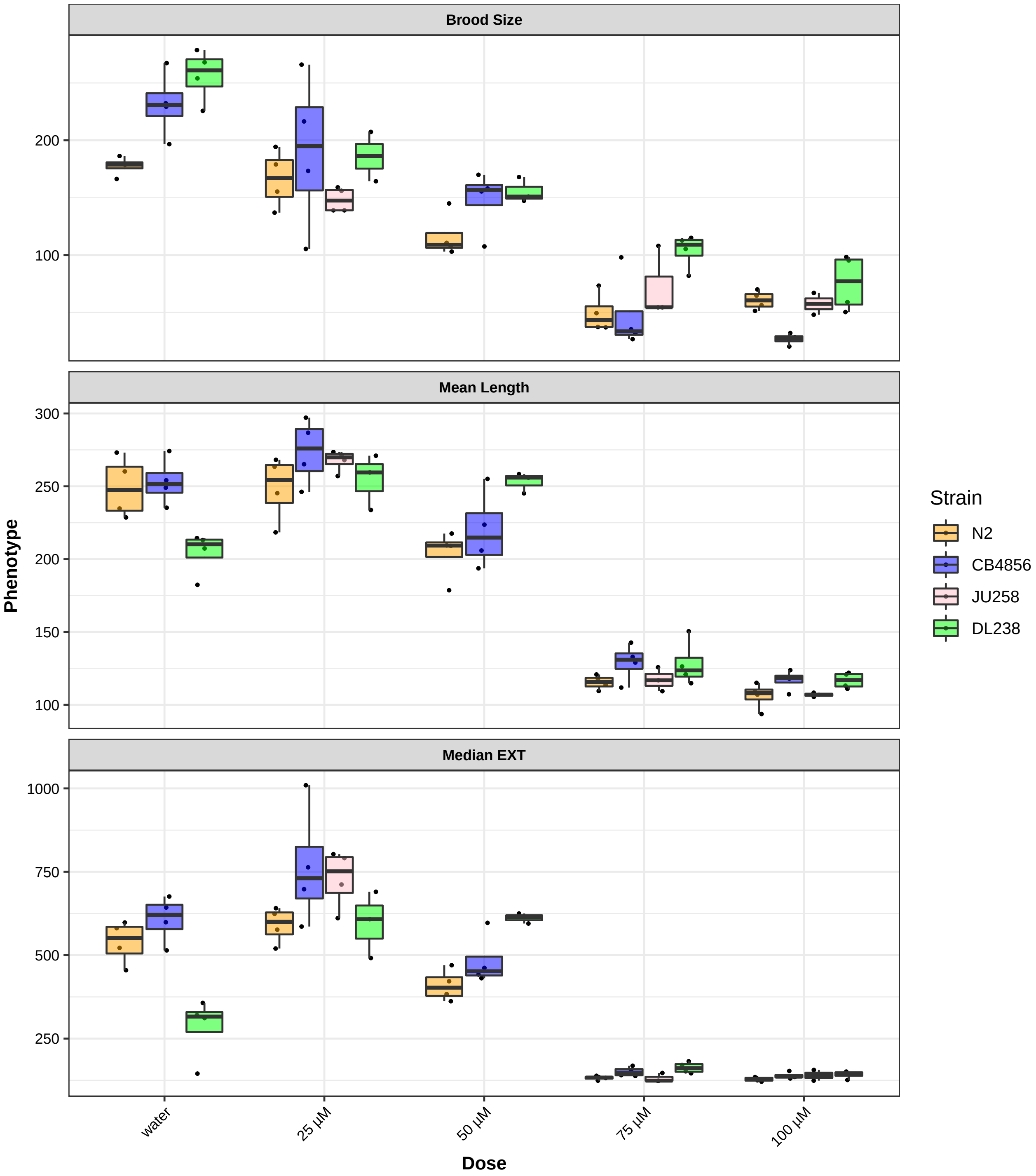


**Figure S2**

Dose-response phenotypes are shown for three high-throughput fitness traits: brood size (norm.n), mean length (mean.TOF), and median optical density (median.EXT). Phenotypes for each of the four wild isolates are shown as Tukey boxplots, colored by strain (N2 - orange, CB4856 - blue, JU258 - pink, DL238 - green). The x-axis shows the concentration of bleomycin (or water) to which the animals were exposed, and the y-axis shows the pruned phenotype. Each point is a biological replicate. The JU258 strain was pruned from the water and 50 µM bleomycin conditions because the wells were contaminated.

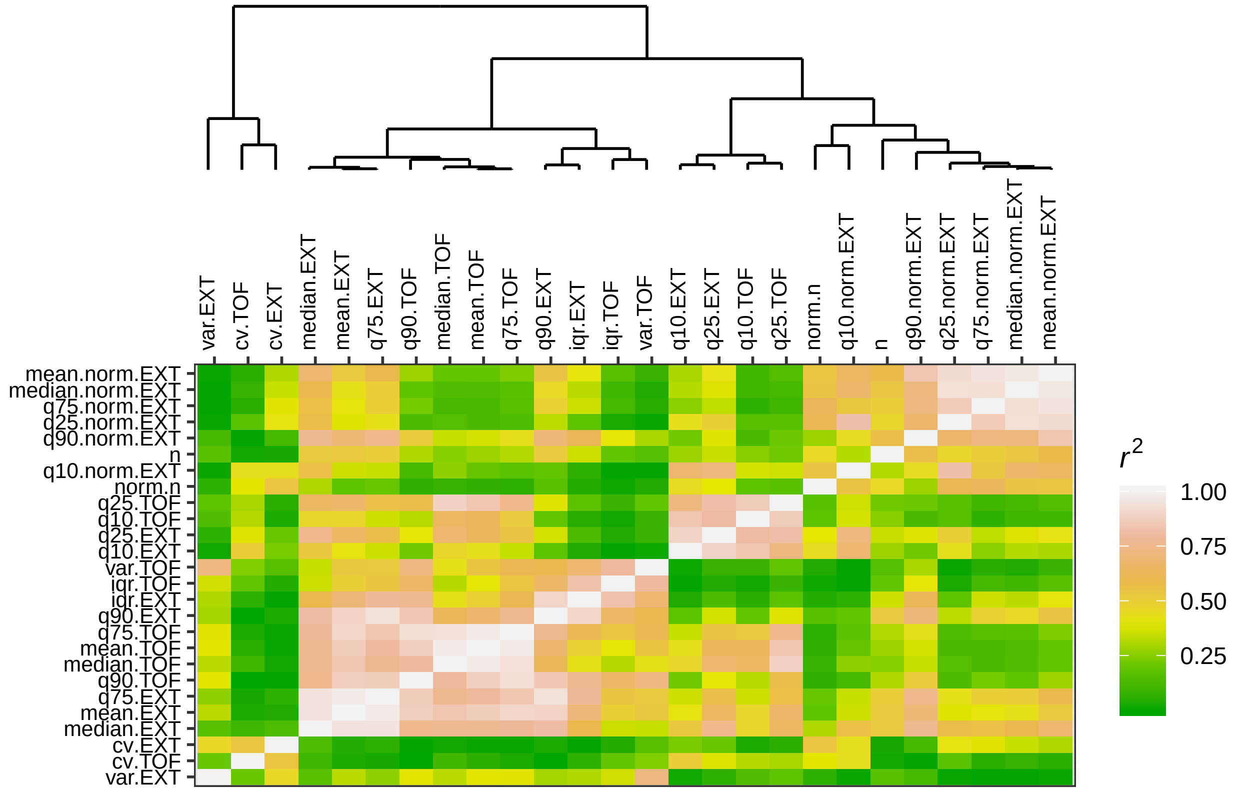


**Figure S3**

Pairwise correlations of high-throughput assay (HTA) traits are shown. (**Top**) A dendrogram of HTA traits, calculated from pairwise correlations of RIAIL phenotypes for each trait, is shown. (**Bottom**) The x- and y-axes list HTA traits for which the pairwise correlation coefficients ($r^2$) are shown as colored tiles, ranging from green to yellow to pink to white.

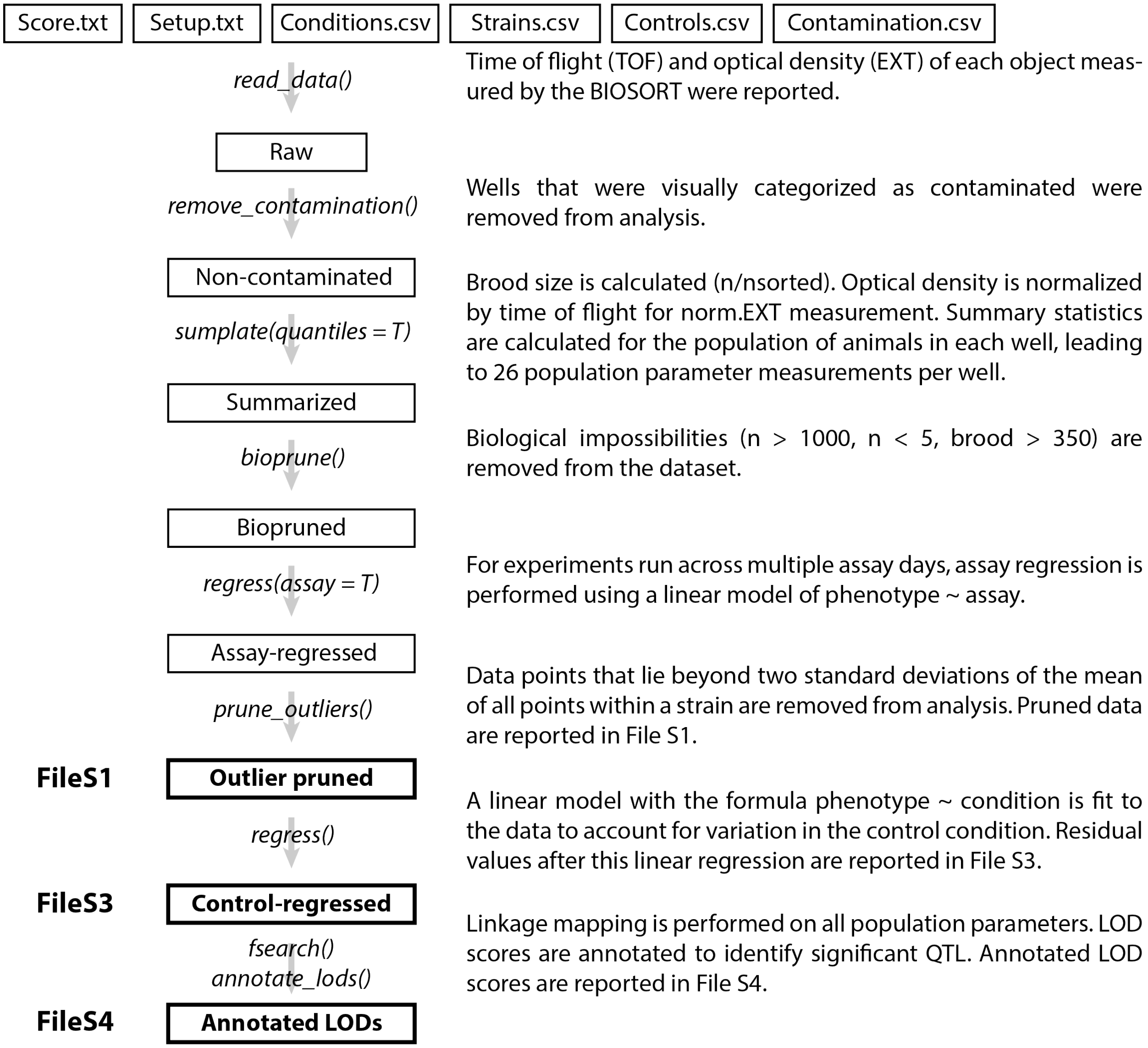


**Figure S4**

A flow chart from raw data to an annotated LOD score dataframe after linkage mapping is shown. **Center:** Intermediate datasets are listed as file names surrounded by a rectangle, and output files in figshare are in boldface with a thick black border. Arrows and italicized text indicate functions in R that were used during analysis. **Left:** File names of data output to figshare. **Right:** Descriptions of analysis steps performed. The *read_data* function merged the setup and score text files from the BIOSORT with csv files that indicated the conditions, strains, controls, and contamination of all wells measured with the high-throughput fitness assay for a particular experiment. Contamination was removed from the raw data using the *remove_contamination* function. Then, *sumplate* calculated brood size, normalized optical density (EXT) to animal length (TOF) for the norm.EXT measurement, and calculated population parameters (including mean, median, variance, covariance, and quantiles) for TOF, EXT, and norm.EXT measurements. Biological impossibilities were removed by the *bioprune* function. If an experiment was measured across multiple days, the *regress(assay = T)* function was used to account for between-day variation. Outliers were removed from each strain using the *prune_outliers* function. These pruned data were output to **File S1**. The *regress* function was used to calculate residual phenotype measurements for all traits within a strain. Regressed data were output to **File S3**. To perform linkage mapping, the *fsearch* and *annotate_lods* functions were used on all traits in **File S3** and mapping results were output to **File S4**.

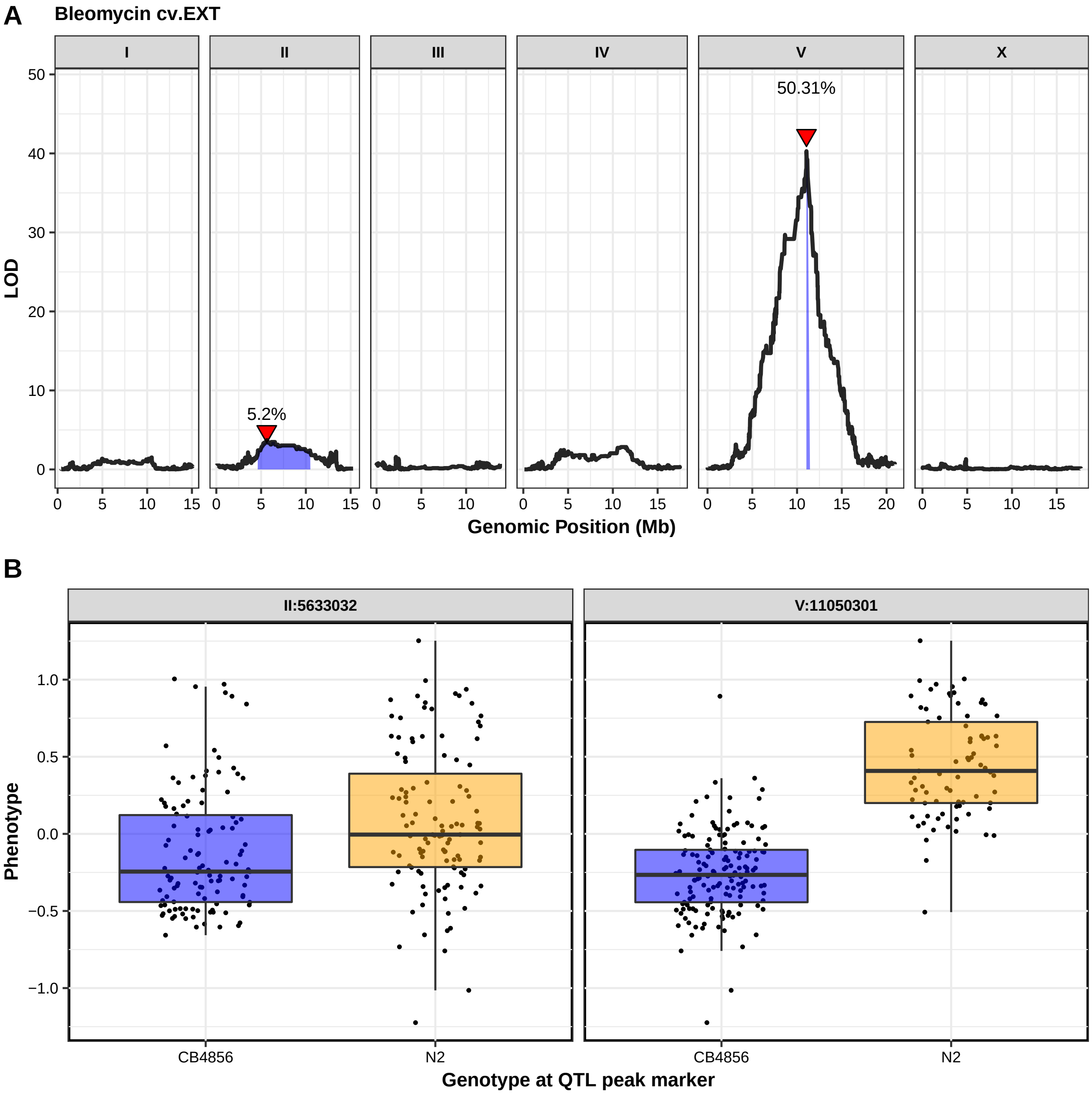
A
Bleomycin cv.EXT
I
II
III
IV
V
X
50.31%
5.2%
LOD
Genomic Position (Mb)
B
II:5633032
V:11050301
Phenotype
CB4856
N2
Genotype at QTL peak marker

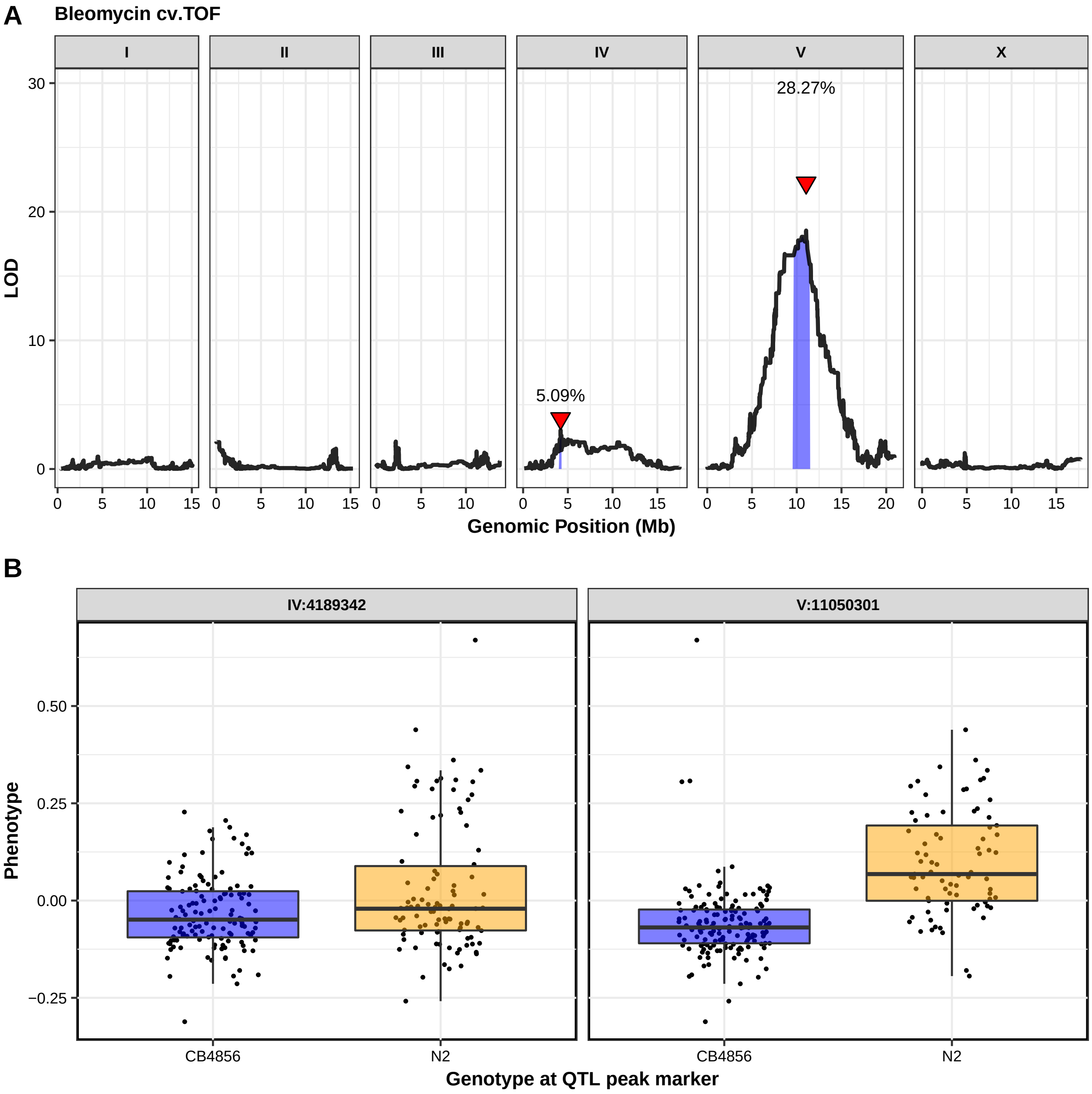
A
Bleomycin cv.TOF
I
II
III
IV
V
X
LOD
Genomic Position (Mb)
5.09%
28.27%
B
IV:4189342
V:11050301
Phenotype
CB4856
N2
Genotype at QTL peak marker

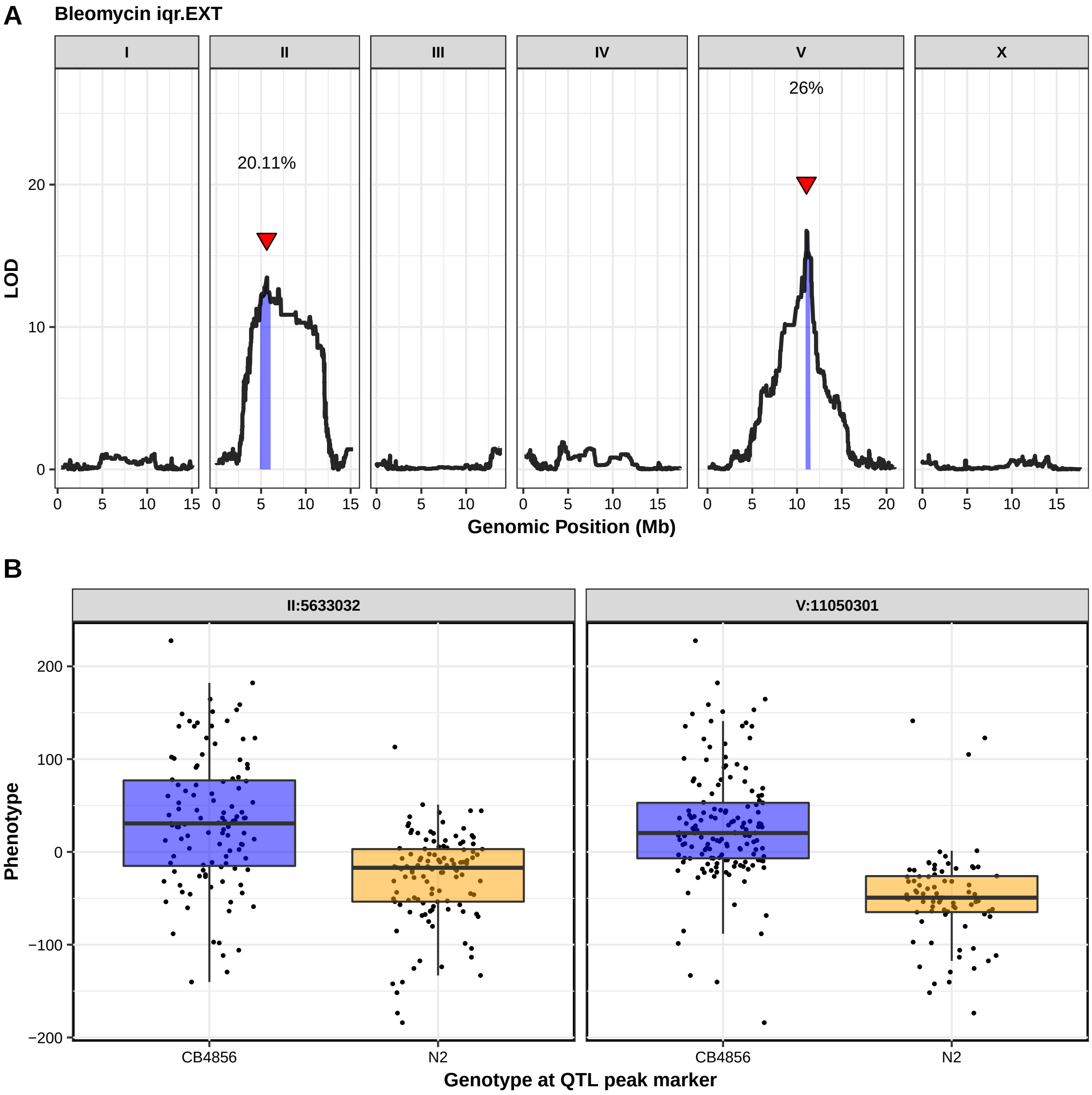
A
Bleomycin iqr.EXT
I
II
III
IV
V
X
20.11%
26%
LOD
20
10
0
0 5 10 15
0 5 10 15 20
Genomic Position (Mb)
B
II:5633032
V:11050301
Phenotype
200
100
0
−100
−200
CB4856
N2
Genotype at QTL peak marker

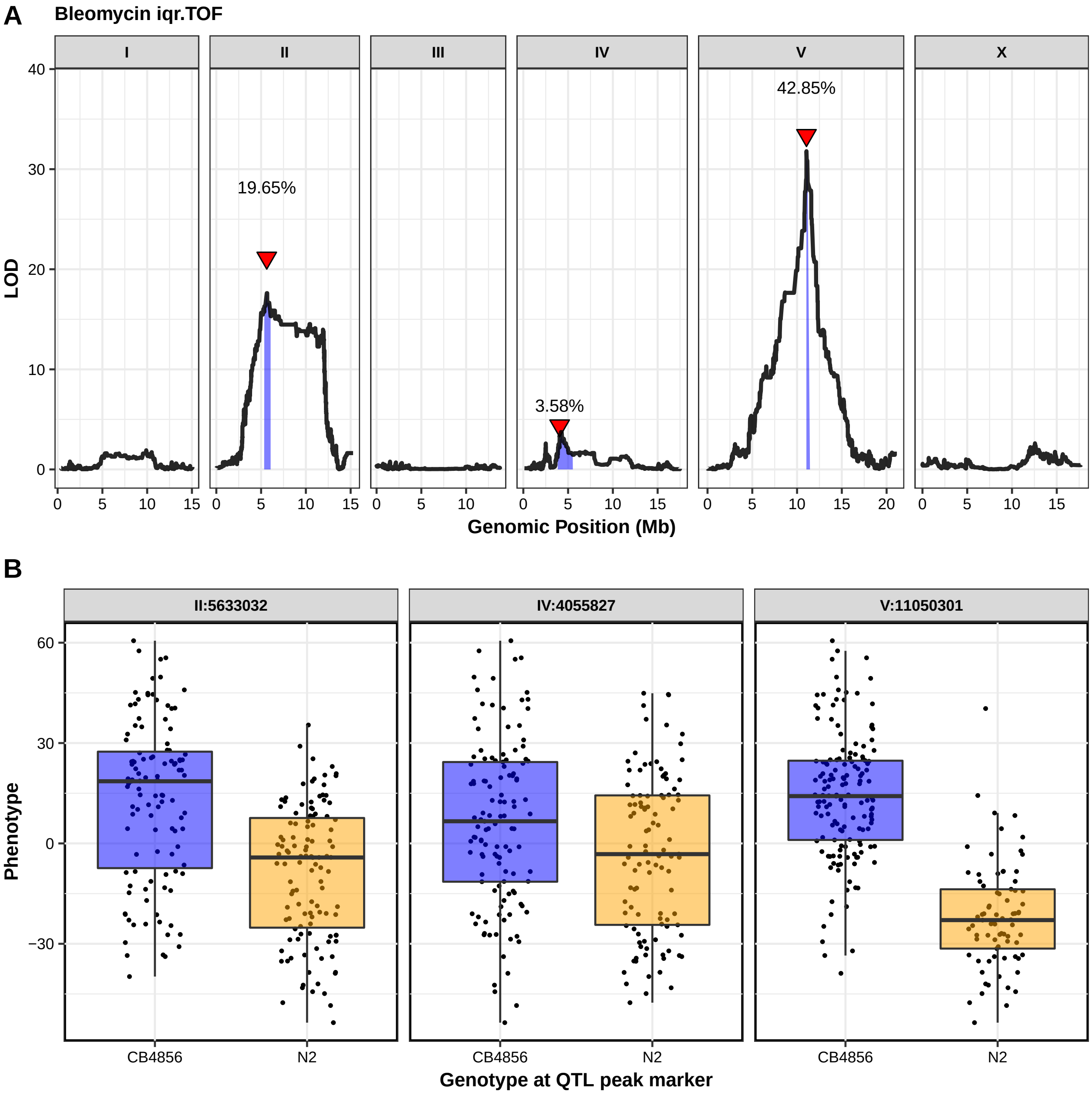

A
Bleomycin iqr.TOF
I
II
III
IV
V
X
LOD
40
30
20
10
0
19.65%
3.58%
42.85%
0
5
10
15
20
Genomic Position (Mb)
B
II:5633032
IV:4055827
V:11050301
Phenotype
60
30
0
−30
CB4856
N2
Genotype at QTL peak marker

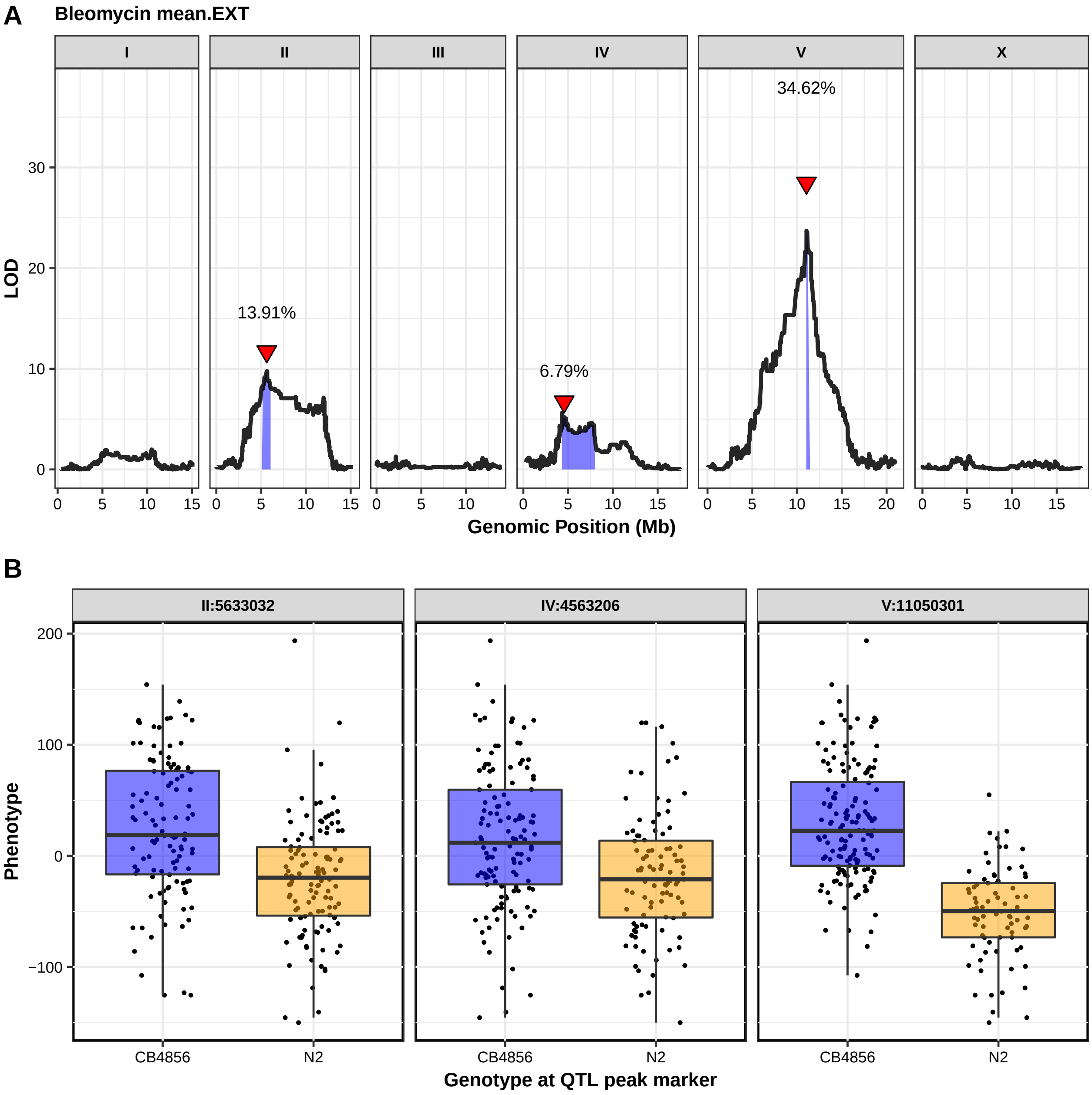
A
Bleomycin mean.EXT
I
II
III
IV
V
X
34.62%
13.91%
6.79%
LOD
30
20
10
0
0 5 10 15
0 5 10 15
0 5 10
0 5 10 15
0 5 10 15 20
0 5 10 15
Genomic Position (Mb)
B
II:5633032
IV:4563206
V:11050301
Phenotype
200
100
0
−100
CB4856
N2
CB4856
N2
CB4856
N2
Genotype at QTL peak marker

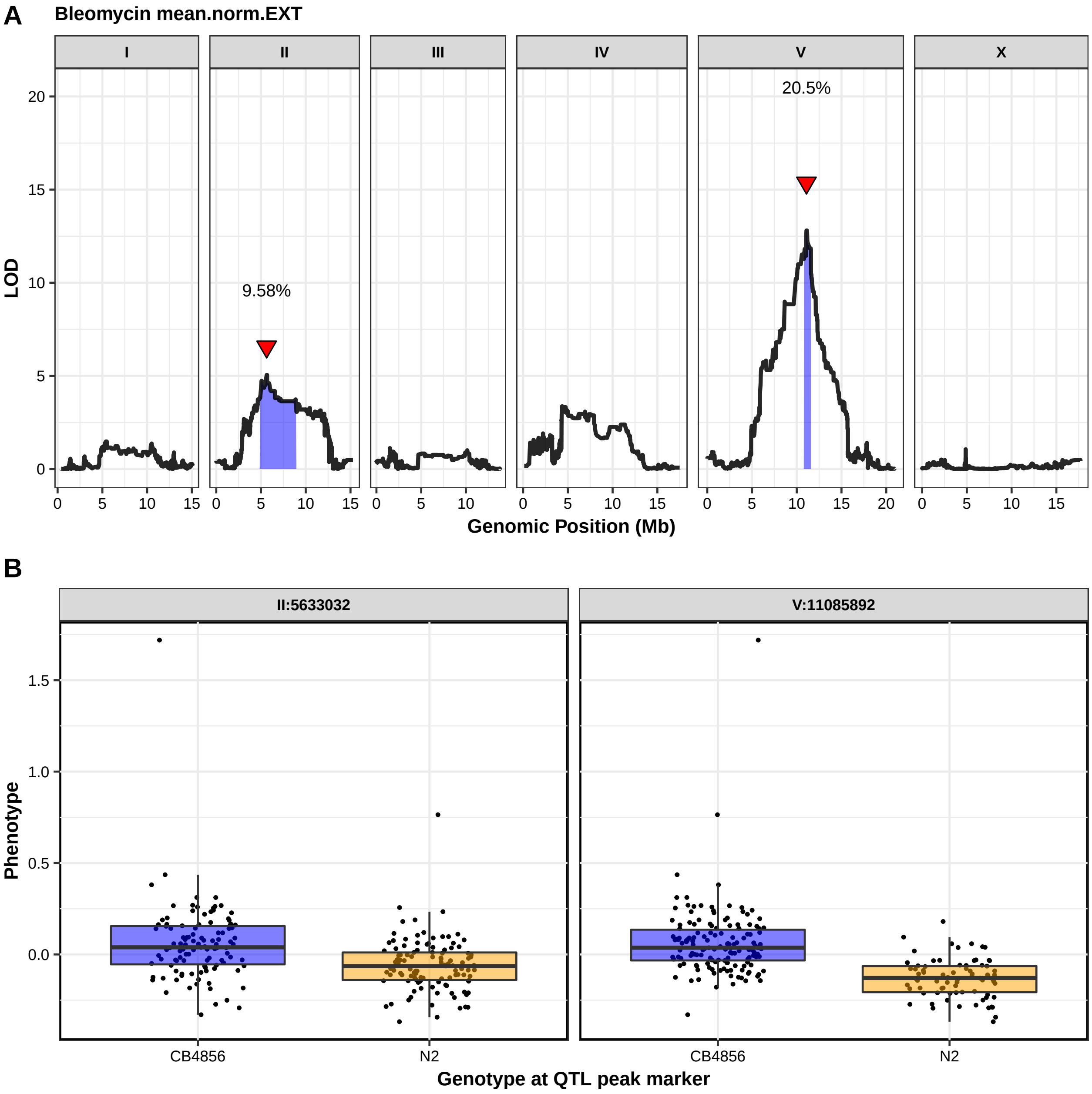
A
Bleomycin mean.norm.EXT
I
II
III
IV
V
X
20.5%
9.58%
LOD
Genomic Position (Mb)
B
II:5633032
V:11085892
Phenotype
CB4856
N2
Genotype at QTL peak marker

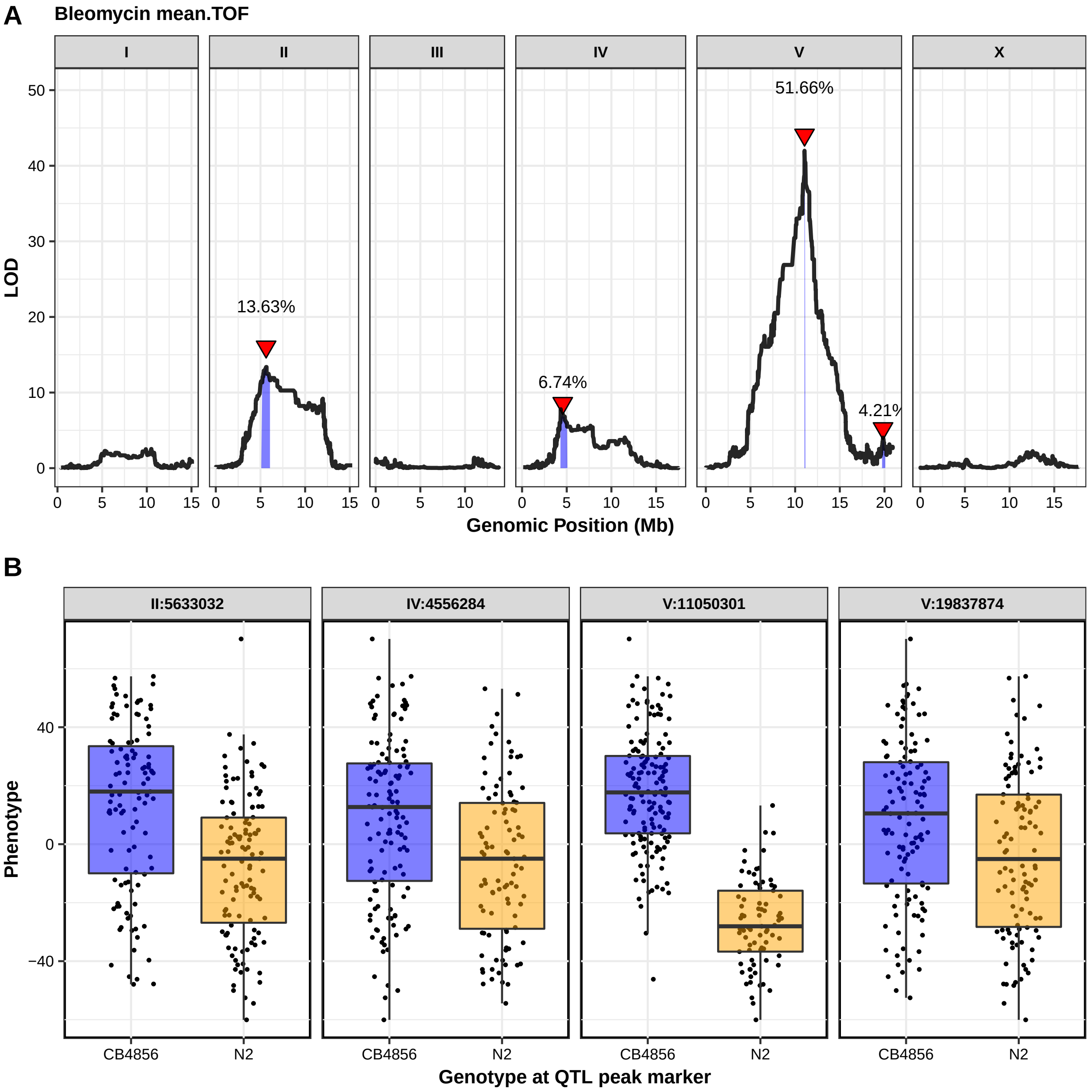
A
Bleomycin mean.TOF
I
II
III
IV
V
X
LOD
13.63%
6.74%
51.66%
4.21%
Genomic Position (Mb)
B
II:5633032
IV:4556284
V:11050301
V:19837874
Phenotype
CB4856
N2
Genotype at QTL peak marker

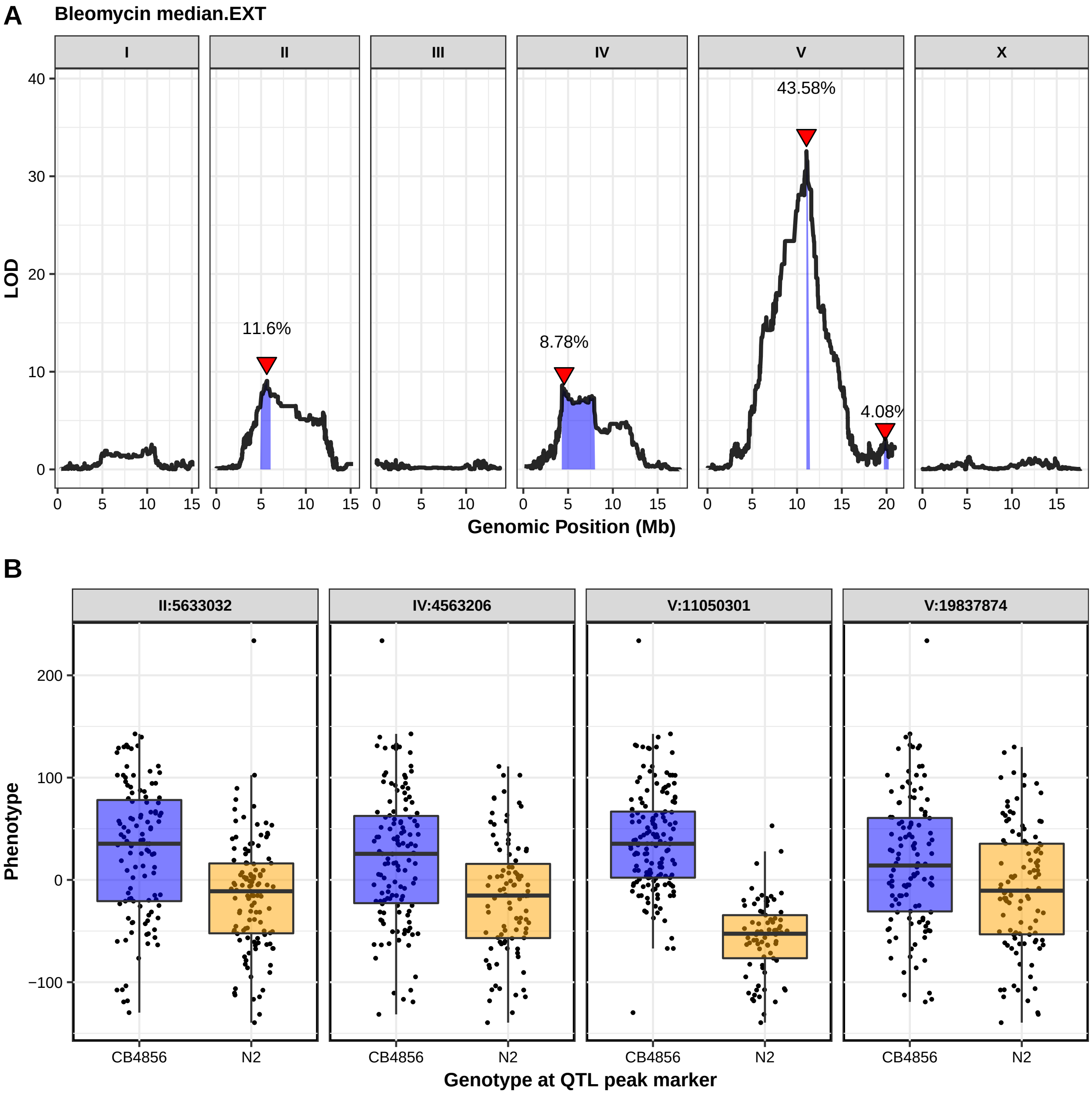
A
Bleomycin median.EXT
I
II
III
IV
V
X
LOD
11.6%
8.78%
43.58%
4.08%
Genomic Position (Mb)
B
II:5633032
IV:4563206
V:11050301
V:19837874
Phenotype
CB4856
N2
Genotype at QTL peak marker

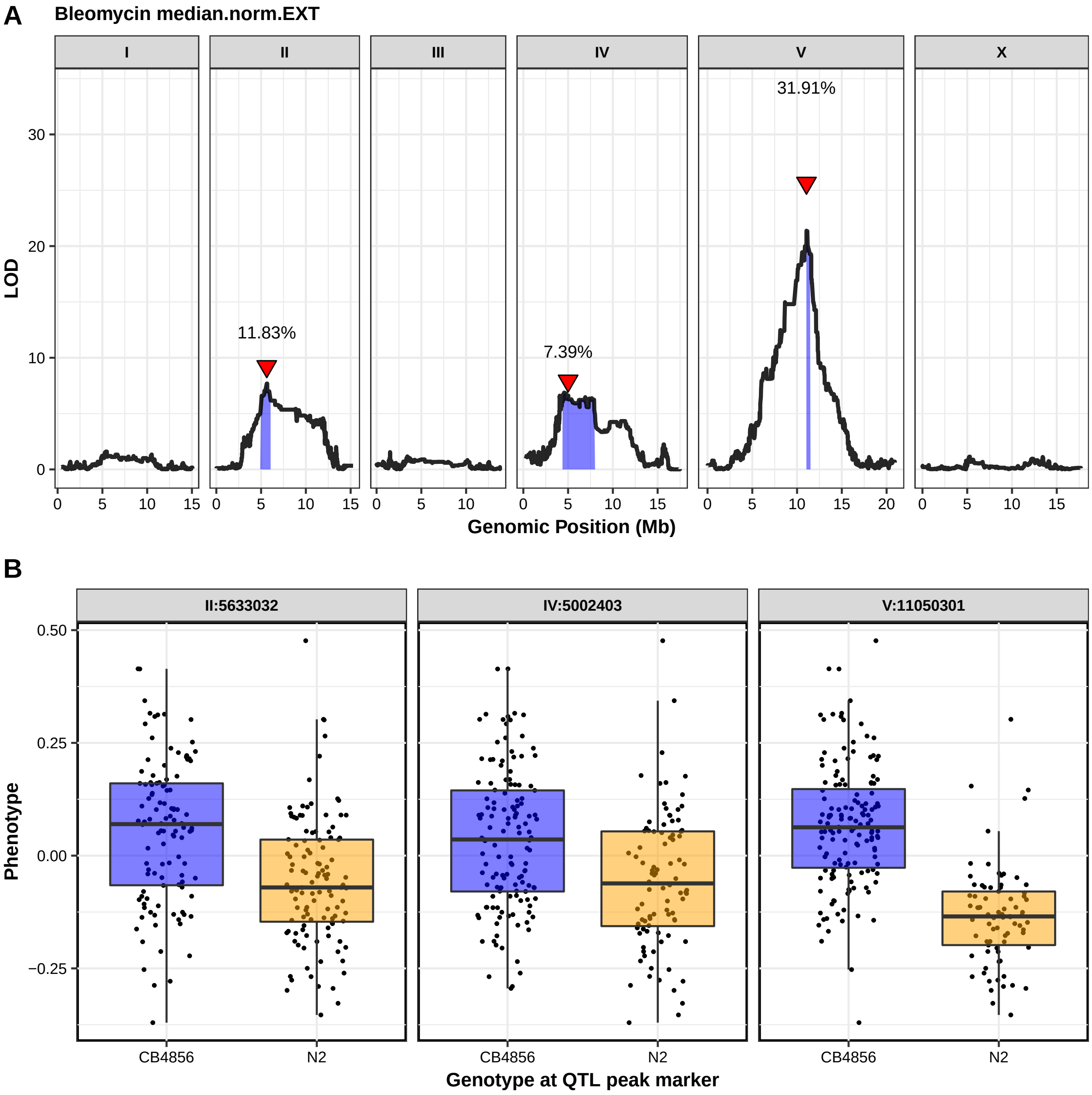
A
Bleomycin median.norm.EXT
I
II
III
IV
V
X
LOD
11.83%
7.39%
31.91%
Genomic Position (Mb)
B
II:5633032
IV:5002403
V:11050301
Phenotype
CB4856
N2
Genotype at QTL peak marker

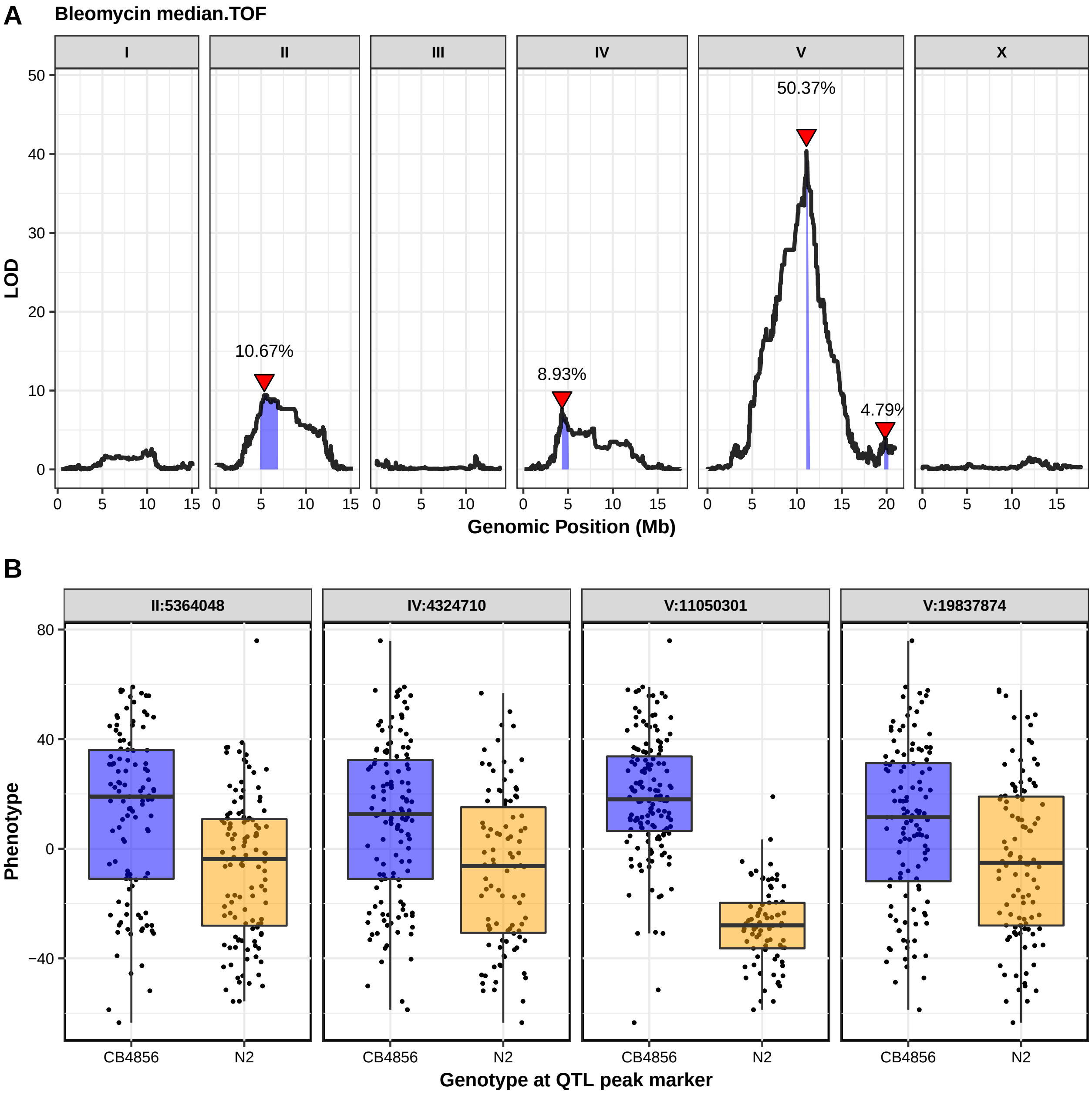
A
Bleomycin median.TOF
I
II
III
IV
V
X
50.37%
10.67%
8.93%
4.79%
LOD
Genomic Position (Mb)
B
II:5364048
IV:4324710
V:11050301
V:19837874
Phenotype
CB4856
N2
Genotype at QTL peak marker

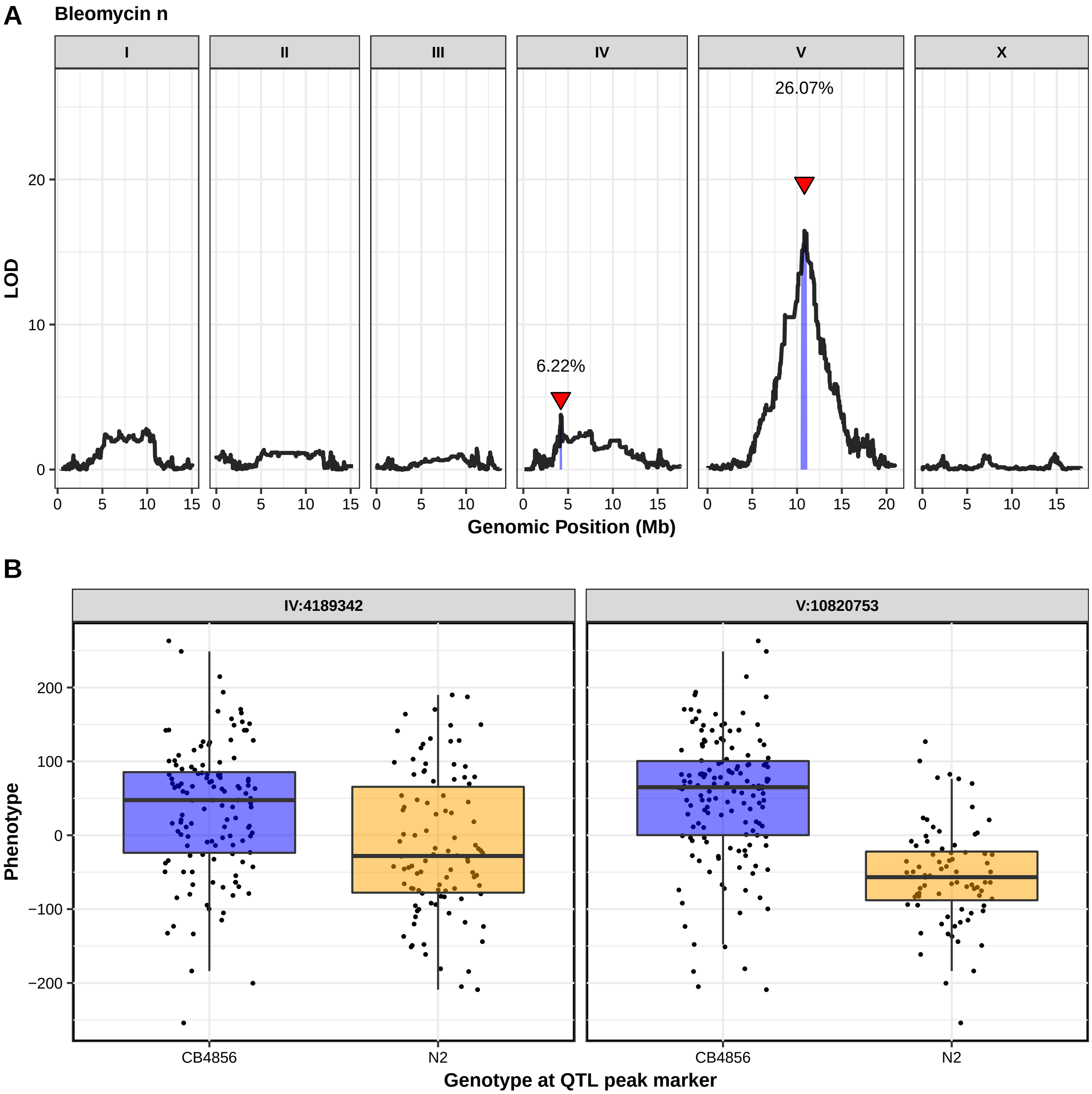
A
Bleomycin n
I
II
III
IV
V
X
26.07%
6.22%
LOD
20
10
0
0
5
10
15
20
Genomic Position (Mb)
B
IV:4189342
V:10820753
Phenotype
200
100
0
−100
−200
CB4856
N2
Genotype at QTL peak marker

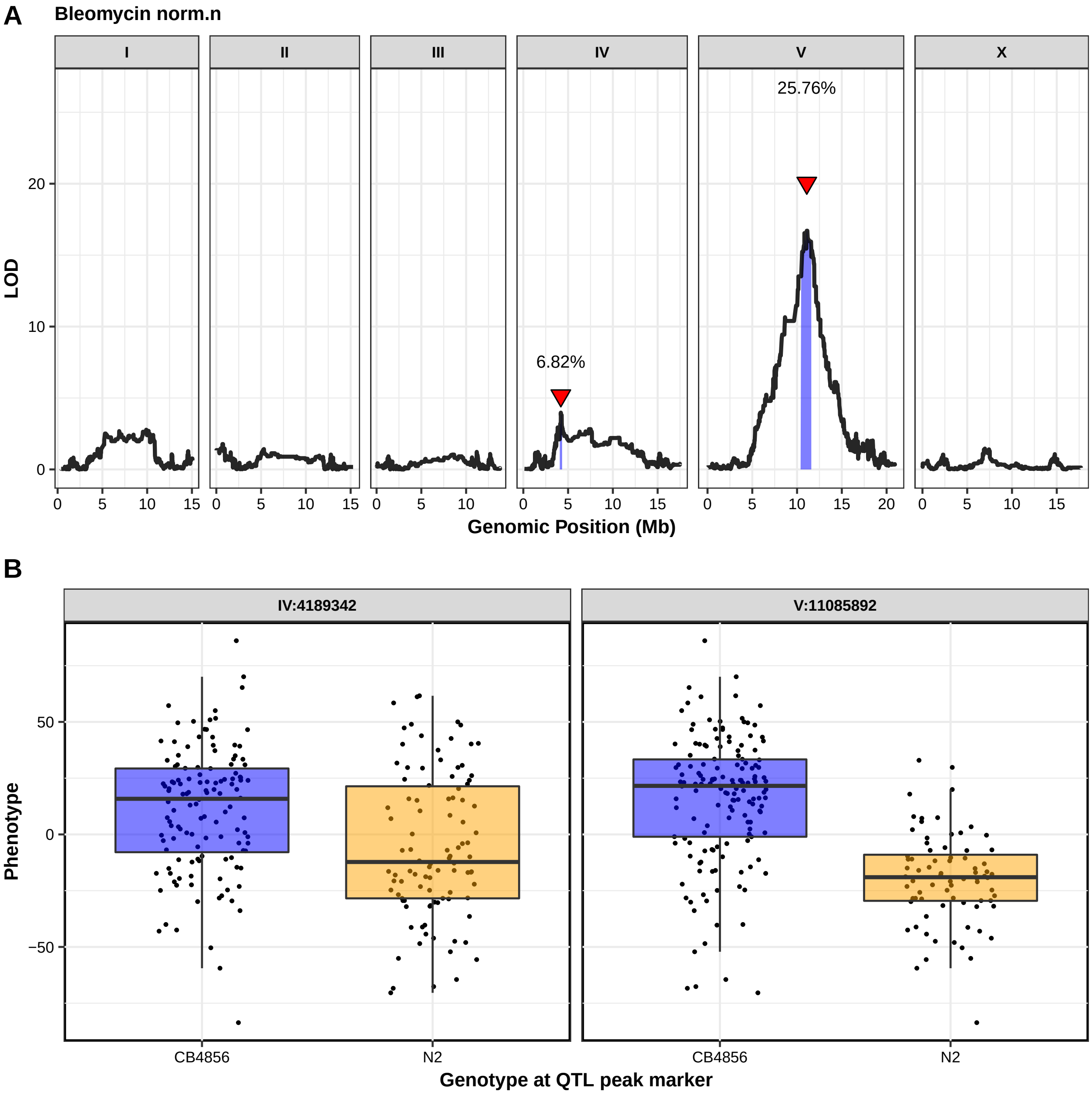
A
Bleomycin norm.n
I
II
III
IV
V
X
25.76%
6.82%
LOD
20
10
0
0 5 10 15
0 5 10 15
0 5 10
0 5 10 15
0 5 10 15 20
0 5 10 15
Genomic Position (Mb)
B
IV:4189342
V:11085892
Phenotype
50
0
−50
CB4856
N2
CB4856
N2
Genotype at QTL peak marker

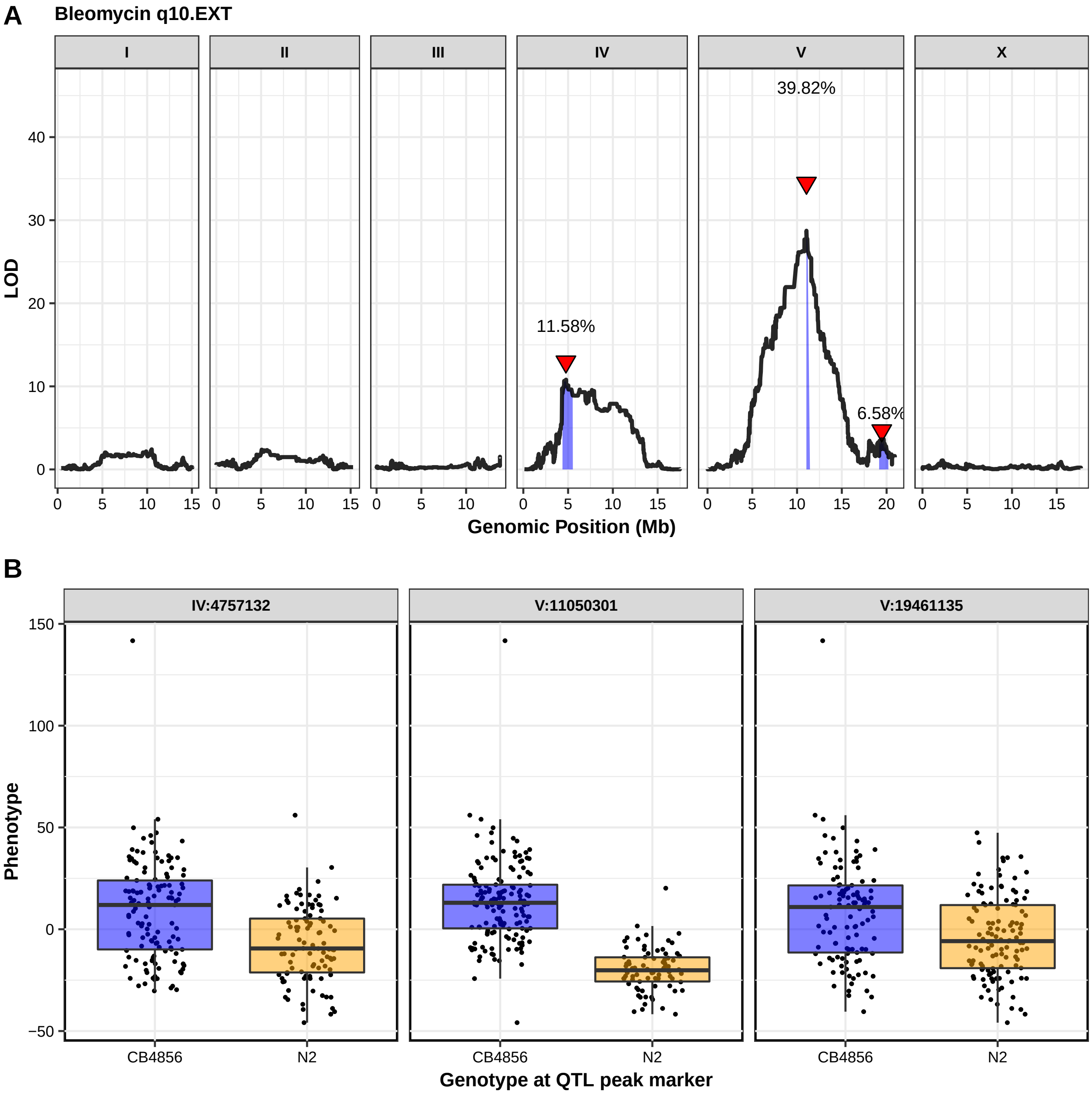
A
Bleomycin q10.EXT
I
II
III
IV
V
X
39.82%
11.58%
6.58%
LOD
Genomic Position (Mb)
B
IV:4757132
V:11050301
V:19461135
Phenotype
CB4856
N2
Genotype at QTL peak marker

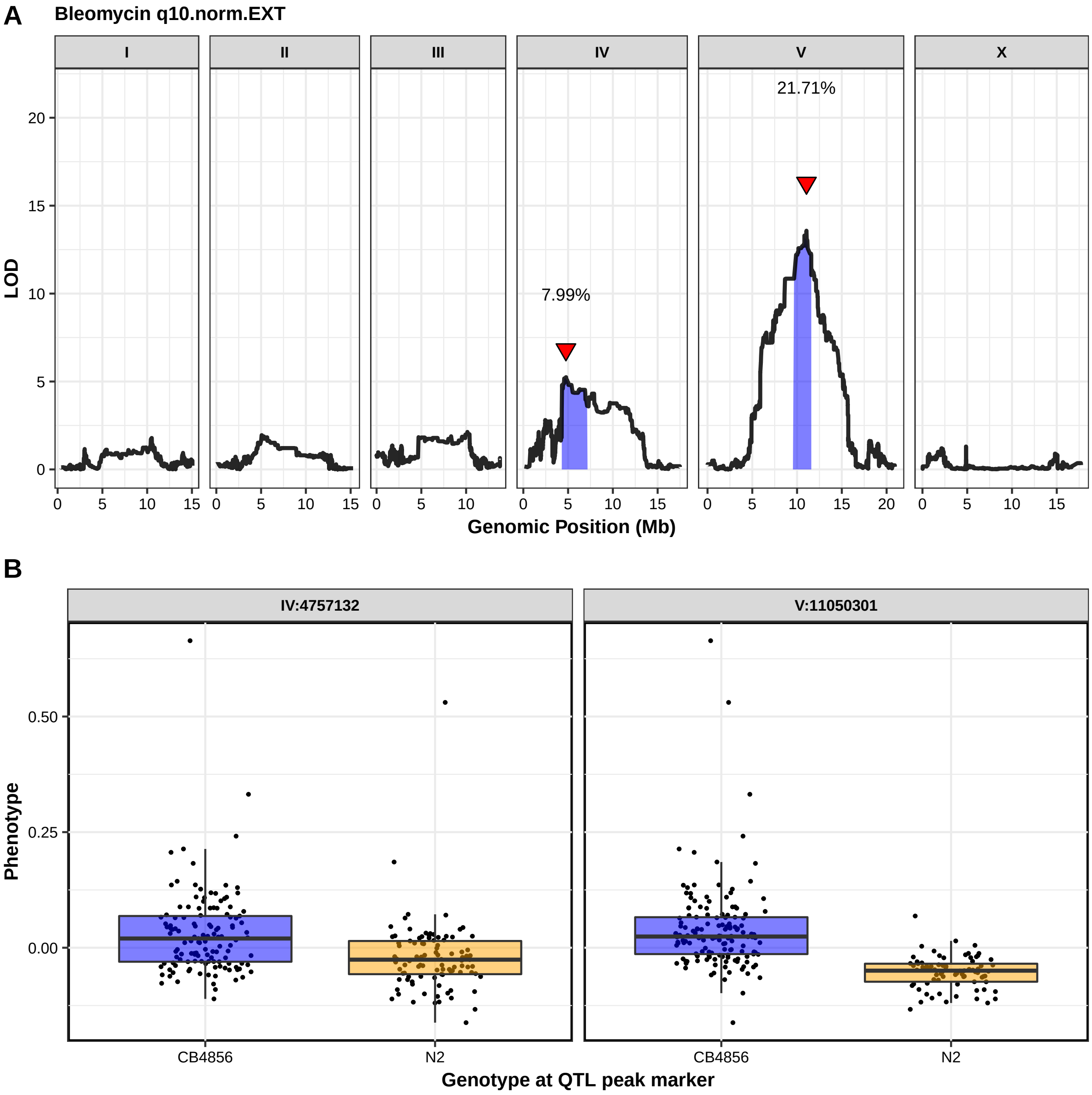
A
Bleomycin q10.norm.EXT
I
II
III
IV
V
X
21.71%
7.99%
LOD
Genomic Position (Mb)
B
IV:4757132
V:11050301
Phenotype
CB4856
N2
Genotype at QTL peak marker

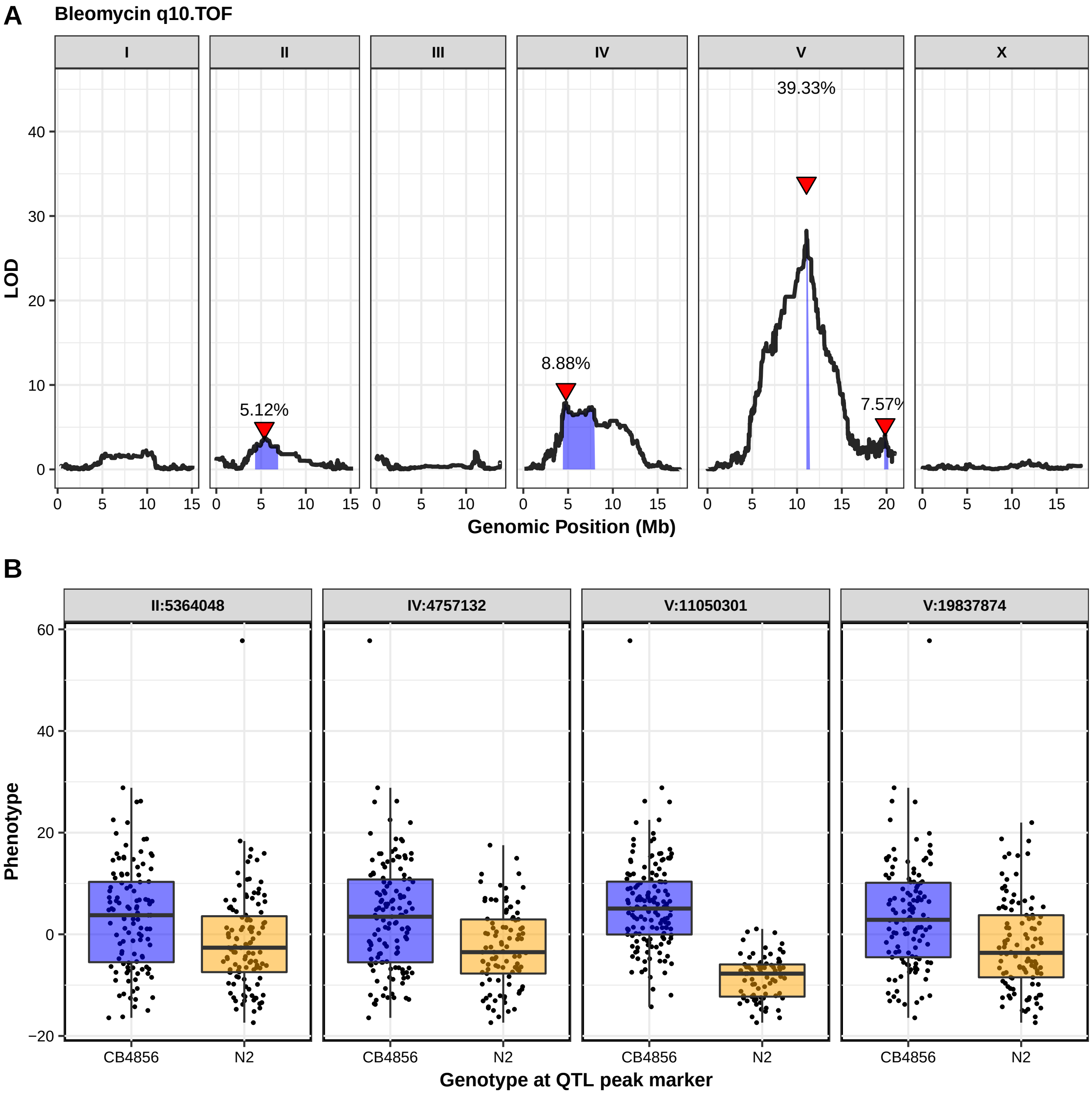
A
Bleomycin q10.TOF
I
II
III
IV
V
X
39.33%
5.12%
8.88%
7.57%
LOD
Genomic Position (Mb)
B
II:5364048
IV:4757132
V:11050301
V:19837874
Phenotype
CB4856
N2
Genotype at QTL peak marker

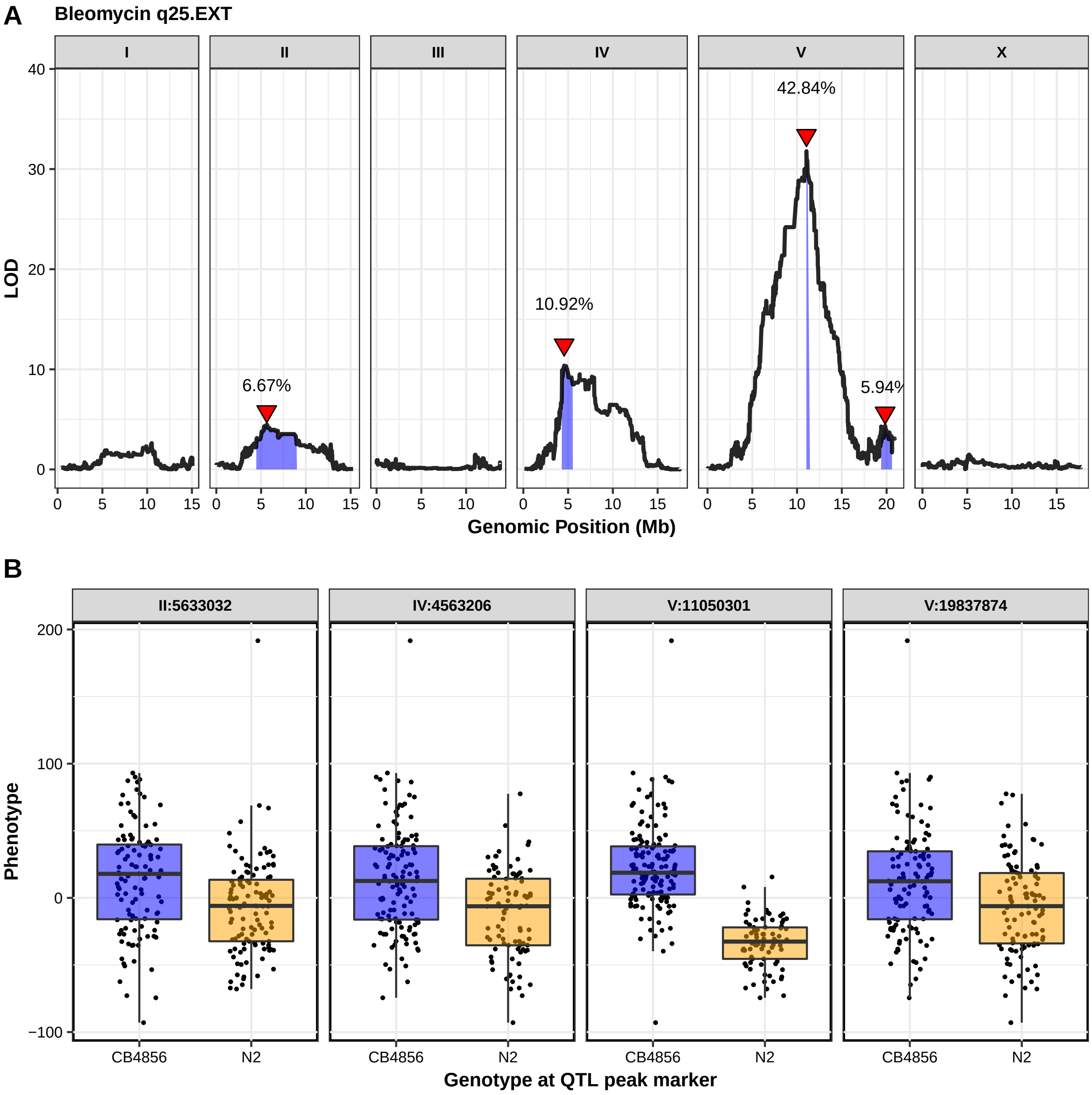

A
Bleomycin q25.EXT
I
II
III
IV
V
X
LOD
Genomic Position (Mb)
6.67%
10.92%
42.84%
5.94%
B
II:5633032
IV:4563206
V:11050301
V:19837874
Phenotype
CB4856
N2
Genotype at QTL peak marker

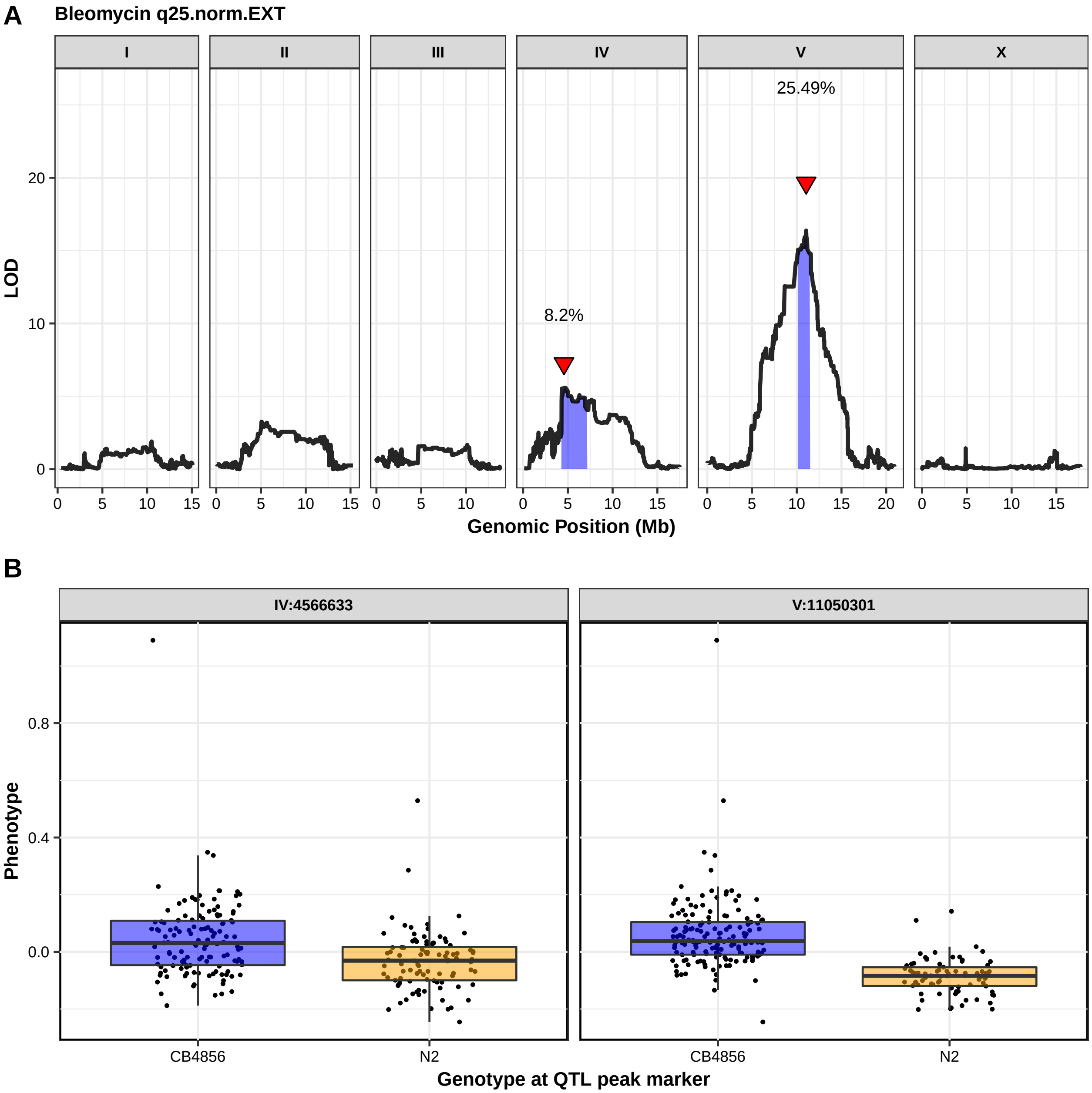
A
Bleomycin q25.norm.EXT
I
II
III
IV
V
X
LOD
Genomic Position (Mb)
8.2%
25.49%
B
IV:4566633
V:11050301
Phenotype
CB4856
N2
Genotype at QTL peak marker

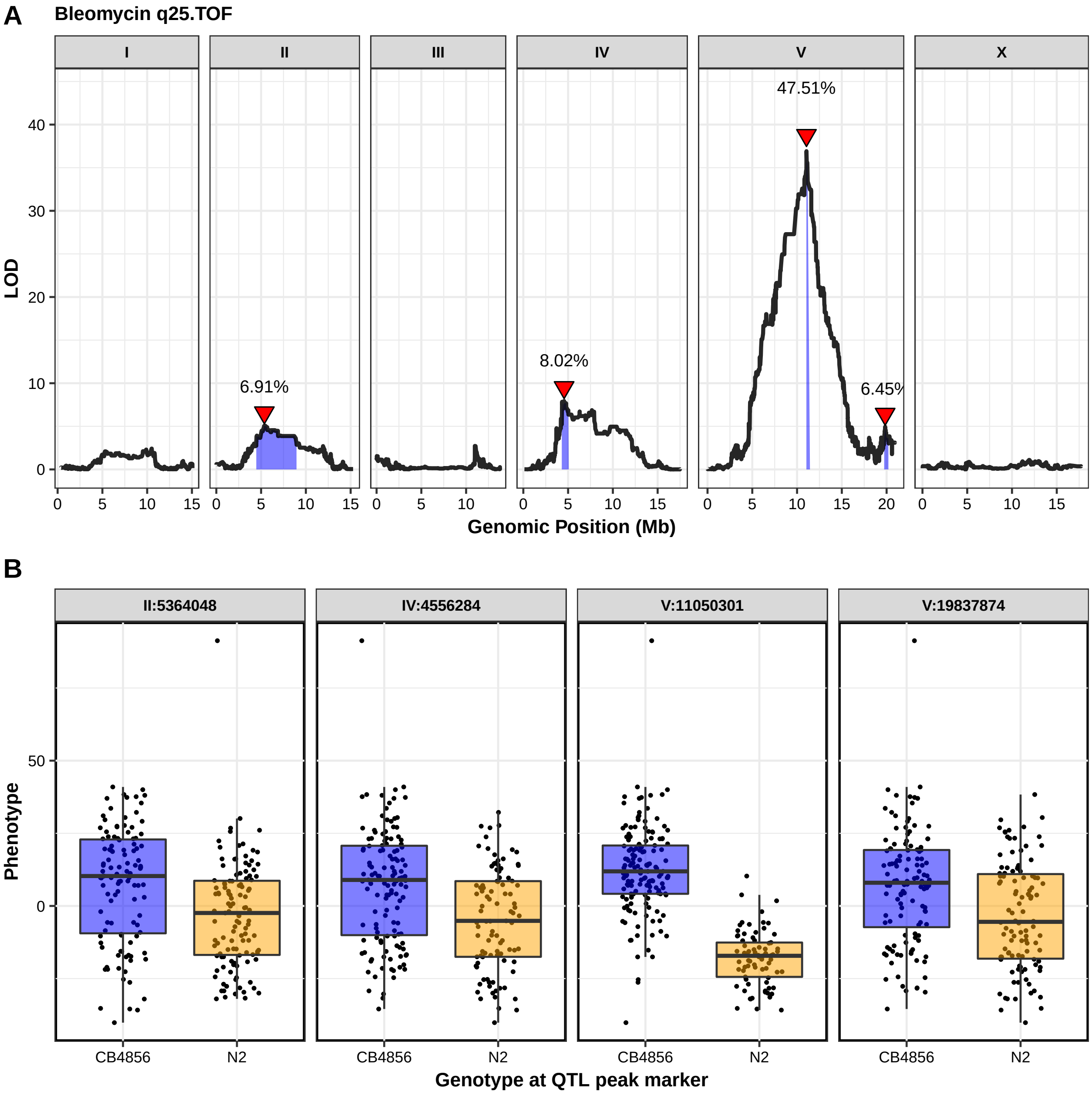
A
Bleomycin q25.TOF
I
II
III
IV
V
X
47.51%
6.91%
8.02%
6.45%
LOD
Genomic Position (Mb)
B
II:5364048
IV:4556284
V:11050301
V:19837874
Phenotype
CB4856
N2
Genotype at QTL peak marker

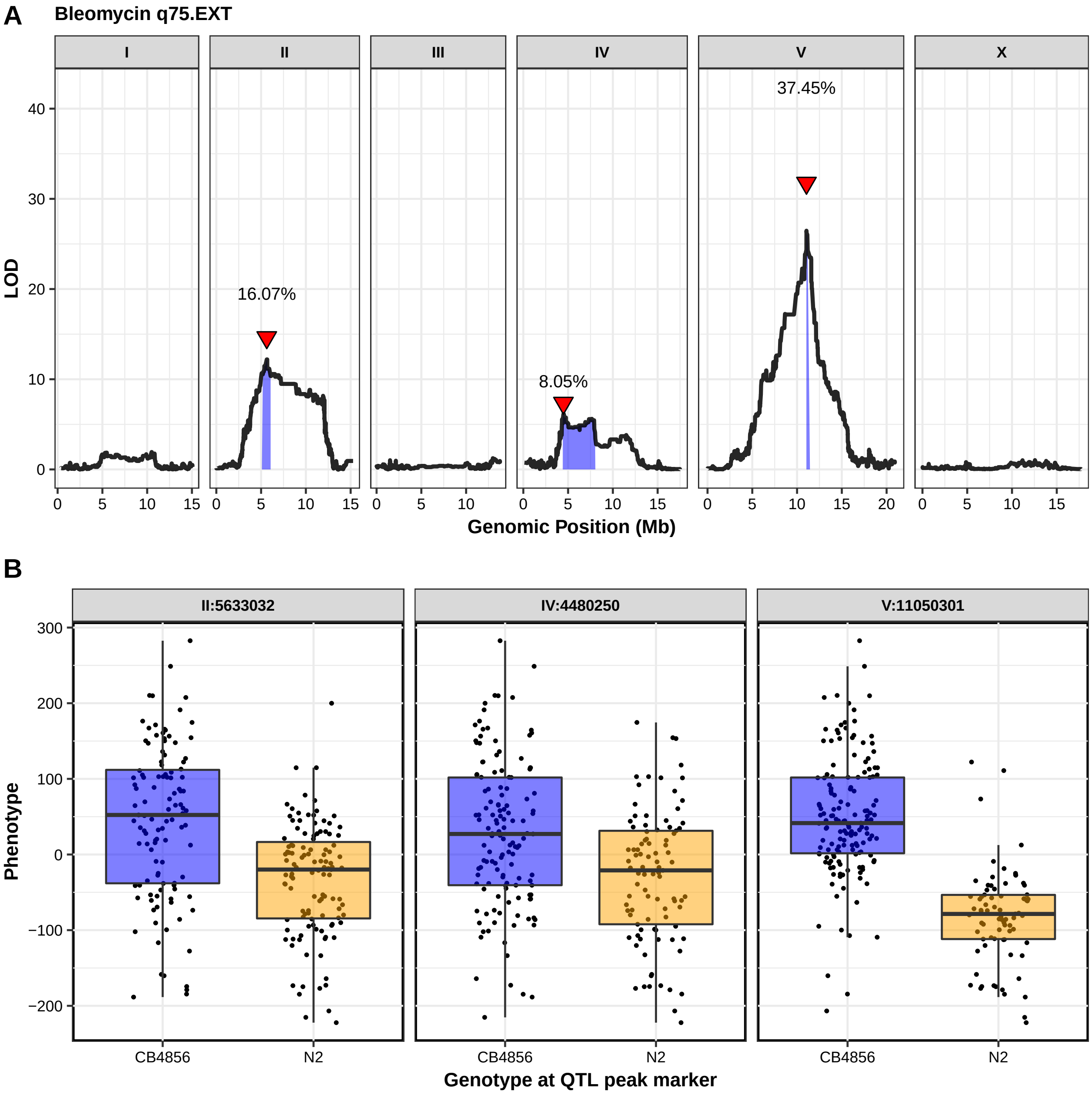
A
Bleomycin q75.EXT
I
II
III
IV
V
X
37.45%
16.07%
8.05%
LOD
Genomic Position (Mb)
B
II:5633032
IV:4480250
V:11050301
Phenotype
CB4856
N2
Genotype at QTL peak marker

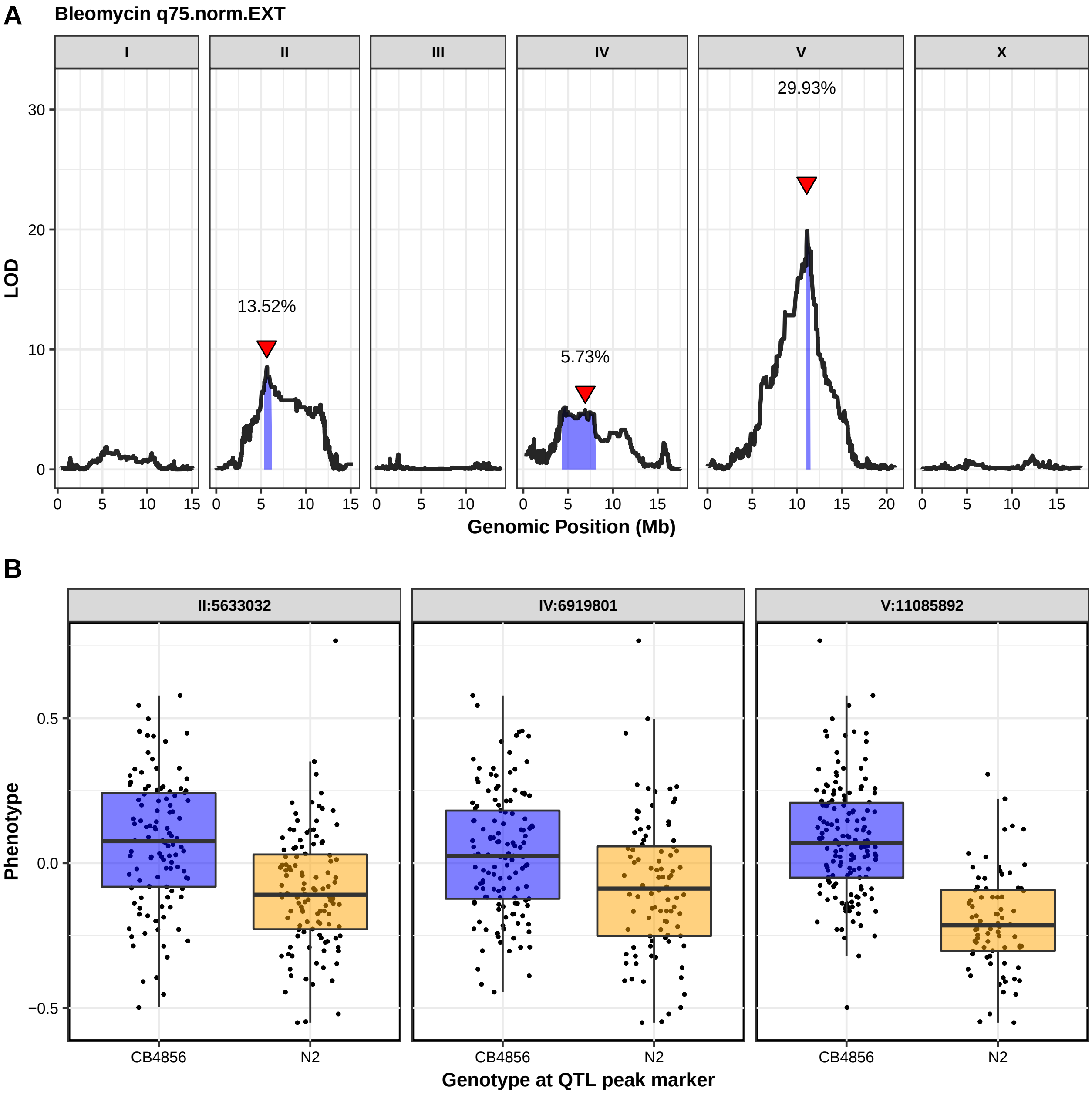
A
Bleomycin q75.norm.EXT
I
II
III
IV
V
X
LOD
30
20
10
0
13.52%
5.73%
29.93%
0
5
10
15
20
Genomic Position (Mb)
B
II:5633032
IV:6919801
V:11085892
Phenotype
0.5
0.0
−0.5
CB4856
N2
Genotype at QTL peak marker

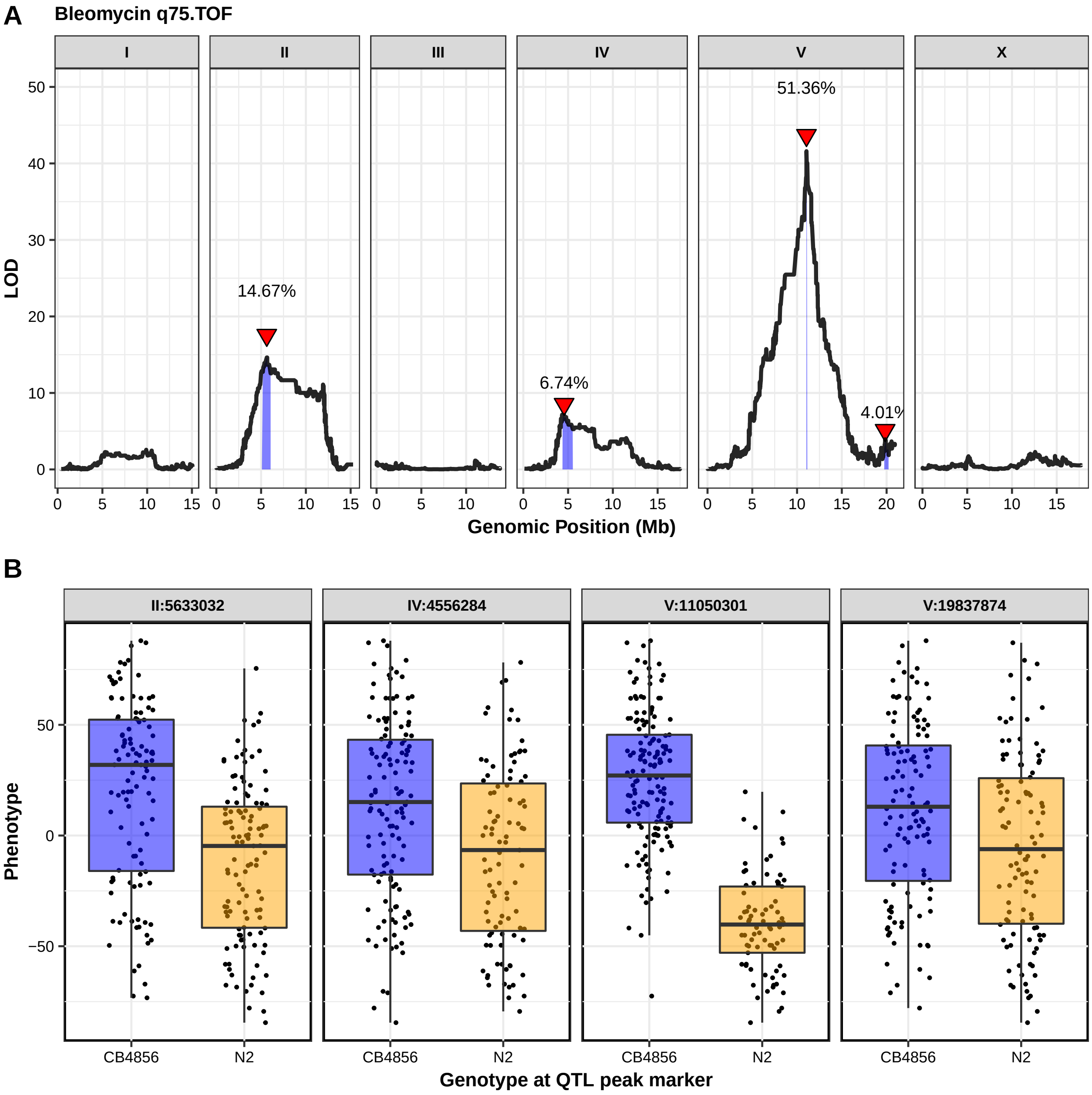
A
Bleomycin q75.TOF
I
II
III
IV
V
X
LOD
Genomic Position (Mb)
14.67%
6.74%
51.36%
4.01%
B
II:5633032
IV:4556284
V:11050301
V:19837874
Phenotype
CB4856
N2
Genotype at QTL peak marker

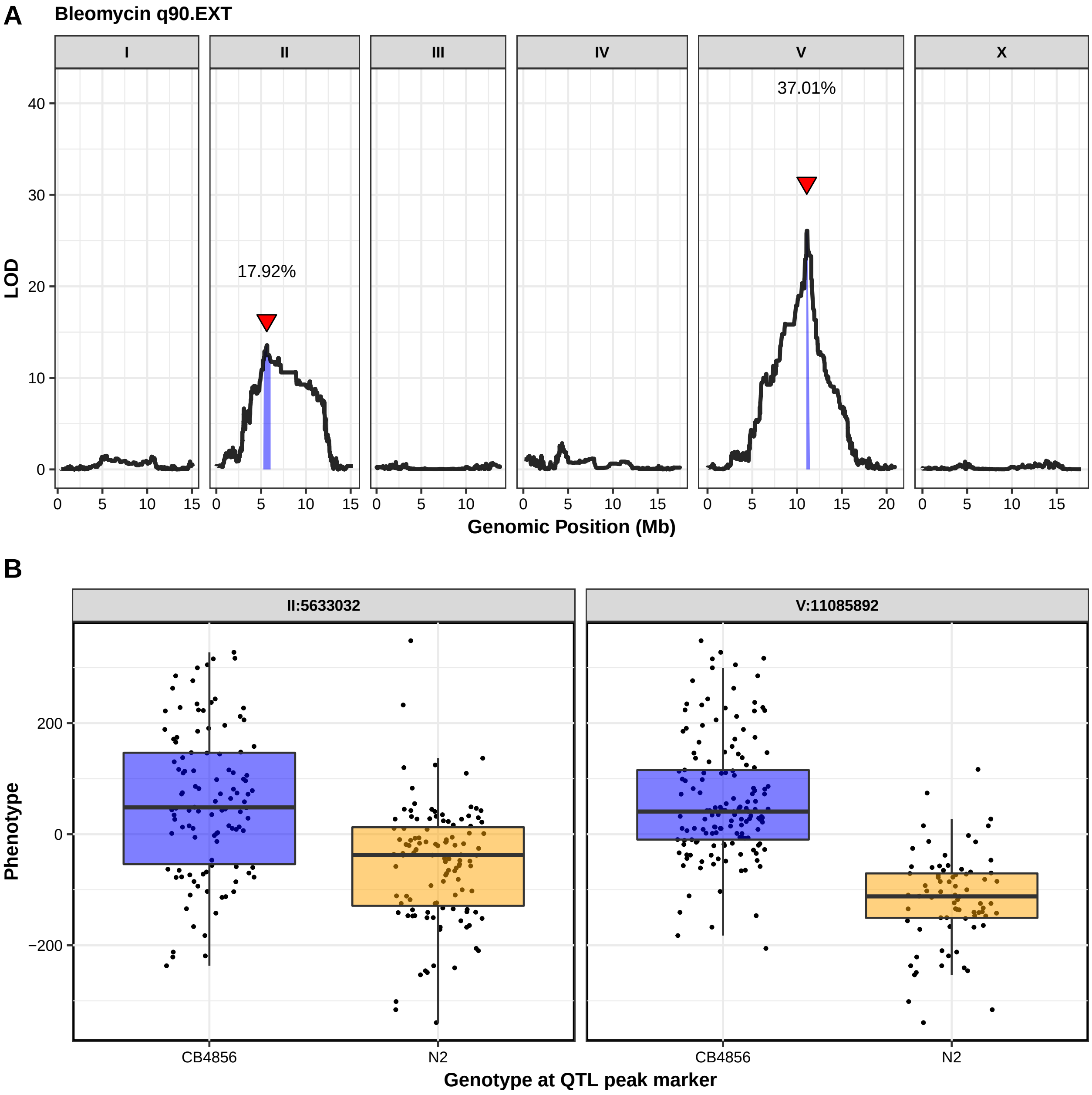
A
Bleomycin q90.EXT
I
II
III
IV
V
X
37.01%
17.92%
LOD
Genomic Position (Mb)
B
II:5633032
V:11085892
Phenotype
CB4856
N2
Genotype at QTL peak marker

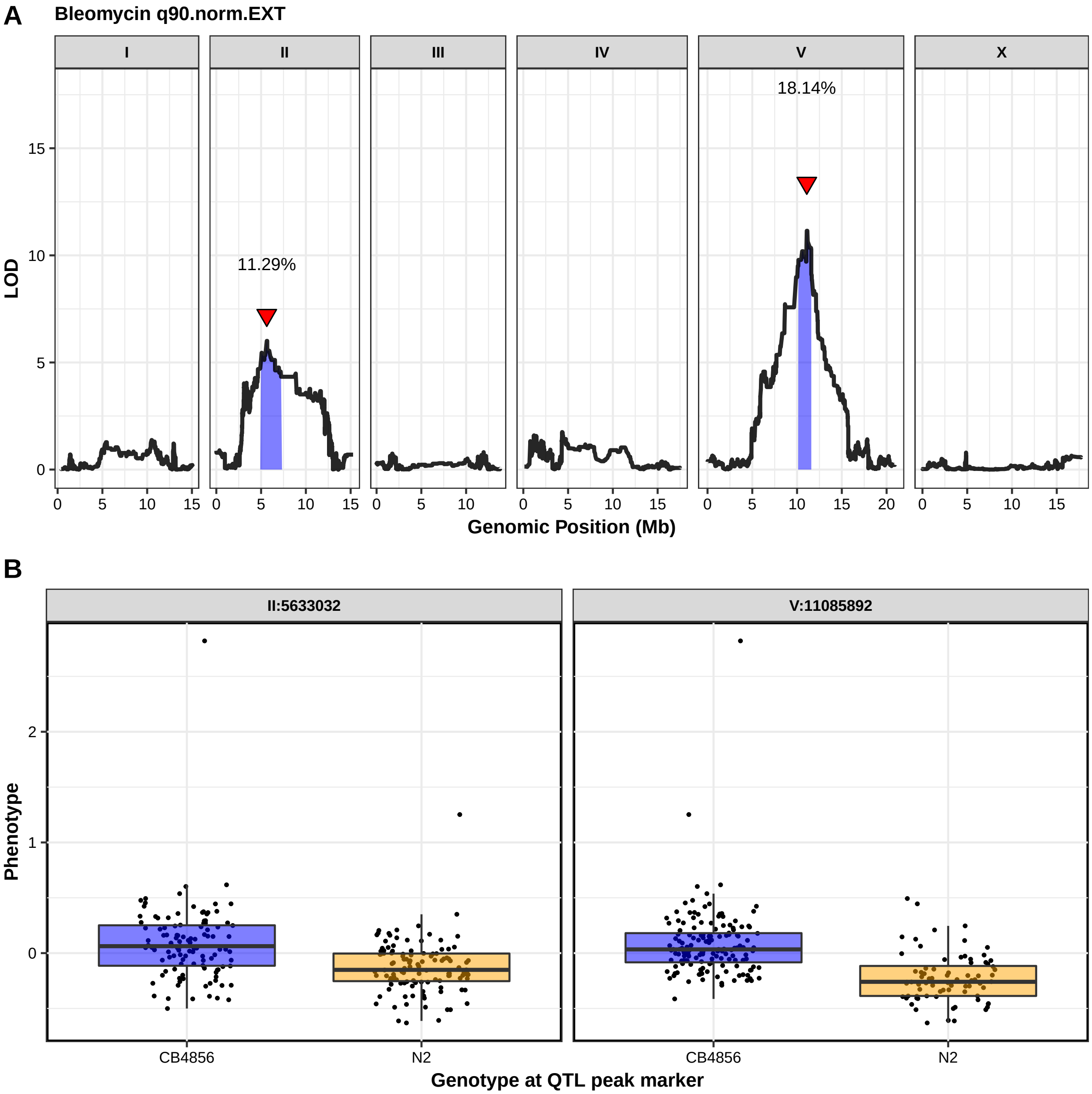
A
Bleomycin q90.norm.EXT
I
II
III
IV
V
X
11.29%
18.14%
LOD
Genomic Position (Mb)
B
II:5633032
V:11085892
Phenotype
CB4856
N2
Genotype at QTL peak marker

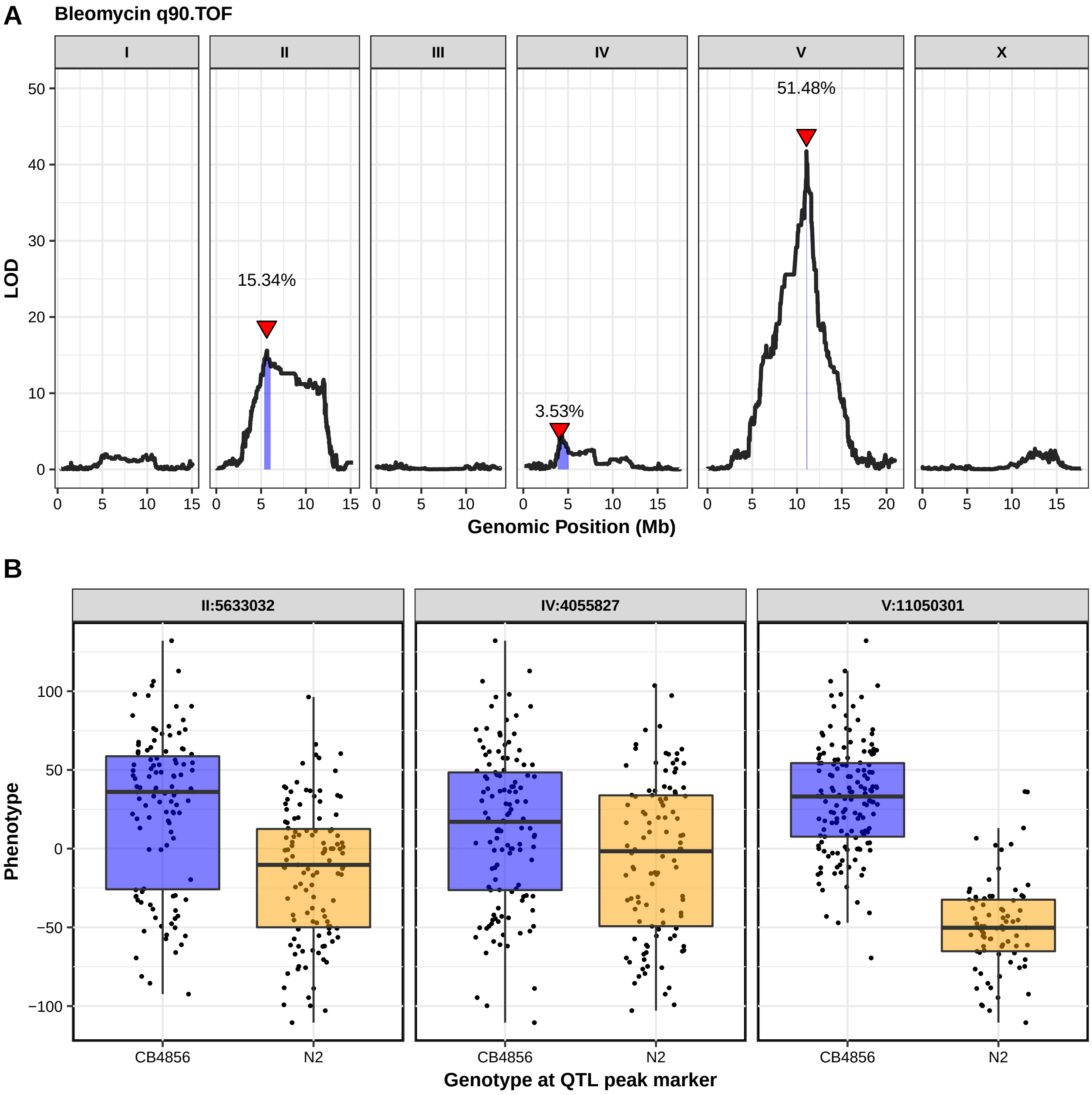
A
Bleomycin q90.TOF
I
II
III
IV
V
X
51.48%
15.34%
3.53%
LOD
Genomic Position (Mb)
B
II:5633032
IV:4055827
V:11050301
Phenotype
CB4856
N2
Genotype at QTL peak marker

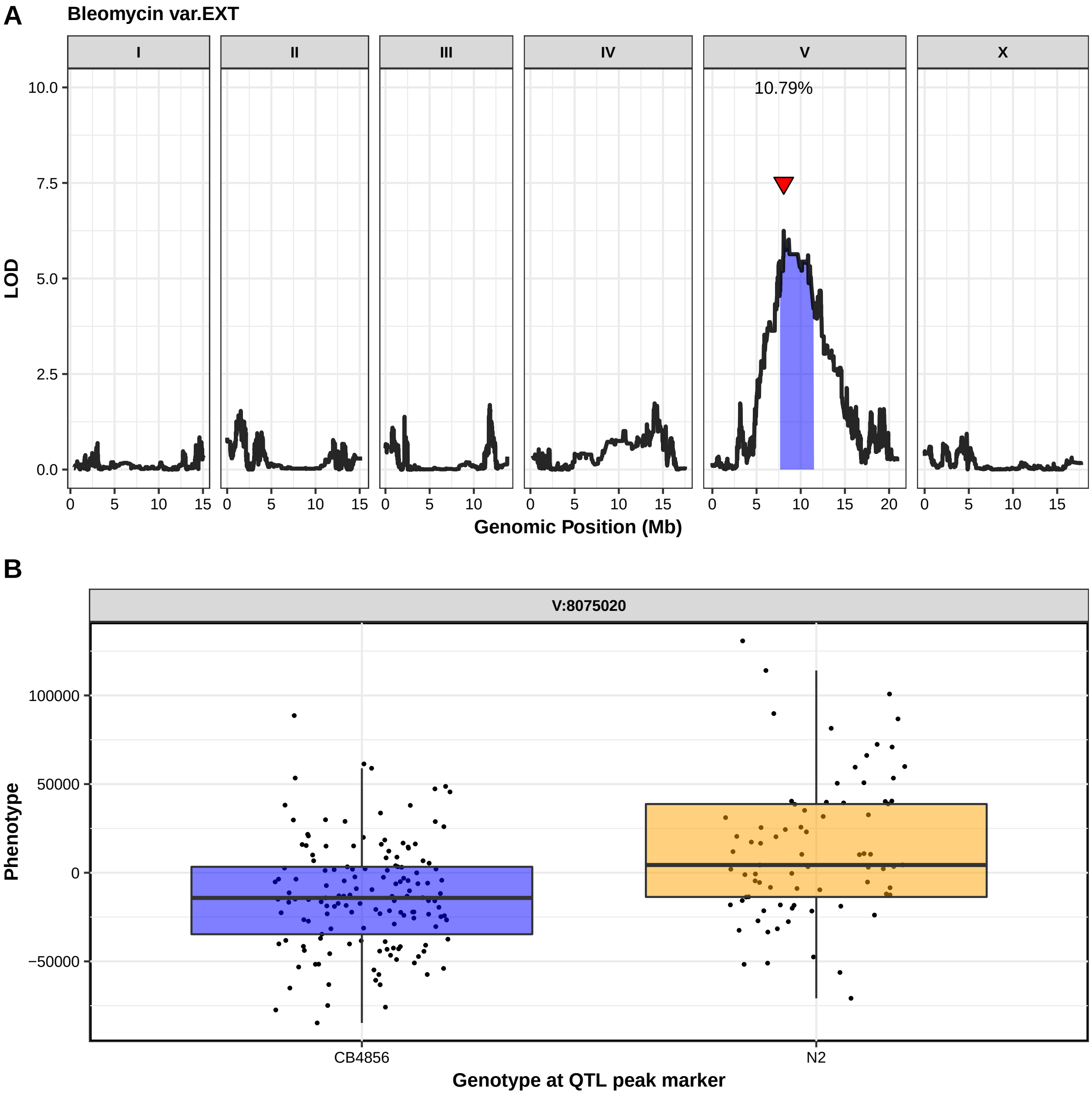
A
Bleomycin var.EXT
I
II
III
IV
V
X
10.79%
LOD
10.0
7.5
5.0
2.5
0.0
0
5
10
15
20
Genomic Position (Mb)
B
V:8075020
Phenotype
100000
50000
0
−50000
CB4856
N2
Genotype at QTL peak marker

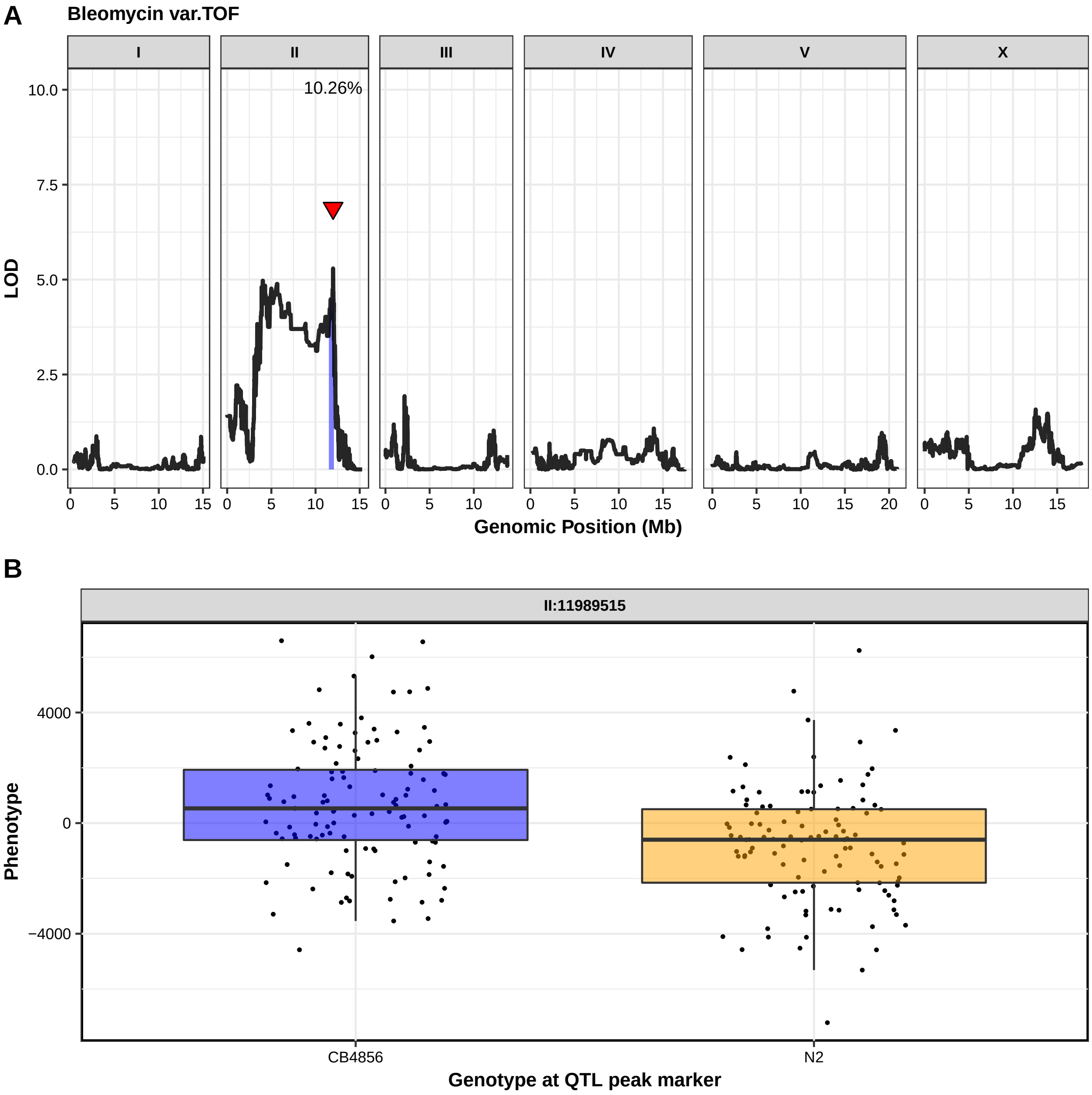
A
Bleomycin var.TOF
I
II
III
IV
V
X
10.26%
LOD
10.0
7.5
5.0
2.5
0.0
Genomic Position (Mb)
B
II:11989515
Phenotype
4000
0
−4000
CB4856
N2
Genotype at QTL peak marker

**Figure S5**

Linkage-mapping analysis of bleomycin-response variation for all 26 high-throughput traits are shown. (**A**) On the x-axis, each of 13,001 genomic markers, split by chromosome, were tested for correlation with phenotypic variation across the RIAIL panel. The log of the odds (LOD) score for each marker is reported on the y-axis. Each significant quantitative trait locus (QTL) is indicated by a red triangle at the peak marker, and a blue ribbon shows the 95% confidence interval around the peak marker. The total amount of phenotypic variance across the RIAIL panel explained by the genotype at each peak marker is shown as a percentage. (**B**) RIAIL phenotypes (y-axis), split by allele at each QTL peak marker (x-axis) are reported. For each significant QTL, phenotypes of RIAILs containing the N2 allele (orange) are compared to phenotypes of RIAILs containing the CB4856 allele (blue). Phenotypes are shown as Tukey box plots, and each point is the phenotype of an individual strain.

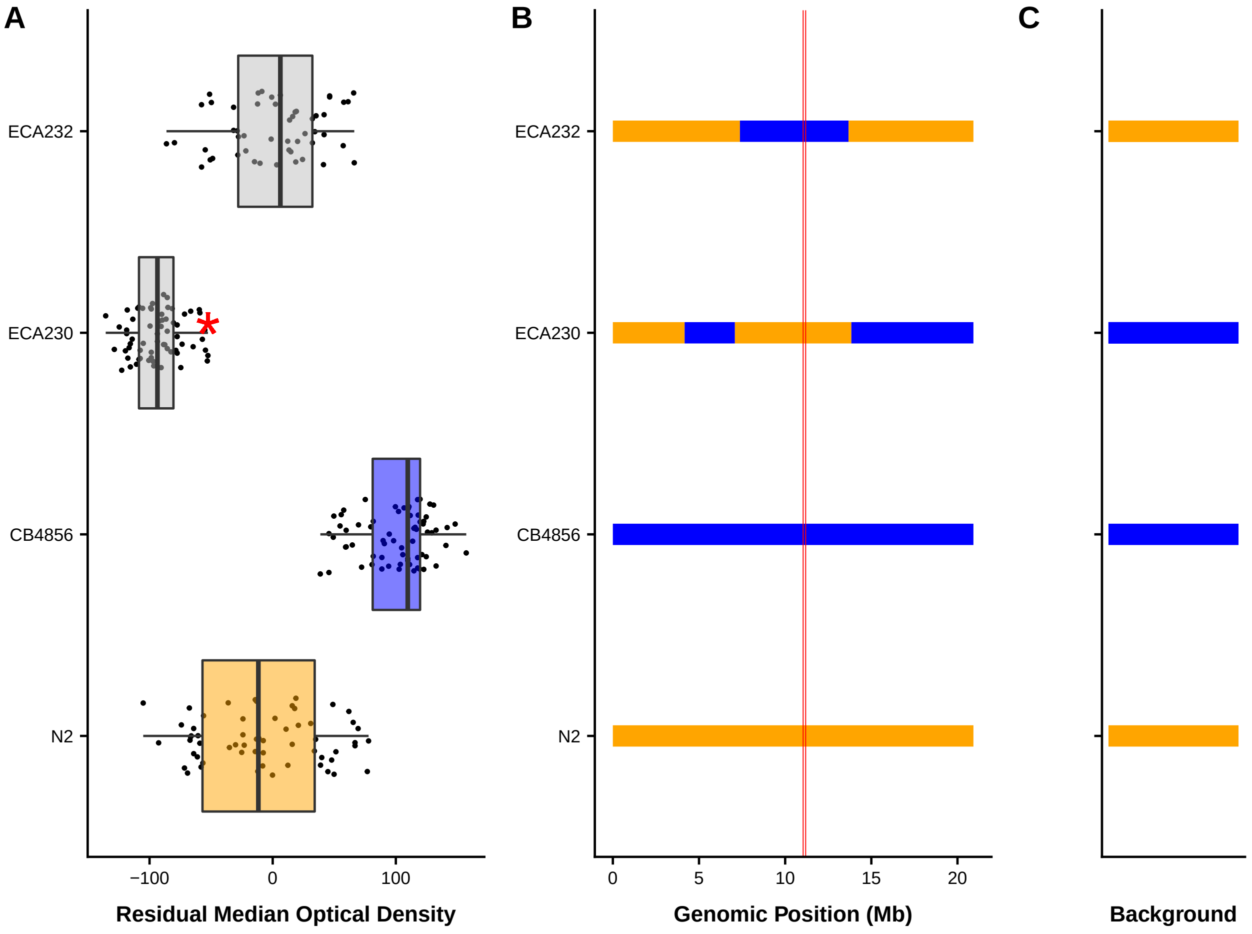


**Figure S6**

Phenotypes and genotypes of NIL strains are shown. (**A**) Phenotypes for each strain are shown as Tukey box plots, with strain name on the y-axis and residual bleomycin median optical density on the x-axis. Each point is a biological replicate. Parental strain box plots are colored by their genetic background, with orange indicating an N2 background and blue indicating a CB4856 genetic background. NILs are shown as grey box plots. A red asterisk indicates a significant difference between the phenotype of a given strain and the phenotype of the corresponding parental strain ($p < 0.05$, Tukey HSD). (**B**) Chromosomal representations of chromosome V are shown for each of the strains in **A**. Strain names are reported on the y-axis, and genomic position (Mb) is shown on the x-axis. Blocks of color indicate genotypes of genomic regions with orange indicating the N2 genotype and blue indicating the CB4856 genotype. Vertical red lines mark the confidence interval of the QTL from linkage mapping. (**C**) Background genotypes are represented as rectangles with colors indicating N2 (orange) or CB4856 (blue) genetic backgrounds.

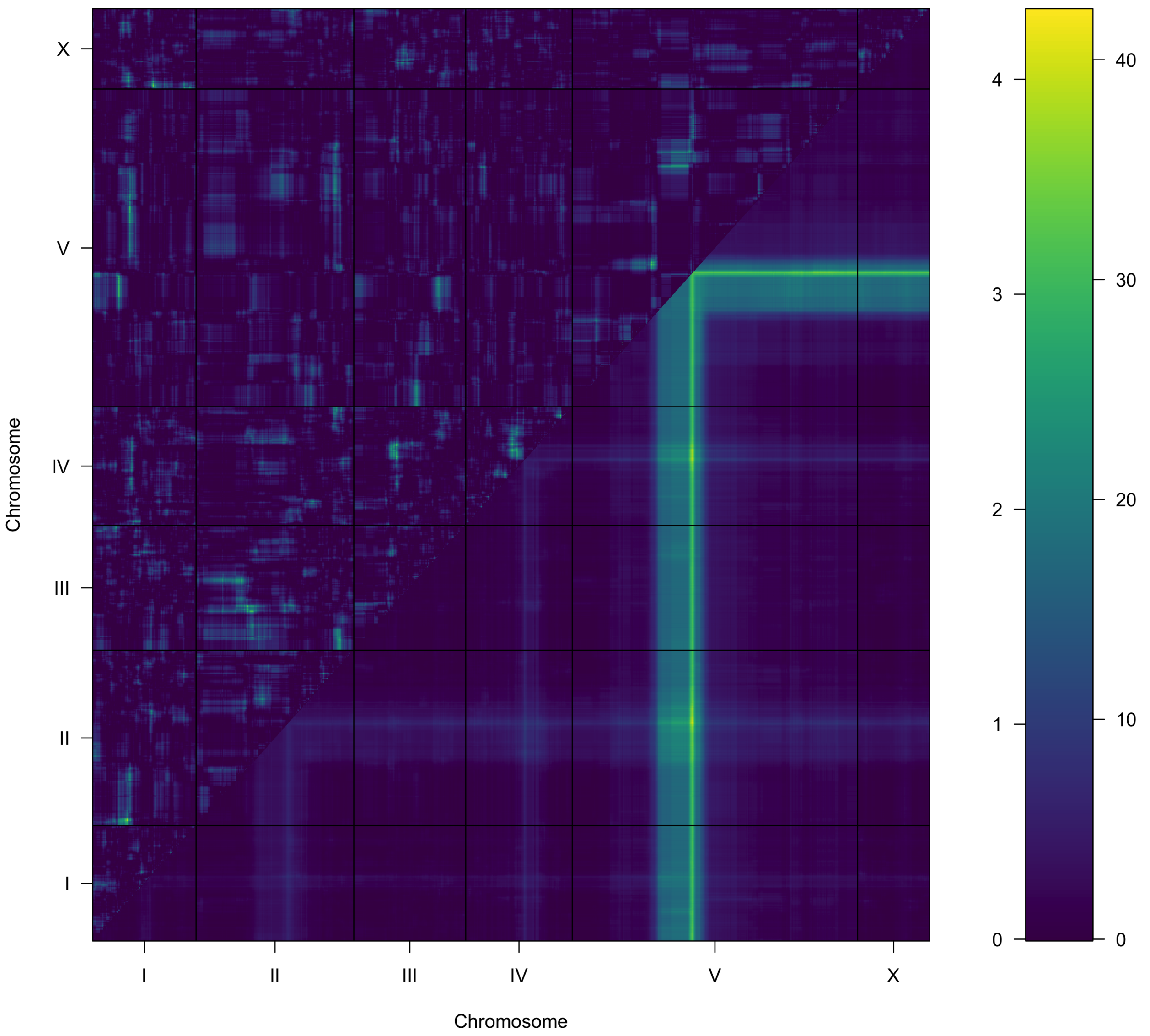


**Figure S7**

A two-factor genome scan for residual optical density in bleomycin is shown. Log of the odds (LOD) scores are shown for each pairwise combination of loci, split by chromosome. The upper-left triangle contains the epistasis LOD scores, and the lower-right triangle contains LOD scores for the full model. LOD scores are shown as colors, with lower scores in blue and higher scores in yellow, as shown in the color scale. The epistatic LOD score axis is on the left of the color scale and the full LOD score axis is on the right of the color scale.

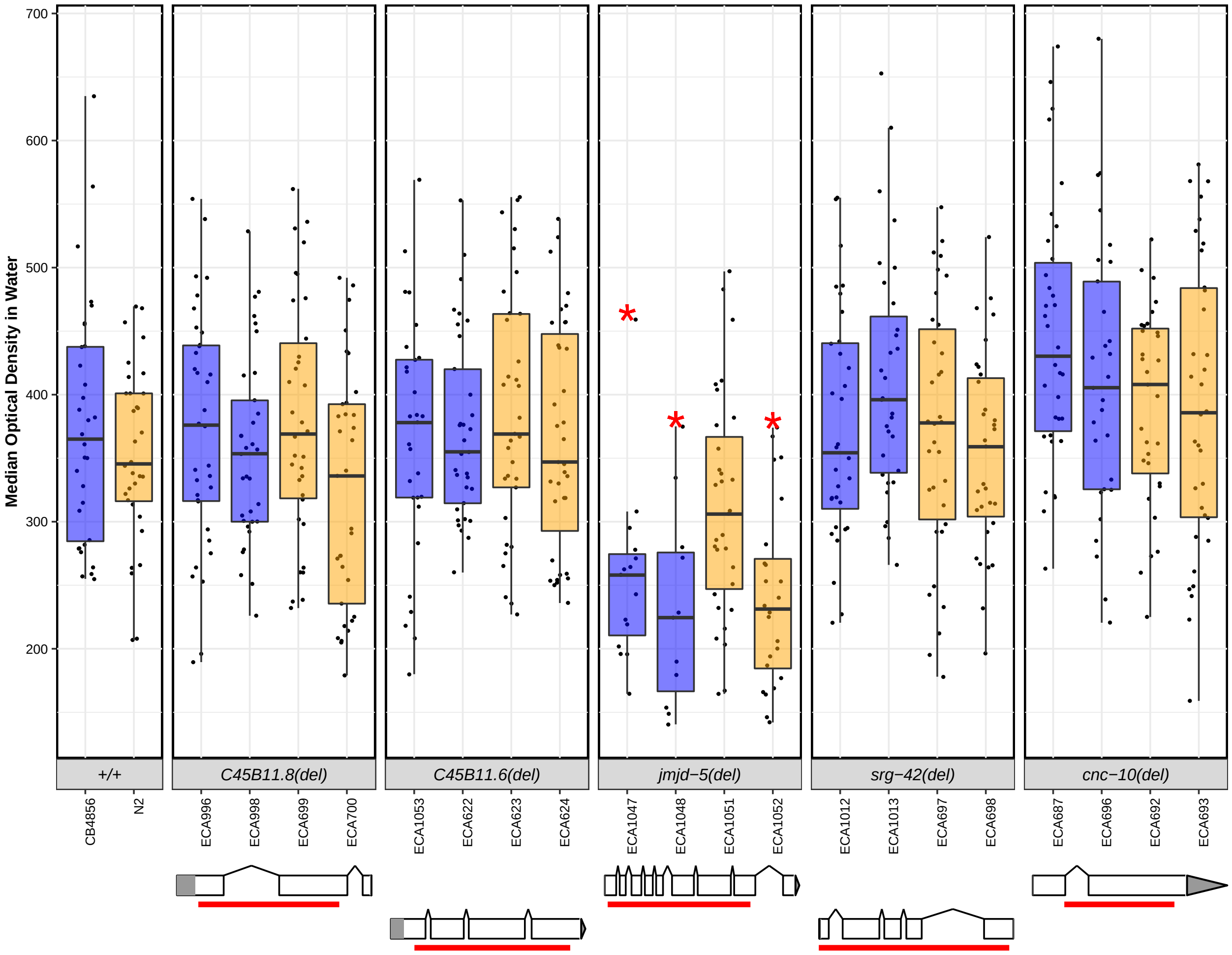


**Figure S8**

Control-condition phenotypes of deletion alleles for each candidate gene are shown. Deletion alleles for each candidate gene were tested in response to the control condition (lysate in K medium with 1% distilled water). Control-condition responses are shown as Tukey box plots, with the strain name on the x-axis, split by gene, and median optical density on the y-axis. Each point is a biological replicate. Strains are colored by their background genotype (orange indicates an N2 genetic background, and blue indicates a CB4856 genetic background). For each gene, two independent deletion alleles in each background were created and tested. Red asterisks indicate a significant difference ($p < 0.05$, Tukey HSD) between a strain with a deletion and the parental strain that has the same genetic background. Depictions of each deletion allele are shown below the phenotype for each candidate gene. White rectangles indicate exons and diagonal lines indicate introns. The 5' and 3' UTRs are shown by grey rectangles and triangles, respectively. The region of the gene that was deleted by CRISPR-Cas9 directed genome editing is shown as a red bar beneath each gene model.

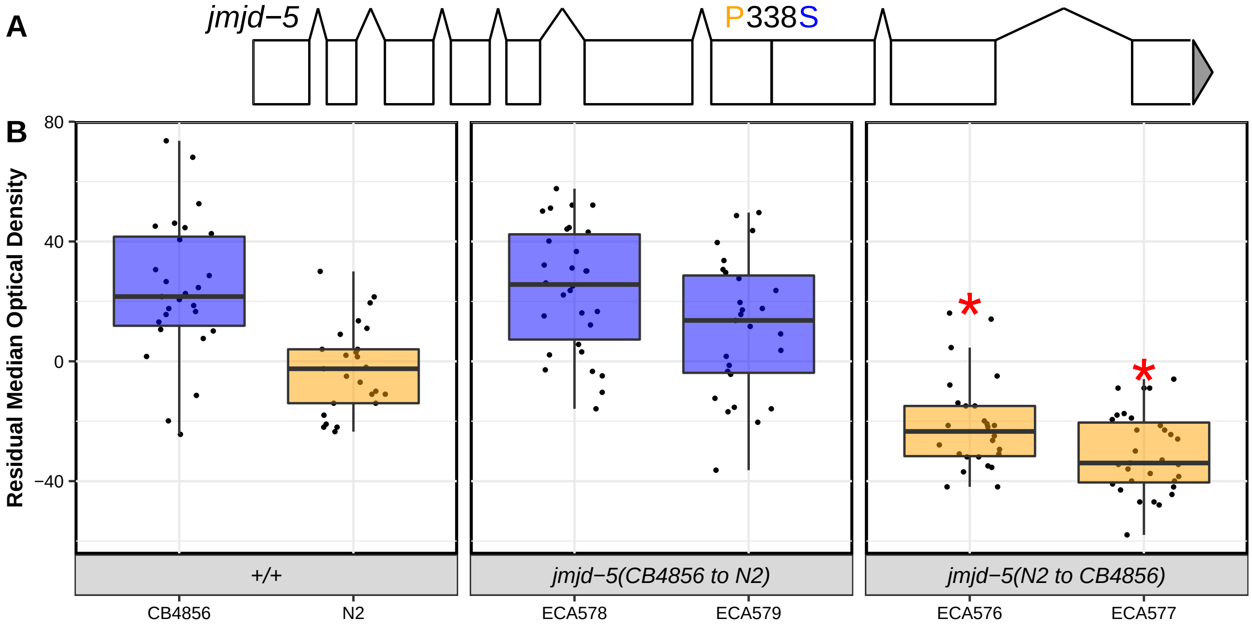


**Figure S9**

Reciprocal allele-replacement strains of *jmjd-5* were tested in response to bleomycin. (**A**) A model of *jmjd-5* is shown with white rectangles indicating exons and black diagonal lines indicating introns. The 5' and 3' UTRs are shown as a grey rectangle and triangle, respectively. The location of the amino-acid variant between the N2 and CB4856 strains is shown in black text above the gene depiction with the N2 residue in orange and the CB4856 residue in blue. (**B**) Bleomycin responses of allele-replacement strains are shown as Tukey box plots, split and colored by genetic background (N2 in orange and CB4856 in blue) with the strain name on the x-axis and residual median optical density on the y-axis. Each point is a biological replicate. Red asterisks indicate a significant difference between an allele-replacement strain and the parental strain sharing its genetic background ($p < 0.05$, Tukey HSD).

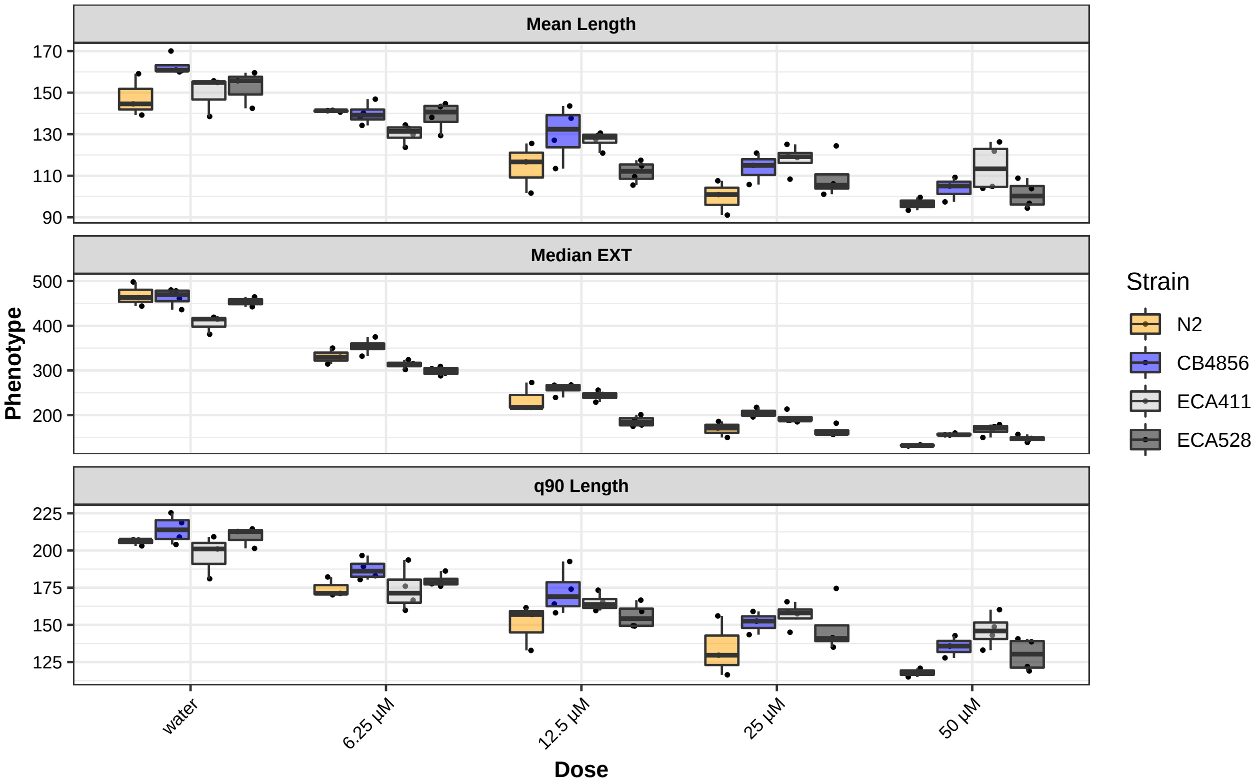


**Figure S10**

Dose-response phenotypes from the modified HTA are shown for three high-throughput fitness traits: mean length (mean.TOF), median optical density (median.EXT), and the 90$^{th}$ quantile of optical density (q90 EXT). Phenotypes for each of the four tested strains are shown as Tukey boxplots, colored by strain (N2 - orange, CB4856 - blue, ECA411 - light grey, ECA528 - dark grey). The x-axis shows the concentration of bleomycin (or water) to which the animals were exposed, and the y-axis shows the pruned phenotype. Each point is a biological replicate.

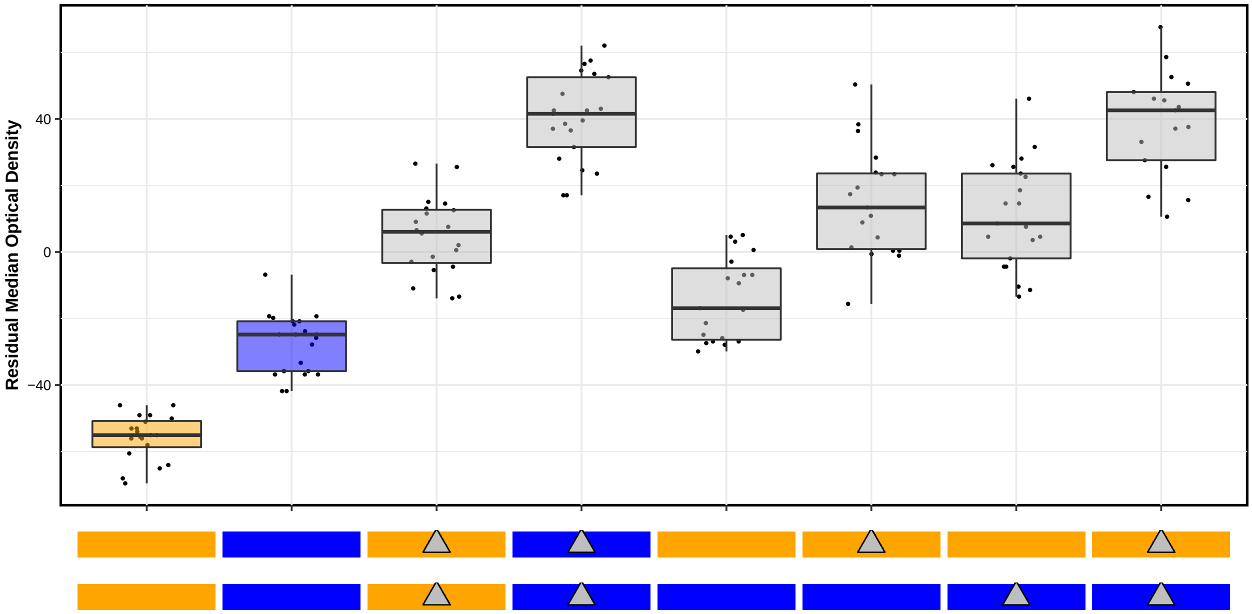


**Figure S11**

Results of the *jmjd-5* reciprocal hemizygosity assay are shown. The y-axis shows the residual median optical density for each strain reported along the x-axis. Bleomycin responses are reported as Tukey box plots where each point is a biological replicate. The genotypes of each strain are shown as colored rectangles beneath each box plot, where each rectangle represents a homolog (orange rectangles are an N2 genotype, and blue rectangles are a CB4856 genotype). The maternal homolog is shown on top and the paternal homolog is shown on bottom. Grey triangles indicate a deletion of *jmjd-5*, placed on the rectangle showing the background into which the deletion was introduced. The box plots for the parental strains (N2 and CB4856, on the left) are colored according to genotype

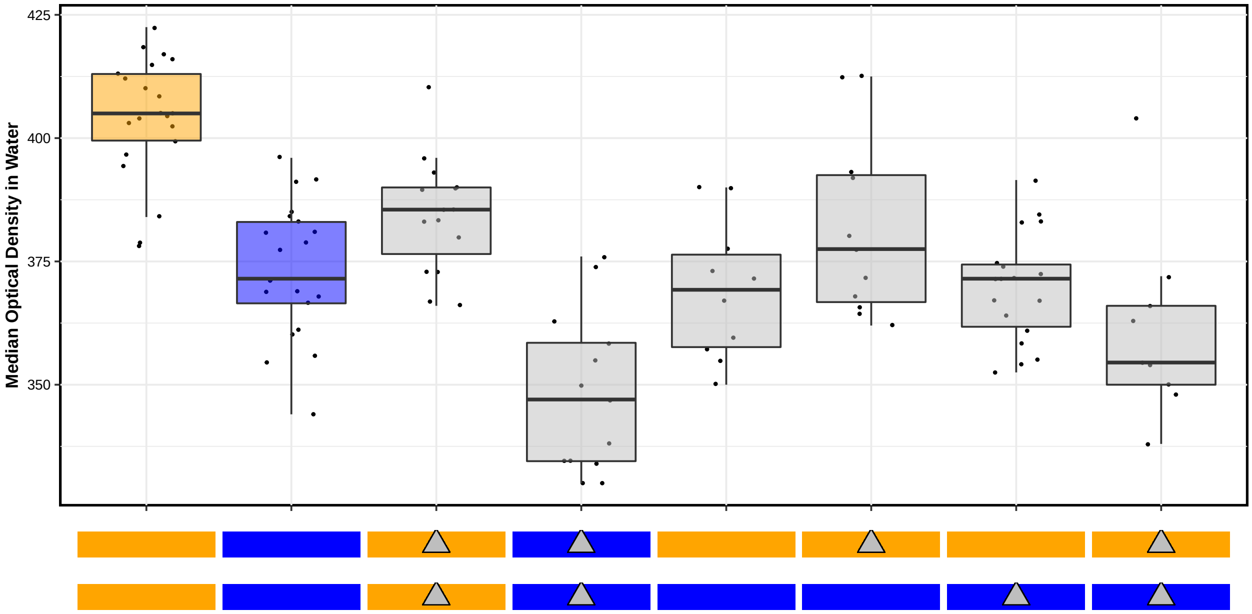


**Figure S12**

Control-condition phenotypes of the strains tested in the reciprocal hemizygosity experiment of *jmjd-5* are shown. The y-axis shows the pruned median optical density upon exposure to the control condition (lysate in K medium plus 1% distilled water) for each strain reported along the x-axis. Control-condition responses are reported as Tukey box plots where each point is a biological replicate. The genotypes of each strain are shown as colored rectangles beneath each box plot, where each rectangle represents a homolog (orange rectangles are an N2 genotype, and blue rectangles are a CB4856 genotype). The maternal homolog is shown on top and the paternal homolog is shown on bottom. Grey triangles indicate a deletion of *jmjd-5*, placed on the rectangle showing the background into which the deletion was introduced. The box plots for the parental strains (N2 and CB4856, on the left) are colored according to genotype

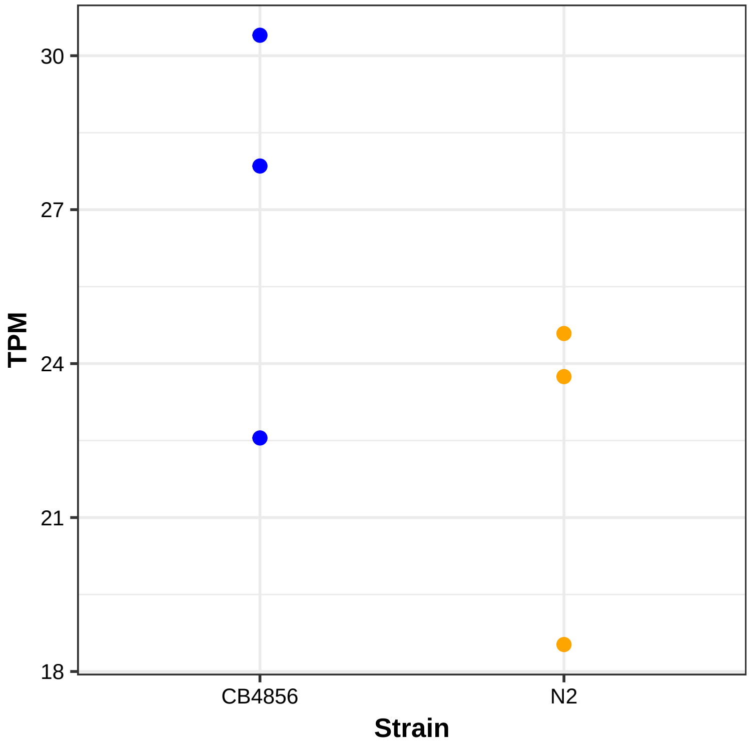


**Figure S13**

RNA-seq measurements of *scb-1* expression for young adult populations of N2 and CB4856 are reported. The x-axis indicates the sample (N2 or CB4856), and the y-axis shows the transcripts per million (TPM) estimate for each replicate. Replicates are plotted as dots, colored by strain (orange for N2 and blue for CB4856).

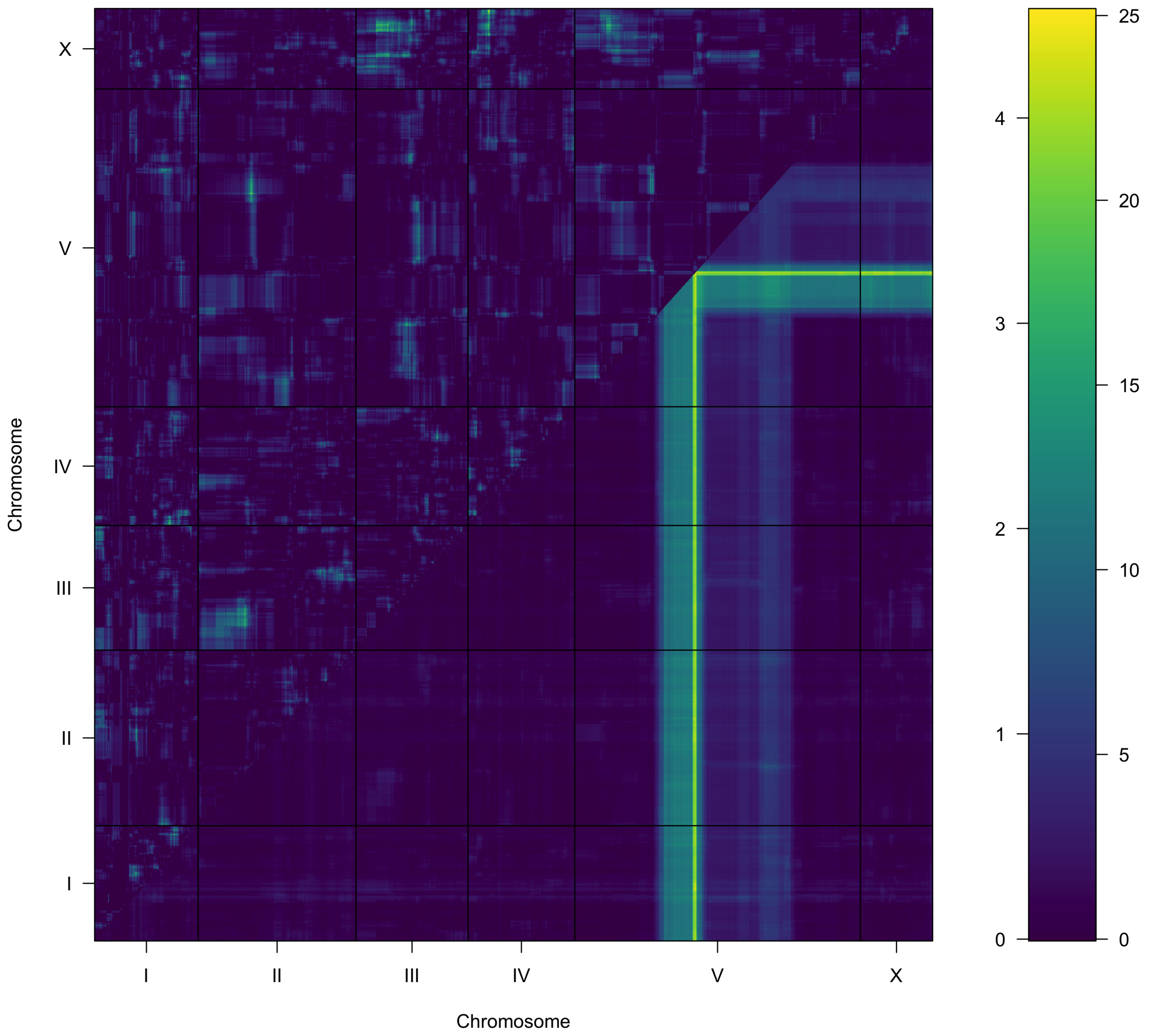


**Figure S14**

A two-factor genome scan for *scb-1* expression variation is shown. Log of the odds (LOD) scores are shown for each pairwise combination of loci, split by chromosome. The upper-left triangle contains the epistasis LOD scores, and the lower-right triangle contains LOD scores for the full model. LOD scores are shown as colors, with lower scores in blue and higher scores in yellow, as shown in the color scale. The epistatic LOD score axis is on the left of the color scale and the full LOD score axis is on the right of the color scale.

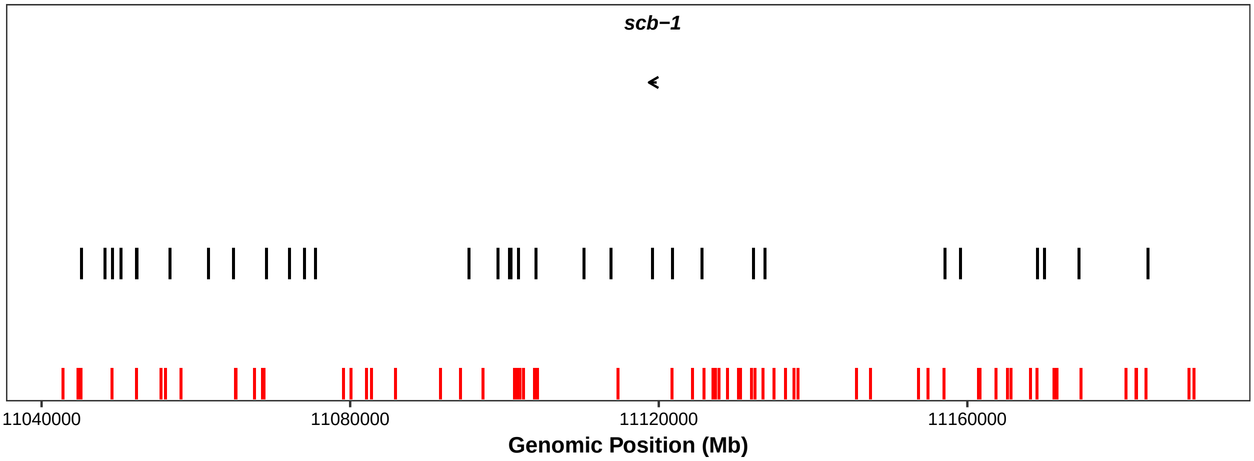


**Figure S15**

All variants in the QTL confidence interval for which CB4856 contains the alternate allele are plotted as vertical lines. On the x-axis, genomic position of the variant is shown. Variants are colored by minor allele frequency (less than 0.05 = red, greater than 0.05 = black). The location of *scb-1* is shown as an arrow, labeled with the gene name.

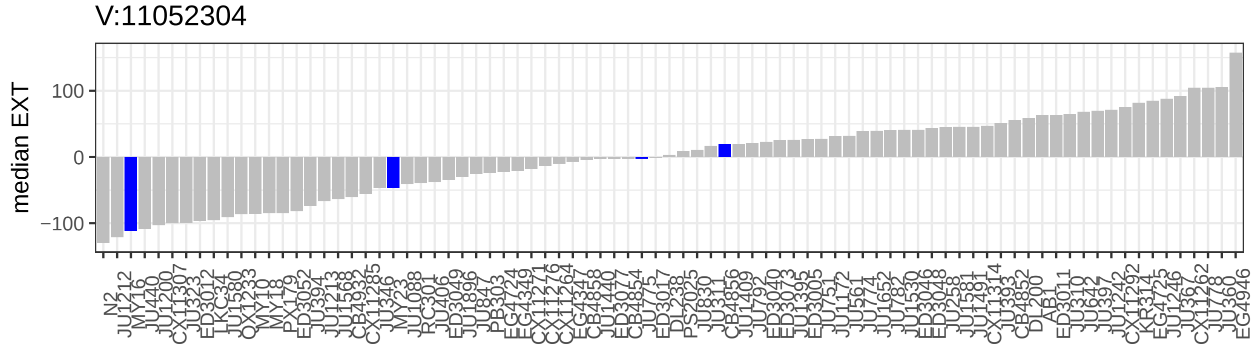
V:11052304
median EXT
100
0
−100
N2 JU1212 MY16 JU440 JU1200 CX11307 JU323 ED3012 LKC34 JU1580 QX1233 MY10 MY18 PX179 ED3052 JU394 JU1213 JU1568 CB4932 CX11285 JU346 MY23 JU1088 RC301 JU406 ED3049 JU1896 JU847 PB303 EG4724 EG4349 CX11271 CX11276 CX11264 EG4347 CB4858 JU1440 ED3077 CB4854 JU775 ED3017 DL238 PS2025 JU830 JU311 CB4856 JU1409 JU792 ED3040 ED3073 JU1395 ED3005 JU751 JU1172 JU561 JU774 JU1652 JU782 JU1530 ED3046 ED3048 JU258 JU1581 JU1491 CX11314 JU393 CB4852 DL200 AB1 ED3011 JU310 JU642 JU397 JU1242 CX11292 KR314 EG4725 JU1246 JU367 CX11262 JU778 JU360 EG4946

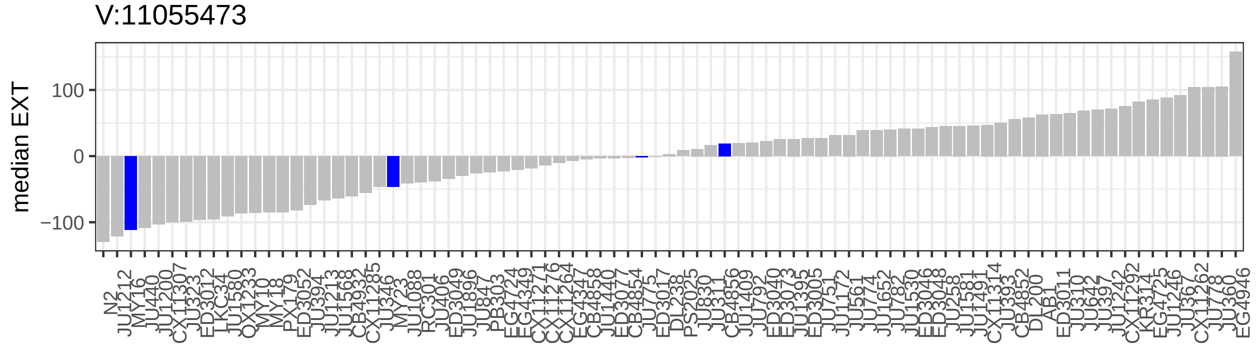
V:11055473
median EXT
100
0
−100
N2 JU1212 MY16 JU440 JU1200 CX11307 JU323 ED3012 LKC34 JU1580 QX1233 MY10 MY18 PX179 ED3052 JU394 JU1213 JU1568 CB4932 CX11285 JU346 MY23 JU1088 RC301 JU406 ED3049 JU1896 JU847 PB303 EG4724 EG4349 CX11271 CX11276 CX11264 EG4347 CB4858 JU1440 ED3077 CB4854 JU775 ED3017 DL238 PS2025 JU830 JU311 CB4856 JU1409 JU792 ED3040 ED3073 JU1395 ED3005 JU751 JU1172 JU561 JU774 JU1652 JU782 JU1530 ED3046 ED3048 JU258 JU1581 JU1491 CX11314 JU393 CB4852 DL200 AB1 ED3011 JU310 JU642 JU397 JU1242 CX11292 KR314 EG4725 JU1246 JU367 CX11262 JU778 JU360 EG4946

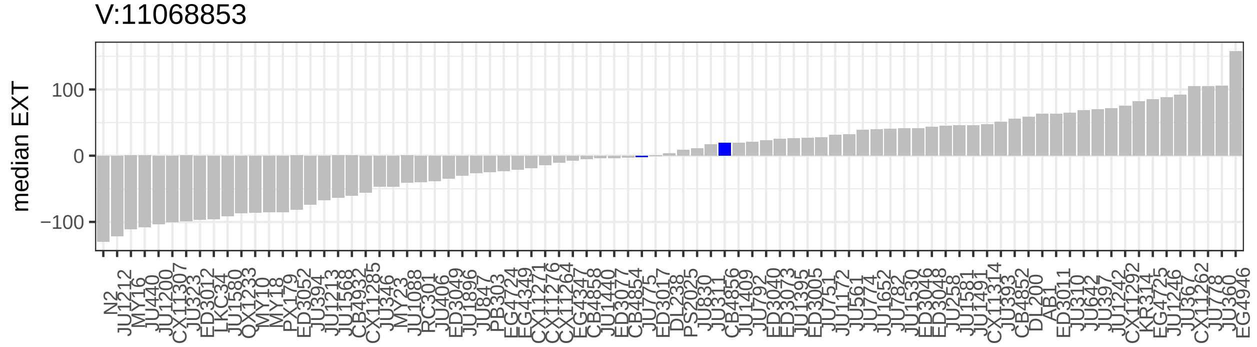
V:11068853
median EXT
100
0
−100
N2 JU1212 MY16 JU440 JU1200 CX11307 JU323 ED3012 LKC34 JU1580 QX1233 MY10 MY18 PX179 ED3052 JU394 JU1213 JU1568 CB4932 CX11285 JU346 MY23 JU1088 RC301 JU406 ED3049 JU1896 JU847 PB303 EG4724 EG4349 CX11271 CX11276 CX11264 EG4347 CB4858 JU1440 ED3077 CB4854 JU775 ED3017 DL238 PS2025 JU830 JU311 CB4856 JU1409 JU792 ED3040 ED3073 JU1395 ED3005 JU751 JU1172 JU561 JU774 JU1652 JU782 JU1530 ED3046 ED3048 JU258 JU1581 JU1491 CX11314 JU393 CB4852 DL200 AB1 ED3011 JU310 JU642 JU397 JU1242 CX11292 KR314 EG4725 JU1246 JU367 CX11262 JU778 JU360 EG4946

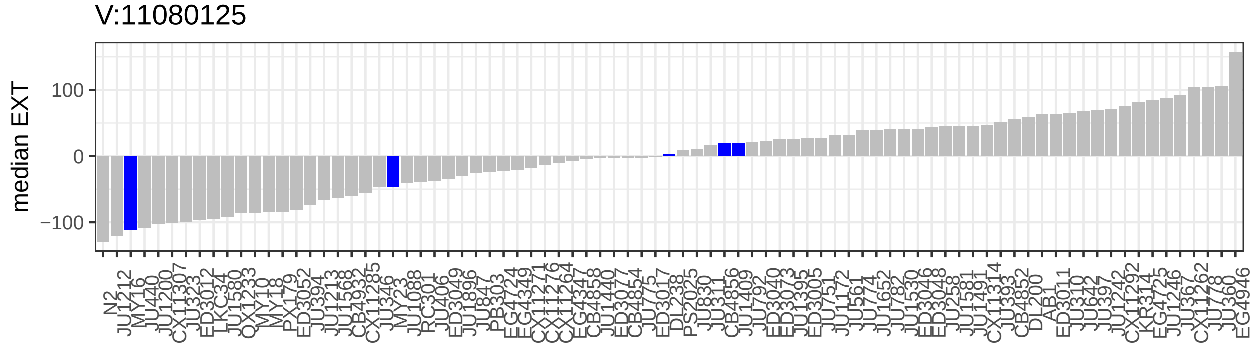
V:11080125
median EXT
100
0
−100
N2 JU1212 MY16 JU440 JU1200 CX11307 JU323 ED3012 LKC34 JU1580 QX1233 MY10 MY18 PX179 ED3052 JU394 JU1213 JU1568 CB4932 CX11285 JU346 MY23 JU1088 RC301 JU406 ED3049 JU1896 JU847 PB303 EG4724 EG4349 CX11271 CX11276 CX11264 EG4347 CB4858 JU1440 ED3077 CB4854 JU775 ED3017 DL238 PS2025 JU830 JU311 CB4856 JU1409 JU792 ED3040 ED3073 JU1395 ED3005 JU751 JU1172 JU561 JU774 JU1652 JU782 JU1530 ED3046 ED3048 JU258 JU1581 JU1491 CX11314 JU393 CB4852 DL200 AB1 ED3011 JU310 JU642 JU397 JU1242 CX11292 KR314 EG4725 JU1246 JU367 CX11262 JU778 JU360 EG4946

V:11085892

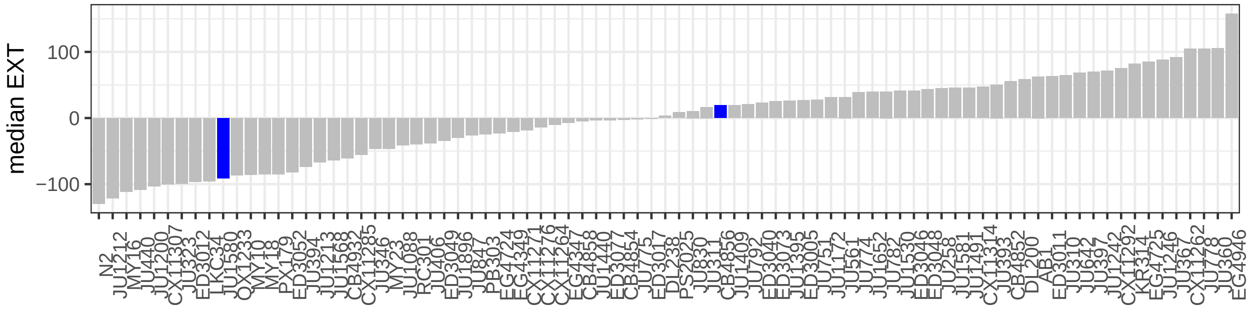


V:11097232

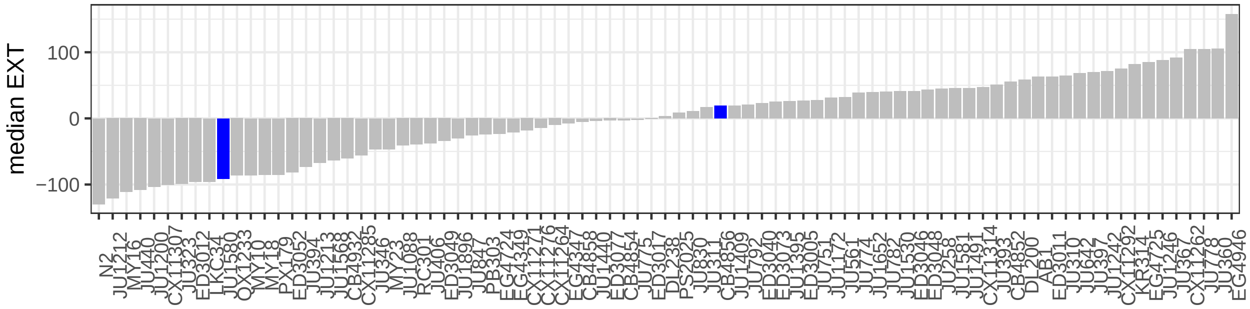


V:11101282

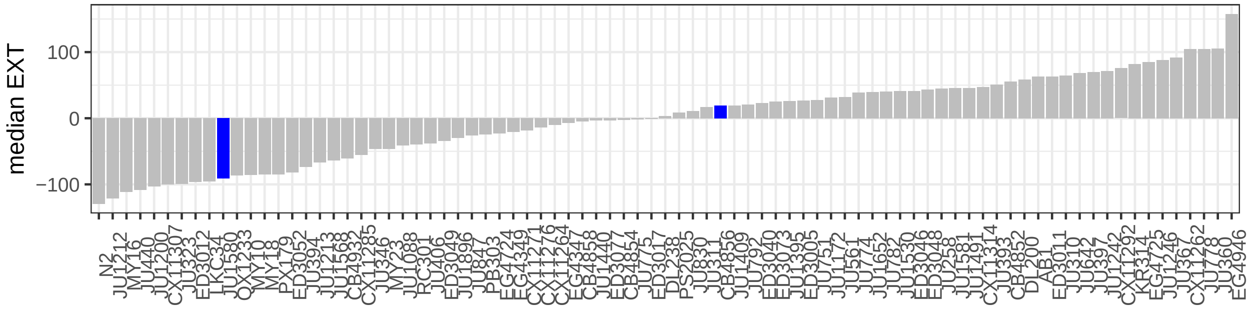


V:11102451

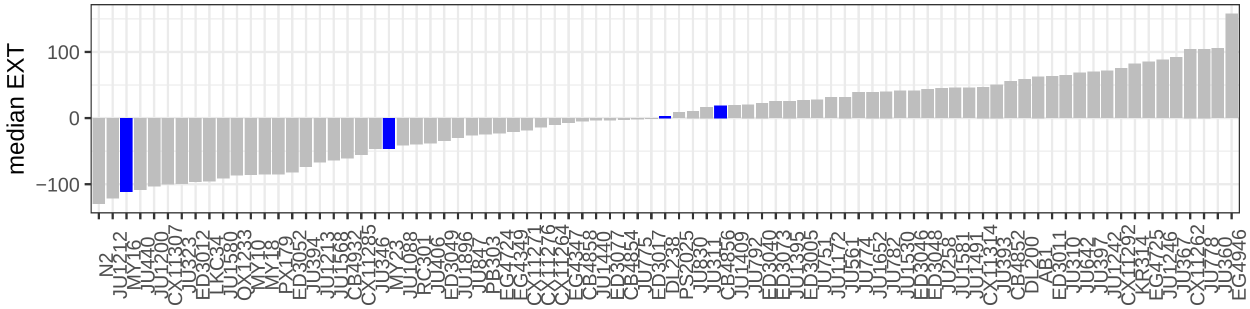

V:11104309

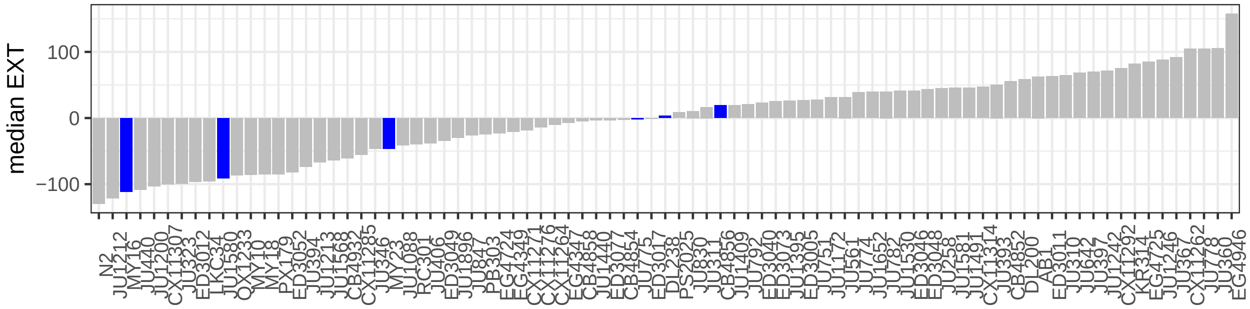


V:11125840

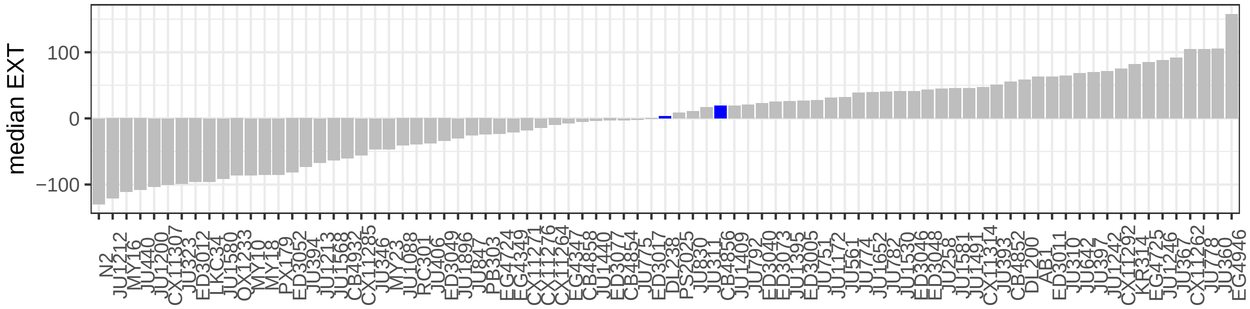


V:11127057

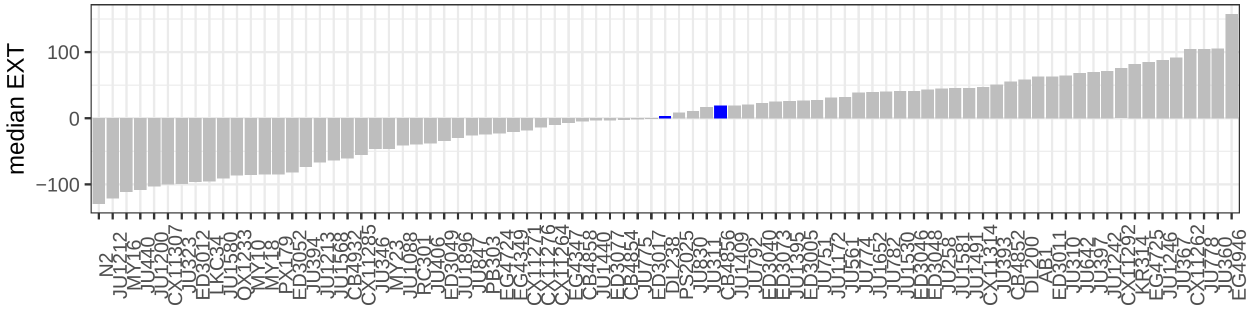


V:11127063

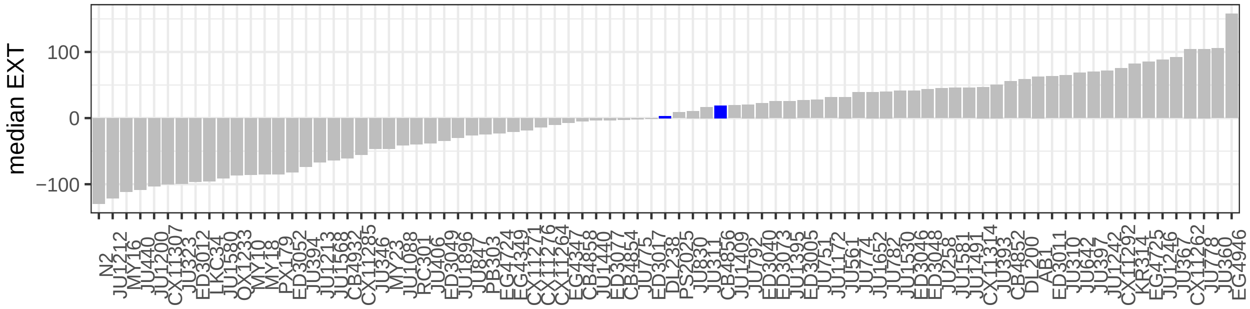

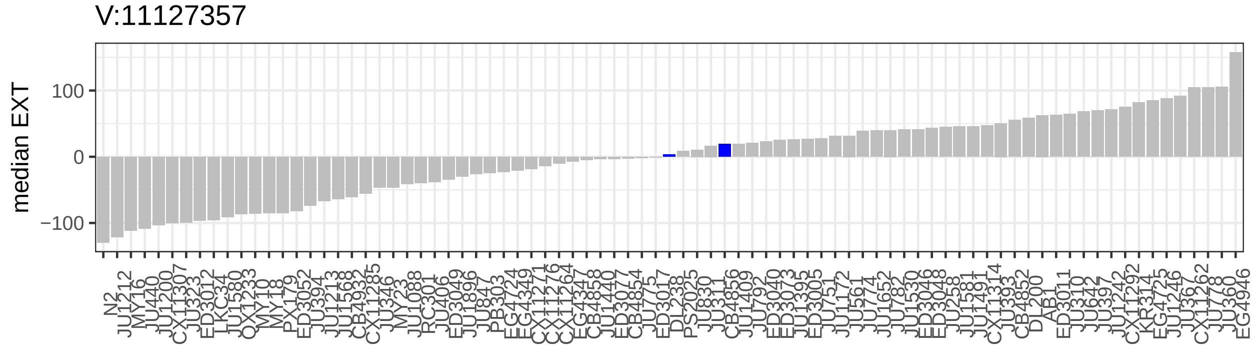
V:11127357
median EXT
100
0
−100

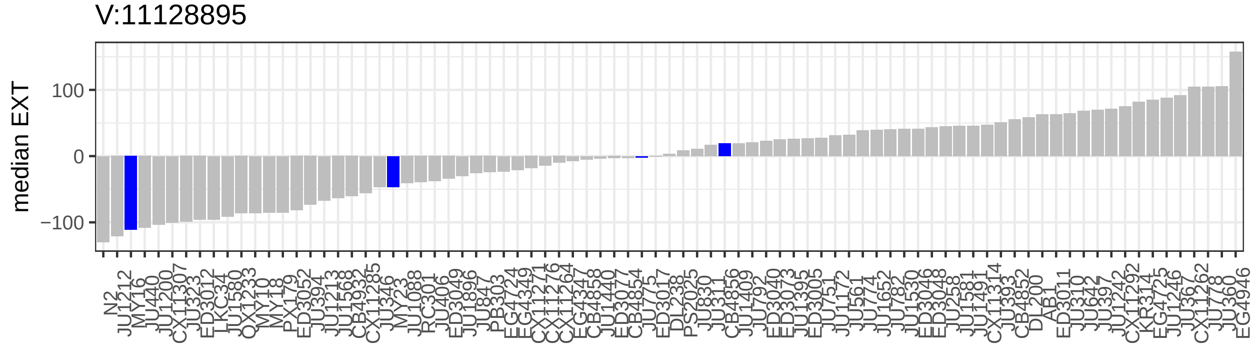
V:11128895
median EXT
100
0
−100

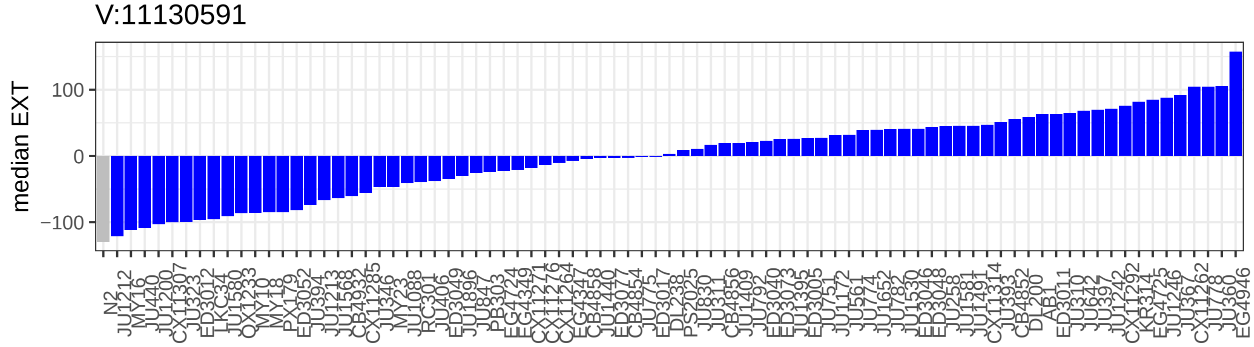
V:11130591
median EXT
100
0
−100

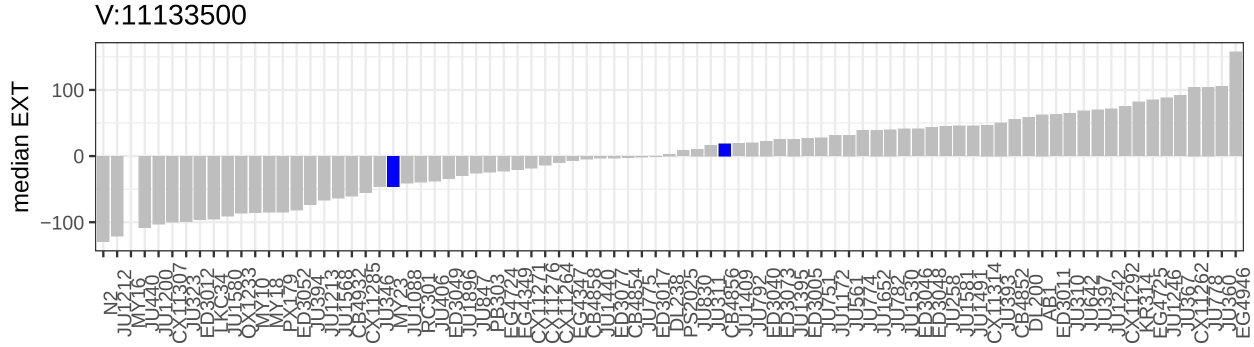
V:11133500
median EXT
100
0
−100

V:11133501

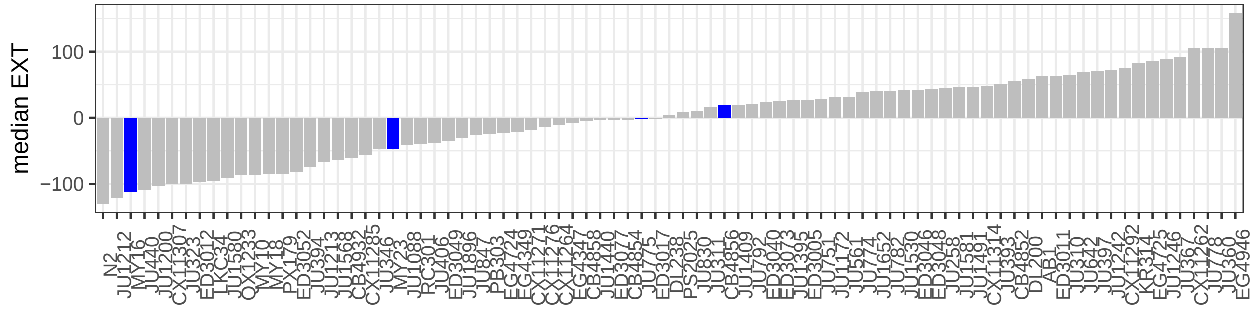


V:11154887

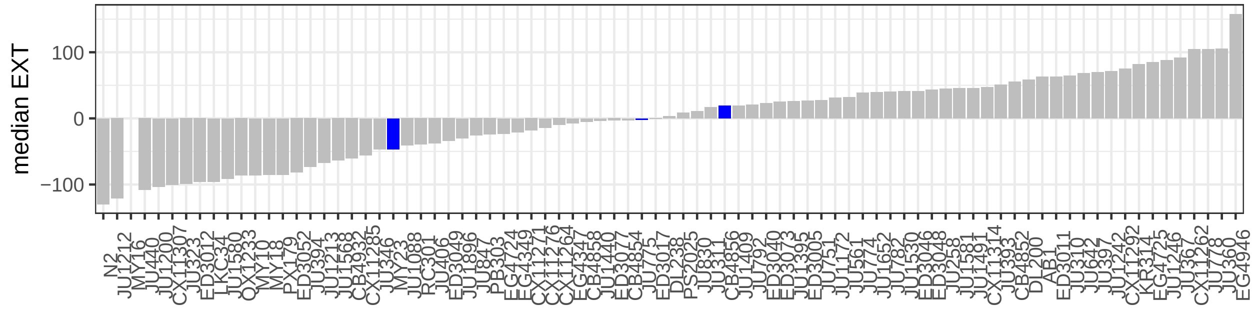


V:11163719

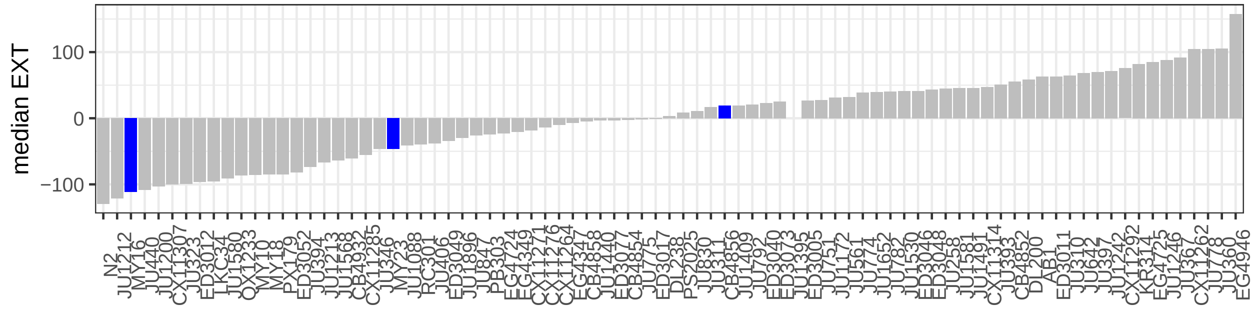


V:11168198

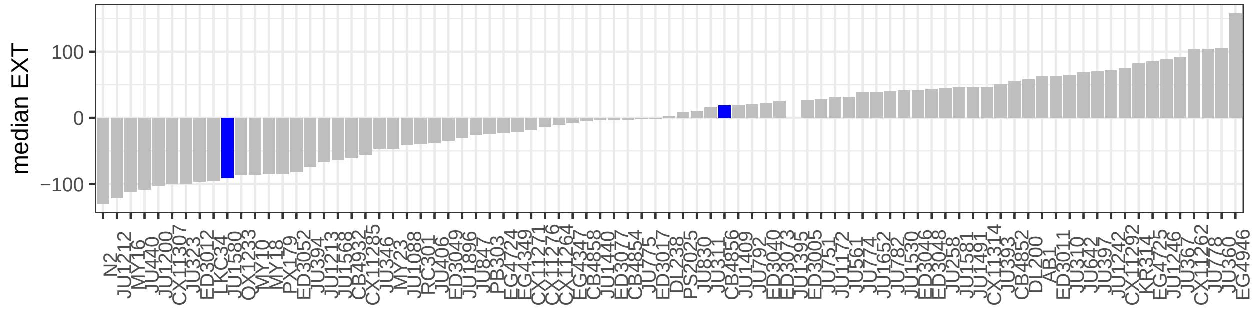

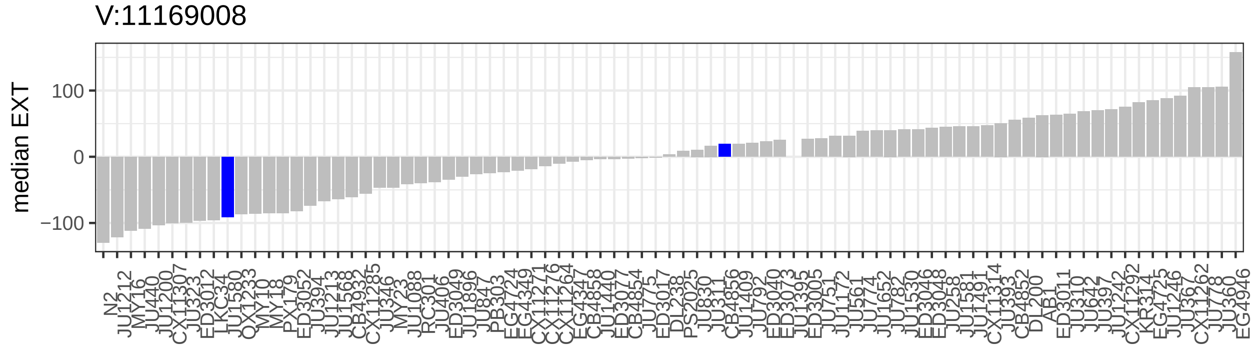
V:11169008
median EXT
100
0
−100

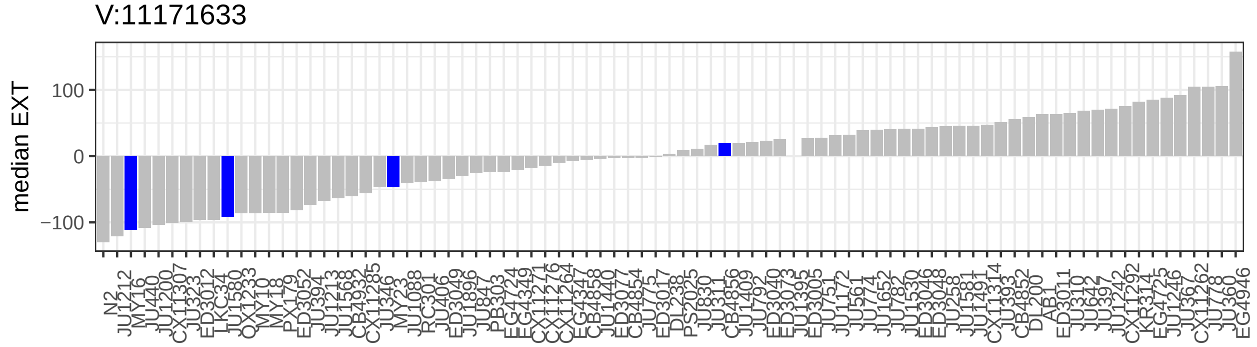
V:11171633
median EXT
100
0
−100

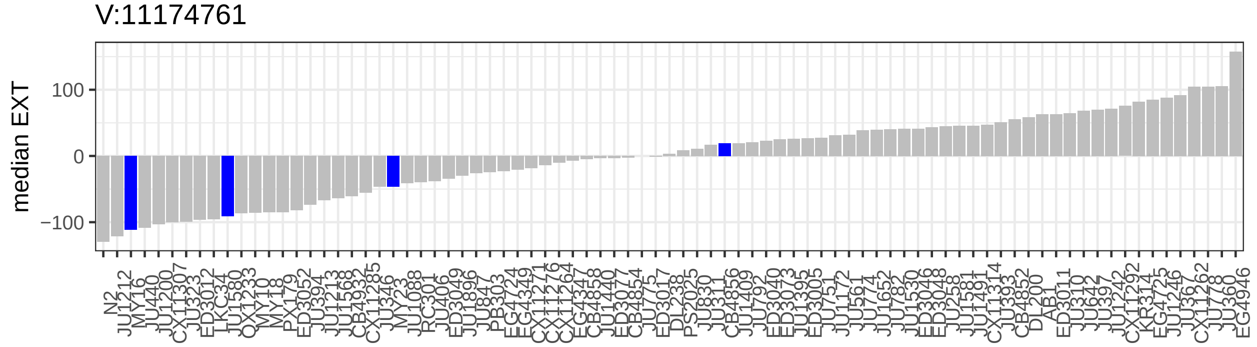
V:11174761
median EXT
100
0
−100

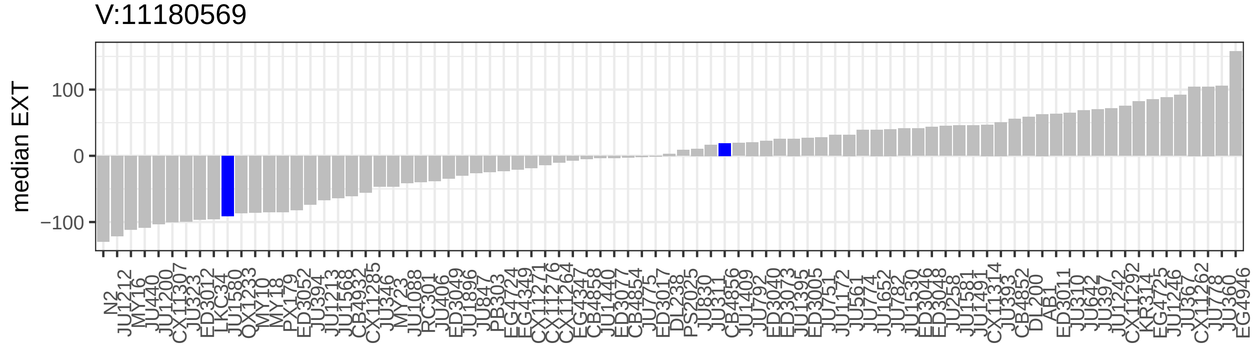
V:11180569
median EXT
100
0
−100

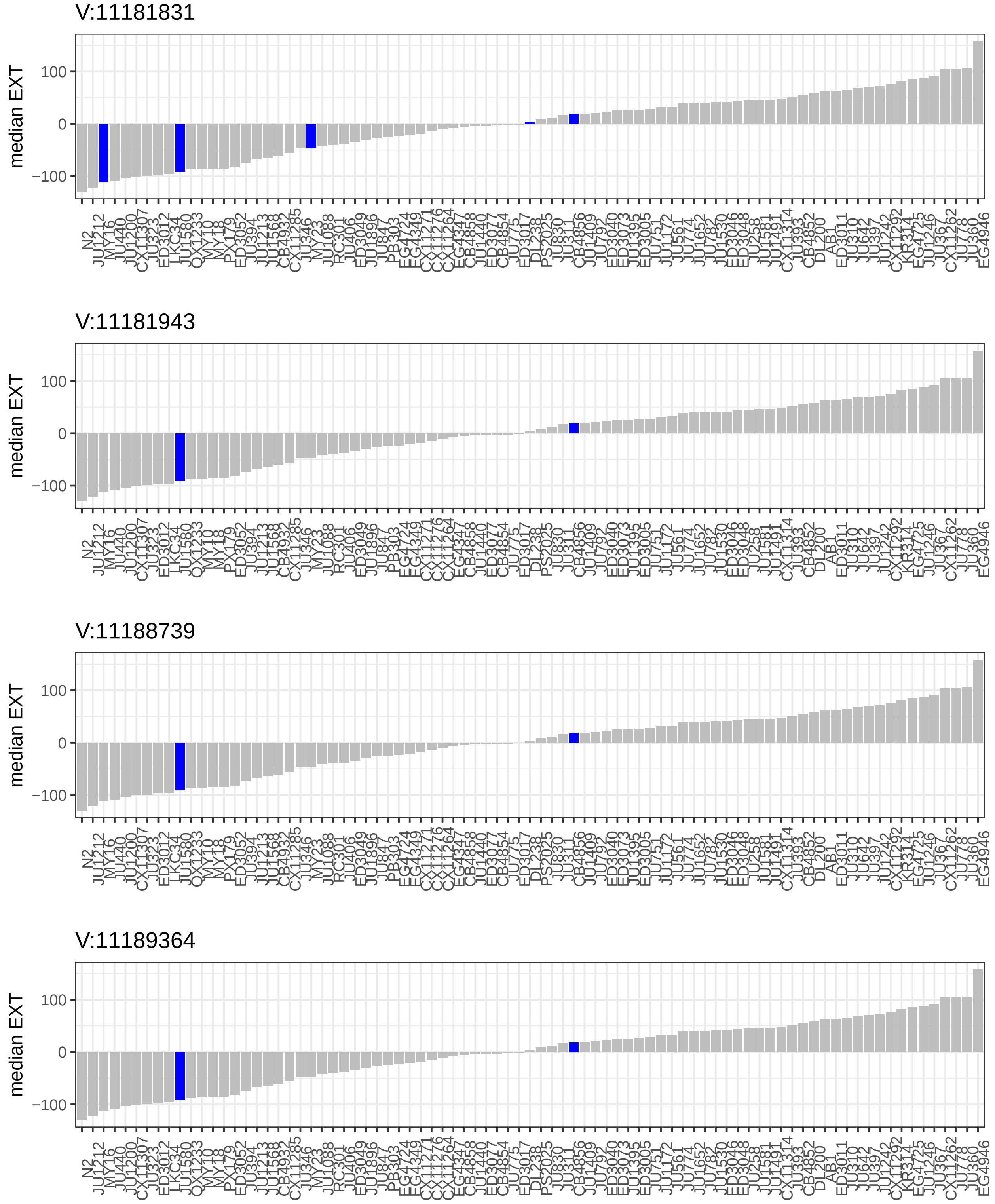


**Figure S16**

Phenotypes of all wild isolates assayed in bleomycin are shown for each of the 28 variants in the QTL region that are rare (MAF < 0.05) but not unique to CB4856. Each plot shows a bar plot indicating the distribution of residual bleomycin median optical density phenotypes across all 83 wild isolates assayed, arranged by phenotype. The x-axis indicates the strain name, and the y-axis indicates the phenotypic value. For each of the 28 variants, the position is written above the phenotypic distribution. Strains with the alternate allele at that site are colored blue, and strains with the reference allele are colored in grey.

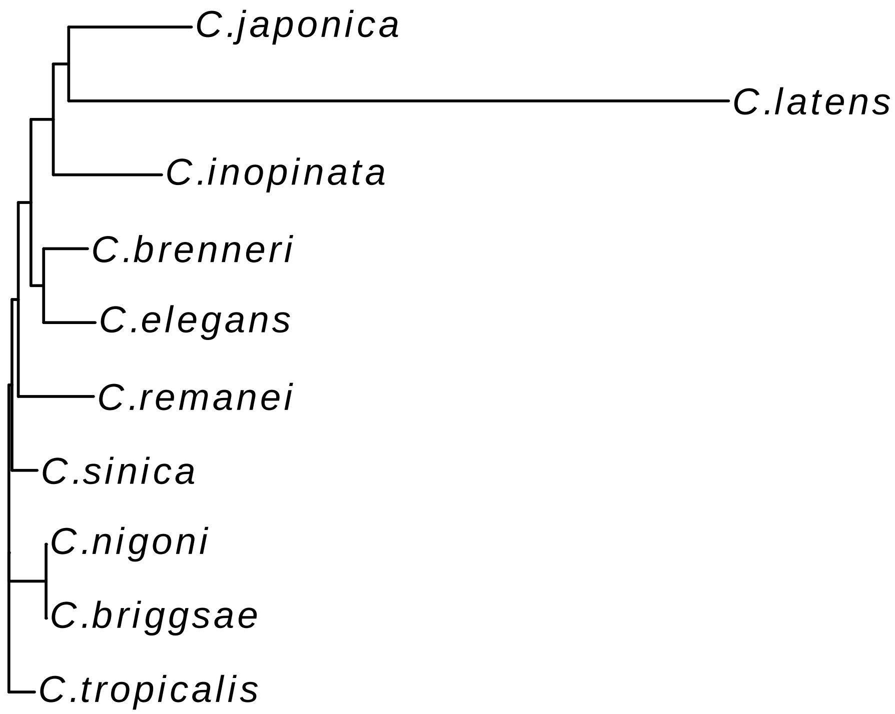


**Figure S17**

A neighbor-joining tree from from a multiple-sequence alignment of SCB-1 homologs is shown.
