## Supplemental Information for "A nematode-specific gene underlies bleomycin-response variation in *Caenorhabditis elegans*"

### Reagents used to generate NILs

| | | |
|---|---|---|
| **ECA230** ***eanIR150(V N2>CB4856)*** | Constructed from: QX131 x CB4856 | N2 into CB4856 |
| **ECA232** ***eanIR152(V CB4856>N2)*** | Constructed from: QX450 x N2 | CB4856 into N2 |
| **ECA411** ***eanIR185(V N2>CB4856)*** | Constructed from: ECA230 x CB4856 | N2 into CB4856 |
| **ECA528** ***eanIR302(V N2>CB4856)*** | Constructed from: ECA230 x CB4856 | N2 into CB4856 |
| Left indel - V: 7,862,556 | | |
| oECA799 | TTCTCGCTACTGGAACACGC | |
| oECA800 | TCAAGAAGCGTTGGGAAGTCT | |
| Right indel - V: 13,110,045 | | |
| oECA745 | TGCAGAGGTGGAGTAACCCT | |
| oECA746 | CTCGGTCTCTCCCCCACTAA | |

### Reagents used to generate genome-edited alleles

crECA36 *dpy-10* guide RNA: GCUACCAUAGGCACCACGAG
crECA37 *dpy-10* repair construct:
CACTTGAACTTCAATACGGCAAGATGAGAATGACTGGAAACCGTACCGCATGCGGTGCCTA GGTAGCGGAGCTTCACAT GGCTTCAGACCAACAGCCTAT

| ***jmjd-5(deletion)*** | |
|---|---|
| ECA1047 *jmjd-5(ean136)* | CB4856 background |
| ECA1048 *jmjd-5(ean137)* | CB4856 background |
| ECA1051 *jmjd-5(ean141)* | N2 background |
| ECA1052 *jmjd-5(ean142)* | N2 background |

| guide RNAs | |
|---|---|
| crECA58 | TGGTACAAAACTATTTCGGA |
| crECA59 | AAAATTGACGAGTGTCGCGA |
| confirmation primers - external | |
| oECA1153 | TCCTCGTATTACAATCCGTTGTCCA |
| oECA1193 | TGTCGTCTGGAAACATATGGCT |
| confirmation primers - internal | |
| oECA1194 | CCGATAAAGGGCTGTGTATGGG |
| oECA1195 | TCGAAAAGGCGATGTTGTGCAA |

| ***C45B11.8(deletion)*** | |
|---|---|
| ECA996 *C45B11.8(ean103)* | CB4856 background |
| ECA998 *C45B11.8(ean105)* | CB4856 background |
| ECA699 *C45B11.8(ean60)* | N2 background |
| ECA700 *C45B11.8(ean61)* | N2 background |
| guide RNAs | |
| crECA22 | GCTGCAGTAGAGGTGACATT |
| crECA23 | GAAGAAGTGAAAGAAGTGGG |
| confirmation primers - external | |
| oECA1269 | TCTCCGTGACTCAAATTTCGACA |
| oECA1270 | AGATGAAGATCACACTCTTGCGA |
| confirmation primers - internal | |
| oECA1267 | TCGCCTAATGTCACCTCTACTGC |
| oECA1268 | TCCTCCCACTTCTTTCACTTCT |

| ***C45B11.6(deletion)*** | |
|---|---|
| ECA1053 *C45B11.6(ean142)* | CB4856 background |
| ECA622 *C45B11.6(ean46)* | CB4856 background |
| ECA623 *C45B11.6(ean47)* | N2 background |
| ECA624 *C45B11.6(ean48)* | N2 background |
| guide RNAs | |
| crECA9 | TCATCAGGATCAATTTCAAG |
| crECA10 | AGAATATCTGAATTGCCGAA |
| confirmation primers - external | |
| oECA1226 | TCCTGGTTTTTCTTTTCAGTGGTTGT |
| oECA1227 | TGTCTTCGGCAATTTTGTGCCC |
| confirmation primers - internal | |
| oECA1224 | GCTGGATTGCATTTGTCAAACCC |
| oECA1225 | AGTTAAGAAAAGCAGCACCTGGA |

| ***srg-42(deletion)*** | |
|---|---|
| ECA1012 *srg-42(ean119)* | CB4856 background |
| ECA1013 *srg-42(ean120)* | CB4856 background |
| ECA697 *srg-42(ean58)* | N2 background |
| ECA698 *srg-42(ean59)* | N2 background |
| guide RNAs | |
| crECA19 | TCAATTACAAACTAGCGATT |
| crECA20 | AGATGGTAAACCATAAATAG |

| confirmation primers - external | |
|---|---|
| oECA1262 | TCACGCGTCACAATTATTGCTGA |
| oECA1263 | AGCCATTGTTCAATTTCCCAGGT |
| confirmation primers - internal | |
| oECA1260 | AGGGACAGTTATGATCACCAGT |
| oECA1261 | GCCTGGCCCTTTTCAGAGACAA |

| ***cnc-10(deletion)*** | |
|---|---|
| ECA687 *cnc-10(ean53)* | CB4856 background |
| ECA696 *cnc-10(ean57)* | CB4856 background |
| ECA692 *cnc-10(ean55)* | N2 background |
| ECA693 *cnc-10(ean56)* | N2 background |
| guide RNAs | |
| oECA1186* | ACAACGTCTGCTCAATTTTA |
| crECA21* | ACGTCTGCTCAATTTTATGG |
| oECA1187 | GCACTAATGGGAGCTGCAAT |
| confirmation primers | |
| oECA1170** | GTCCTTACTGAGGCGTGTCCAT |
| oECA1266** | TCCAGGATCTACGCAAAAATGAACT |
| oECA1171 | CAGGTTCAAATCCTGCGGACAG |

*oECA1186 and oECA1187 were used to generate the CB4856 deletions. crECA21 and oECA1187 were used to generate the N2 deletions.
**oECA1170 and oECA1171 were used to confirm CB4856 deletions. oECA1266 and oECA1171 were used to confirm N2 deletions.

| ***H19N07.3(deletion)*** |
|---|

| | |
|---|---|
| ECA1133 *H19N07.3(ean179)* | CB4856 background |
| ECA1134 *H19N07.3(ean180)* | CB4856 background |
| ECA1131 *H19N07.3(ean177)* | N2 background |
| ECA1132 *H19N07.3(ean178)* | N2 background |
| guide RNAs | |
| crECA84 | GCGAGCACAACTTCAAGAAA |
| crECA85 | CGTATGGCTGCCAAGGCCAG |
| confirmation primers | |
| oECA1173 | TCTTGCAGACACATGGGTCC |
| oECA1174 | ATCGGTGGGCACAATGTGAT |

| ***jmjd-5(CB4856 to N2)*** | |
|---|---|
| ECA578 *jmjd-5(ean12[S338P])* | CB4856 background |
| ECA579 *jmjd-5(ean13[S338P])* | CB4856 background |
| guide RNA | |
| oECA1196 | GGAATTTGAAAGTGGAATTA |
| repair template | |
| oECA1199 | ACTAGCATGGTTAATTCATGAAAATTTACCTGGTGTGTCATCTGATGATTGGATTCATTCGAGTTTTCAGTTCAATACAACTAATACGTATCCTGCGTTAATTCCACTTTCAAATTCCAAATCTATCGATGAATGTGATGAAGATGA |
| confirmation primers - to check for *Bsa*AI site introduction | |
| oECA1194 | CCGATAAAGGGCTGTGTATGGG |
| oECA1195 | TCGAAAAGGCGATGTTGTGCAA |

| ***jmjd-5(N2 to CB4856)*** | |
|---|---|
| ECA576 *jmjd-5(ean10[P338S])* | N2 background |
| ECA577 *jmjd-5(ean11[P338S])* | N2 background |
| guide RNA | |
| oECA1196 | GGAATTTGAAAGTGGAATTA |
| repair template | |
| oECA1198 | ACTAGCATGGTTAATTCATGAAAATTTACC TGGTGTGTCATCTGATGATTGGATTCATT CGAGTTTTCAGTTCAATACAACTAATACGT ATTCTGCGTTAATTCCACTTTCAAATTCCA AATCTATCGATGAATGTGATGAAGATGA |
| confirmation primers - to check for *BsaA*I site introduction | |
| oECA1194 | CCGATAAAGGGCTGTGTATGGG |
| oECA1195 | TCGAAAAGGCGATGTTGTGCAA |

**Data availability:** All data are available on figshare. **File S1** contains all pruned data from the high-throughput bleomycin assays. **File S2** contains the broad-sense heritability estimates calculated for each drug concentration for all 26 HTA traits for the HTA dose response as well as for the 24 HTA traits in the modified HTA dose response. **File S3** contains all control-regressed data for the 26 HTA traits for all assays. **File S4** contains the annotated linkage mapping data for the 26 control-regressed HTA traits. **File S5** is a VCF that reports the genotype from whole-genome sequence for all NILs in the manuscript. **File S6** is a simplified version of **File S5** that contains information on recombination locations for all NILs and can be used for more user-friendly visualization of NIL genotypes. **File S7** contains all statistical information for HTA phenotypic differences reported in the manuscript. **File S8** is a summary of the scantwo analysis for bleomycin responses in the RIAILs and reports the maximum interaction LOD score for each chromosome pair. **File S9** contains information on all genes in the QTL confidence interval plus 20 kb on either side. **File S10** contains locations of the exons, introns, and transcription start and stop sites for all genes in the region. **File S11** reports predicted non-synonymous variants between the N2 and CB4856 strains in the region. **File S12** is derived from the Rockman *et al.* 2010 RIAIL microarray expression data, and reports the expression measurements for each of the 13,107 microarray probes across 209 RIAILs. **File S13** contains all significant QTL identified by linkage mapping of **File S12** data. **File S14** contains the annotated linkage mapping of the *H19N07.3* expression data. **File S15** reports the *H19N07.3* expression and residual median optical density for strains of the

RIAIL panel that were assayed for both of those traits. **File S16** contains *H19N07.3* RNA-seq expression data for populations of young adults of N2 and CB4856. **File S17** is a summary of the scantwo analysis for *H19N07.3* expression in the RIAILs and reports the maximum interaction LOD score for each chromosome pair. **File S18** contains control-regressed phenotypic data for all wild isolates assayed in response to bleomycin. **File S19** contains genome-wide association mapping for the phenotypes in **File S18**. **File S20** contains genotype information for each strain measured in **File S18** across all variants within the linkage mapping confidence interval around the QTL for which CB4856 contains the alternate allele. **File S21** is a FASTA file containing the protein sequences for all *H19N07.3* homologs. **File S22** is a neighbor-joining tree derived from a multiple sequence alignment of all sequences from **File S21** in Newick tree format.

**File S1 -- allpruned.csv**

| **Column** | **Description** |
|---|---|
| date | Date on which the assay was scored, in YYYYMMDD format |
| experiment | The experiment run on that date - either HTAdose (dose response with four wild isolates), RIAILs (for linkage mapping), largeNIL (ECA230/ECA232 comparison), smallNIL (ECA411/ECA528 comparison), deletions (CRISPR/Cas9 deletions of *C45B11.8, C45B11.6, jmjd-5, srg-42,* and *cnc-10* in both parental backgrounds), jmjdswap (reciprocal allele replacement of *jmjd-5*), hemi_dose (dose response for hemizygosity assay), hemizygosity_jmjd (hemizygosity of *jmjd-5* deletion), H19 (*H19N07.3* deletions), or hemizygosity_H19 (hemizygosity of *H19N07.3* deletion). |
| round | In the case of RIAIL experiments, numerical value used to aggregate plates that tested the same condition across multiple days |
| assay | In the case of RIAIL assays, letter indicating the experimental block |
| plate | The numerical value of a 96-well plate |
| condition | The condition present in a given well. “Bleomycin” indicates a 50 µM concentration in the case of the standard HTA or 12.5 µM in the case of the hemizygosity assay. “Water” or “1percwater” indicates the control condition. “Bleomycin” followed by a number indicates the concentration of bleomycin in µM for dose responses. |
| control | If applicable, the control condition for a given well (to be used in control regression). |
| strain | Strain name in a given well |
| row | Letter indicating the row of a 96-well plate |
| col | Numerical value indicating the column of a 96-well plate |

| | |
|---|---|
| trait | Population parameter measured |
| phenotype | Numerical value indicating the trait measurement for a given well |

**File S2 -- dose_H2.csv**

| Column | Description |
|---|---|
| condition | The condition to which animals were exposed, either “water” or “bleomycin” followed by the concentration in µM |
| trait | The population parameter measured in the dose response for which heritability was calculated |
| $H^2$ | The broad-sense heritability estimate, calculated using *lmer::lme4(phenotype ~ 1 + 1\|strain)* with dose-response data |
| experiment | Either HTA_dose (the original dose response) or hemi_dose (the dose response using the alternate version of the HTA) that indicates from which asay the data are derived |

**File S3 -- allregressed.csv**

| Column | Description |
|---|---|
| date | Date on which the assay was scored, in YYYYMMDD format |
| experiment | The experiment run on that date - RIAILs (for linkage mapping), largeNIL (ECA230/ECA232 comparison), smallNIL (ECA411/ECA528 comparison), deletions (CRISPR/Cas9 deletions of *C45B11.8, C45B11.6, jmjd-5, srg-42,* and *cnc-10* in both parental backgrounds), jmjdswap (reciprocal allele replacement of *jmjd-5*), hemizygosity_jmjd (hemizygosity of *jmjd-5* deletion), H19 (*H19N07.3* deletions), or hemizygosity_H19 (hemizygosity of *H19N07.3* deletion). |
| round | In the case of RIAIL experiments, numerical value used to aggregate plates that tested the same condition across multiple days |
| assay | In the case of RIAIL assays, letter indicating the experimental block |
| condition | The condition present in a given well. “Bleomycin” indicates a 50 µM concentration in the case of the standard HTA or 12.5 µM in the case of the hemizygosity assay. “Water” or “1percwater” indicates the control condition. “Bleomycin” followed by a number indicates the concentration of bleomycin in µM for dose responses. |

| | |
|---|---|
| control | If applicable, the control condition for a given well (to be used in control regression). |
| plate | The numerical value of a 96-well plate |
| row | Letter indicating the row of a 96-well plate |
| col | Numerical value indicating the column of a 96-well plate |
| strain | Strain name in a given well |
| trait | Population parameter measured |
| phenotype | Numerical value indicating the trait measurement for a given well |

**File S4 -- annotatedLODs.csv**

| **Column** | **Description** |
|---|---|
| marker | Genetic marker at which the correlation between genotype and phenotype was tested |
| chr | Chromosome on which the marker can be found |
| pos | Position, in bp, at which the genetic marker can be found |
| trait | HTA trait for which RIAIL phenotypes were measured |
| lod | Log of odds ratio for correlation between genotype at the genetic marker and phenotype of RIAILs for the trait of interest |
| threshold | Genome-wide error rate threshold for a particular iteration of the mapping, above which a LOD is considered significant |
| iteration | Numerical value indicating the mapping-process iteration |
| var_exp | If applicable (in the case of a significant QTL), amount of phenotypic variation across RIAILs that can be explained by genetic variation at the QTL. |
| eff_size | If applicable (in the case of a significant QTL), effect size of the QTL, calculated as the slope of a linear model with the formula (phenotype ~ genotype). A positive value indicates that RIAILs with the CB4856 allele have more positive phenotypes than those with the N2 allele, and *vice versa*. |
| ci_l_marker | The genetic marker indicating the left boundary of a 95% confidence interval around the peak marker |
| ci_l_pos | The position, in bp, of the left boundary of a 95% confidence interval around the peak marker |

| | |
|---|---|
| ci_r_marker | The genetic marker indicating the right boundary of a 95% confidence interval around the peak marker |
| ci_r_pos | The position, in bp, of the right boundary of a 95% confidence interval around the peak marker |

**File S5 -- NILgenos.vcf**
A file in variant caller format containing whole-genome sequence data for each of the NILs (ECA230, ECA232, ECA411, and ECA528) mentioned in the manuscript.

**File S6 -- simpleNIL.csv**

| Column | Description |
|---|---|
| breaks | For each strain, the number of breakpoints between the N2 and CB4856 genotypes that are supported by lengths of at least 100 reads |
| cleanend | After cleaning for breakpoints supported by at least 100 reads, the genotype of a region of the genome, where 1 indicates the N2 genotype and 2 indicates the CB4856 genotype |
| chrom | Chromosome, as a roman numeral (or MtDNA for mitochondrial genome) |
| sample | Strain name (ECA230, ECA232, ECA411, or ECA528) |
| groupstart | After cleaning for breakpoints supported by at least 100 reads, the start position, in bp, of a given genotype block |
| groupend | After cleaning for breakpoints supported by at least 100 reads, the end position, in bp, of a given genotype block |

**File S7 -- allstats.csv**

| Column | Description |
|---|---|
| experiment | The name of the assay that was used for statistical comparison |
| condition | The condition used for statistical comparison, either raw_water (pruned, not control-regressed) or regressed_bleomycin (pruned and control-regressed |
| trait | The trait whose phenotype was used for the statistical comparison |
| comp | The strains being compared, separated by a hyphen. In the case of heterozygous animals, the two genotypes comprising the heterozygote are separated by an underscore and a hyphen separates the two strains being compared. |

| | |
|---|---|
| p adj | P-value statistic, adjusted for sample size, of a Tukey HSD test for an analysis of variance with the formula phenotype ~ strain |

**File S8 -- bleopheno_scantwo_summary.csv**

| **Column** | **Description** |
|---|---|
| chr1 | Chromosomal location of the first of the paired loci |
| chr2 | Chromosomal location of the second of the paired loci |
| pos1 | Location in cM of the first of the paired loci |
| pos2 | Location in cM of the second of the paired loci |
| lod.full | LOD score for the full model, evidence for at least one QTL |
| lod.fv1 | LOD score indicating the improvement of the full model over a single-QTL model (evidence for a second QTL) |
| lod.int | LOD score indicating the improvement of the full model over an additive model (evidence for interaction between loci) |
| lod.add | LOD score with a strictly additive model |
| lod.av1 | LOD score indicating the improvement of the additive model over a single-QTL model (evidence for second additive QTL) |

**File S9 -- region_elements.csv**

| **Column** | **Description** |
|---|---|
| biotype | Annotation for region element, either protein_coding, ncRNA, pseudogene, piRNA, miRNA, tRNA, lincRNA, or snoRNA |
| locus | Either the WormBase gene ID or the common gene name |
| gene_id | WormBase gene ID |
| sequence_name | Primary accession sequence ID |
| small_region | True/false indicating whether the element is within the *secb-1* QTL confidence interval (V:11,042,745-11,189,364) |

**File S10 -- exoninfo.csv**

| **Column** | **Description** |
|---|---|
| gene | Primary accession sequence ID |
| chr | Roman numeral indicating on which chromosome a gene is positioned |
| strand | Sense (+) or antisense (-) strand of DNA on which the gene is found |
| txstart | Position, in bp, of the transcription start site |
| txend | Position, in bp, of the transcription end site |
| codingstart | Position, in bp, of the coding sequence start |
| codingend | Position, in bp, of the coding sequence end |
| numexons | Number of exons within the gene |
| exonstarts | Positions, in bp, of the start of each exon, separated by commas |
| exonends | Positions, in bp, of the end of each exon, separated by commas |
| wbgene | WormBase gene ID |
| type | Indicates “coding” or “noncoding” sequence |

**File S11 -- snpeff.csv**

| **Column** | **Description** |
|---|---|
| CHROM | The chromosome, in roman numerals, on which the variant is located |
| POS | The position, in bp, at which the variant was identified |
| strain | The strain name of the sample used to identify the variant |
| REF | The reference allele at the variant site |

| | |
|---|---|
| ALT | The alternate allele at the variant site |
| GT | REF or ALT indicating which allele the particular strain has at the variant site |
| effect | Predicted effect of the variant |
| impact | Predicted level of impact of the variant on gene product function |
| gene_name | Well-known gene name that is impacted by a given variant |
| gene_id | WormBase gene ID of the gene impacted by a given variant |
| feature_id | Primary accession sequence ID of the element impacted by a given variant |
| exon_intron_rank | A fraction with the numerator indicating in which exon the variant was identified and the denominator indicating the total number of exons in the gene |
| nt_change | The position in the gene sequence at which the variant was identified and the nucleotide change in the format REF>ALT |
| aa_change | The predicted amino acid change introduced by the identified variant in the format REF ### ALT, where ### is the position of the amino acid affected by the variant |

**File S12-- expression_probes.csv**

| **Column** | **Description** |
|---|---|
| strain | Name of RIAIL for which gene expression was measured |
| trait | Probe name on the microarray |
| phenotype | Expression level of a given probe |

**File S13 -- expression_peaks.csv**

| Column | Description |
|---|---|
| marker | Genomic marker at which the correlation between genotype and probe expression was tested |
| chr | Chromosome on which the marker can be found |
| pos | Position, in bp, at which the genetic marker can be found |
| trait | Probe for which RIAIL expression was measured |
| lod | Log of odds ratio for correlation between genotype at the genetic marker and gene expression of RIAILs |
| threshold | Genome-wide error rate threshold for a particular iteration of the mapping, above which a LOD is considered significant |
| iteration | Numerical value indicating the mapping process iteration |
| var_exp | Amount of variation in gene expression across RIAILs that can be explained by genetic variation at the QTL |
| eff_size | Effect size of the QTL, calculated as the slope of a linear model with the formula (expression ~ genotype). A positive value indicates that RIAILs with the CB4856 allele have more gene expression than those with the N2 allele, and *vice versa* |
| ci_l_marker | The genetic marker indicating the left boundary of a 95% confidence interval around the peak marker |
| ci_l_pos | The position, in bp, of the left boundary of a 95% confidence interval around the peak marker |
| ci_r_marker | The genetic marker indicating the right boundary of a 95% confidence interval around the peak marker |
| ci_r_pos | The position, in bp, of the right boundary of a 95% confidence interval around the peak marker |

**File S14 -- annotated_expressionmap.csv**

| Column | Description |
|---|---|
| marker | Genomic marker at which the correlation between genotype and probe expression was tested |
| chr | Chromosome on which the marker can be found |
| pos | Position, in bp, at which the genetic marker can be found |

| | |
|---|---|
| trait | Probe for which RIAIL expression was measured (.A_12_P104350, for *H19N07.3*) |
| lod | Log of odds ratio for correlation between genotype at the genetic marker and gene expression of RIAILs |
| threshold | Genome-wide error rate threshold for a particular iteration of the mapping, above which a LOD is considered significant |
| iteration | Numerical value indicating the mapping process iteration |
| var_exp | If applicable (in the case of a significant QTL), amount of variation in gene expression across RIAILs that can be explained by genetic variation at the QTL |
| eff_size | If applicable (in the case of a significant QTL), effect size of the QTL, calculated as the slope of a linear model with the formula (expression ~ genotype). A positive value indicates that RIAILs with the CB4856 allele have more gene expression than those with the N2 allele, and *vice versa* |
| ci_l_marker | The genetic marker indicating the left boundary of a 95% confidence interval around the peak marker |
| ci_l_pos | The position, in bp, of the left boundary of a 95% confidence interval around the peak marker |
| ci_r_marker | The genetic marker indicating the right boundary of a 95% confidence interval around the peak marker |
| ci_r_pos | The position, in bp, of the right boundary of a 95% confidence interval around the peak marker |

**File S15 -- bleopheno_expression_corr.csv**

| **Column** | **Description** |
|---|---|
| strain | Name of strain in the RIAIL panel |
| bleopheno | Value indicating a strain’s residual median optical density in bleomycin |
| expression | Value indicating a strain’s expression level of *scb-1* |

**File S16 -- H19_RNAseq.csv**

| Column | Description |
|---|---|
| target_id | The gene isotype for which RNA seq data is reported |
| sample | Sample name |
| est_counts | Estimated counts of transcript in the sample |
| tpm | Number of transcripts of the given target per million transcripts |
| eff_len | Effective length of the transcript, calculated as (gene length - sequencing depth +1) |
| len | Length of transcript |
| condition | Genotype of the mixed-stage sample, either N2 or CB4856 |

**File S17 -- expression_scantwo_summary.csv**

| Column | Description |
|---|---|
| chr1 | Chromosomal location of the first of the paired loci |
| chr2 | Chromosomal location of the second of the paired loci |
| pos1 | Location in cM of the first of the paired loci |
| pos2 | Location in cM of the second of the paired loci |
| lod.full | LOD score for the full model, evidence for at least one QTL |
| lod.fv1 | LOD score indicating the improvement of the full model over a single-QTL model (evidence for a second QTL) |
| lod.int | LOD score indicating the improvement of the full model over an additive model (evidence for interaction between loci) |
| lod.add | LOD score with a strictly additive model |
| lod.av1 | LOD score indicating the improvement of the additive model over a single-QTL model (evidence for second additive QTL) |

**File S18 -- bleo_gwaspheno.csv**

| Column | Description |
|---|---|

| | |
|---|---|
| trait | Trait measured by the BIOSORT (bleomycin_median.EXT) |
| strain | Strain for which the trait was measured with the HTA fitness assay |
| phenotype | Residual phenotype value of a particular strain for the given trait |

**File S19 -- GWAprocessed.csv**

| **Column** | **Description** |
|---|---|
| marker | Name of a genomic marker tested for correlation between genotype and phenotype |
| CHROM | Chromosome, in roman numerals, on which the genomic marker resides |
| POS | Position, in bp, of the genomic marker |
| trait | HTA trait for which phenotypic values were measured and correlated to genomic markers |
| log10p | $\log_{10}(p)$, value of the correlation between genomic marker and phenotype |
| BF | Bonferroni-corrected p-value above which correlations are considered to be significant |
| aboveBF | Number indicated whether a marker reaches the BF significance level (0 = False, 1 = True) |
| startPOS | If applicable, the leftmost position of a particular peak confidence interval |
| peakPOS | If applicable, the position for a peak at which the log10p score is maximized |
| endPOS | If applicable, the rightmost position of a particular peak confidence interval |

**File S20 --allvars.csv**

| **Column** | **Description** |
|---|---|
| CHROM | Chromosome location of a given variant |

| | |
|---|---|
| POS | Position, in bp, of a given variant |
| REF | The reference (N2) allele at the variant site |
| ALT | The alternate allele at the variant site |
| AB1:WN2002 | Names of wild isolates tested with the HTA for bleomycin. -1 indicates a REF allele and 1 indicates an ALT allele at a given variant. |
| freq | If the minor allele frequency of a given variant within the strains tested is < 0.05, then “rare”, otherwise “common” |

**File S21 -- scb1FASTA.fa**
FASTA file of protein sequences for homologs of SCB-1

**File S22 -- scb1_tree.ph**
Parenthetical format of a neighbor-joining tree constructed from a multiple-sequence alignment of the SCB-1 homolog FASTA file
